# Malignant and stromal partial EMT programmes and cetuximab outcome in head and neck cancer

**DOI:** 10.64898/2026.10.09.757840

**Authors:** Dania Zuhier Ragheb Alkhatib, Sophia Lunetto, Priyanka Chakraborty, Mohit Kumar Jolly, Michael P. Philpott, Adrian Biddle, Gehad Youssef, Namshik Han

## Abstract

Partial epithelial-to-mesenchymal transition (pEMT) was defined in single malignant cells of head and neck squamous cell carcinoma but is usually scored in bulk tumours. In three single-cell datasets, its laminin-332 and hemidesmosome genes were expressed by malignant cells (the malignant arm) and its fibrillar collagens and vimentin by fibroblasts (the stromal arm). In bulk tumours the stromal arm dominated co-expression, so standard pEMT scores mainly measure stroma. We recovered the malignant arm with a classifier trained on single cells and a 12-gene core. In 40 patients treated with platinum and cetuximab, both malignant measures were higher in those with long progression-free survival (area under the curve 0.78 and 0.82), and the stromal arm was not. The malignant arm was inseparable from the Basal subtype, whose score did as well or better in these patients (0.93) and in two xenograft panels. Bulk pEMT scores should be read by cell of origin.

## Introduction

Head and neck squamous cell carcinoma (HNSCC), which arises in the oral cavity, pharynx and larynx, is among the most common cancers worldwide^1^. About 10% of patients have distant metastases at diagnosis and a further 20 to 30% develop them later, and median survival with metastatic disease is about 10 months^2^.

Cetuximab is the only drug targeting the epidermal growth factor receptor (EGFR) approved for HNSCC^3^. Since KEYNOTE-048^4^, first-line treatment of recurrent and metastatic disease has been based on pembrolizumab. Cetuximab with platinum remains an option for recurrent or metastatic tumours with low PD-L1 expression or when immunotherapy is unsuitable, and in locally advanced disease cetuximab is combined with radiotherapy in patients unfit for cisplatin. Biomarkers tested at scale have mainly measured the receptor itself. Neither EGFR protein in RTOG 0522 (380 tumours assessed)^5^ nor EGFR copy number in the EXTREME trial (n = 312)^6^ predicted treatment benefit. Two trials that replaced cisplatin with cetuximab in human papillomavirus (HPV)-positive oropharyngeal cancer, RTOG 1016^7^ and De-ESCALaTE^8^, found cetuximab inferior to cisplatin, and as a single agent cetuximab produces responses in about 13% of platinum-refractory patients^9^. By contrast, subcutaneous amivantamab, an EGFR–MET bispecific antibody, produced responses in 42% of 102 patients with HPV-unrelated disease after platinum and PD-1 blockade^10^ and is under FDA review for this indication (Johnson & Johnson press release, 30 July 2026, https://www.jnj.com/media-center/press-releases/johnson-johnsons-rybrevant-faspro-amivantamab-and-hyaluronidase-lpuj-receives-u-s-fda-priority-review-as-potential-first-in-class-egfr-and-met-targeted-subcutaneous-treatment-for-advanced-head-and-neck-cancer). No biomarker yet guides the use of either drug.

Tumour-cell plasticity is one candidate explanation for the variation in response between patients. In epithelial-to-mesenchymal transition (EMT), epithelial cells lose adhesion and gain mesenchymal traits, including migration, invasion and resistance to apoptosis. EMT has been linked to immune evasion^11^ and poorer survival across carcinomas^12^. EMT is a spectrum, and hybrid epithelial/mesenchymal (E/M) cells are more stem-like than cells at either pole and promote metastasis and therapy resistance across carcinomas^13^. In primary oral tumours disseminating cells with an EMT cancer stem cell profile predict metastasis^14^. These states are therefore candidate biomarkers and targets for treatment.

Puram and colleagues sequenced about 6,000 single cells from 18 oral cavity tumours and found that malignant cells varied in a few recurring expression programmes, among them cell cycle, stress, hypoxia, epithelial differentiation and partial EMT (pEMT)^15^. Malignant cells high in the pEMT programme kept their epithelial identity while expressing some mesenchymal and matrix genes and lay at the leading edge of the tumour, and the programme predicted nodal metastasis in bulk tumours. The programme was published as a list of its 100 top-ranked genes, the pEMT gene list. Puram and colleagues separated it from the fibroblast programme, although fibroblasts express many of the same genes. They also showed in The Cancer Genome Atlas (TCGA) that the Mesenchymal expression subtype reflects cancer-associated fibroblasts (CAFs). Once the expected contribution of non-malignant cells was regressed out of each bulk profile, Basal and Mesenchymal tumours merged into a single malignant-basal subtype, in which pEMT was highest^15^.

Computational scores quantify the EMT status of tumours for prognosis and treatment studies^16^. The 76-gene signature (76GS) score, derived in non-small cell lung cancer^17^, weights each gene by its correlation with CDH1 (E-cadherin), so that higher scores are more epithelial^16^. The multinomial logistic regression (MLR) score assigns each sample probabilities of the epithelial, hybrid and mesenchymal states from a model trained on the NCI-60 cell lines, with CLDN7 and the ratio of VIM to CDH1 as predictors^18^. The pEMT signature and its genes have been scored mostly in bulk transcriptomes, to stratify patients by survival^19,20^ and response to radiotherapy^20^, and in an immunohistochemical form to predict nodal metastasis in oral cavity cancer^21^. In bulk tissue, gene covariation largely follows cell-type proportions, so a signature whose genes are also expressed by abundant stroma reports stromal content. Bulk mesenchymal signatures are known to track fibroblast content^22^, a three-gene fibroblast signature reproduces the prognostic value attributed to EMT markers^23^, and the published EMT scoring metrics agree broadly but not in every dataset^16^. Puram and colleagues corrected whole profiles for composition, but it is not known whether the pEMT gene list itself can be divided gene by gene between the cells that express it, or whether the parts carry different clinical information.

Here we divide the pEMT list by the cell type that expresses each gene, show how each part behaves in bulk tumours, recover the malignant part with a classifier trained on single cells, and test both parts against outcome in patients and xenografts treated with cetuximab (Figure 1).

**Figure 1.**
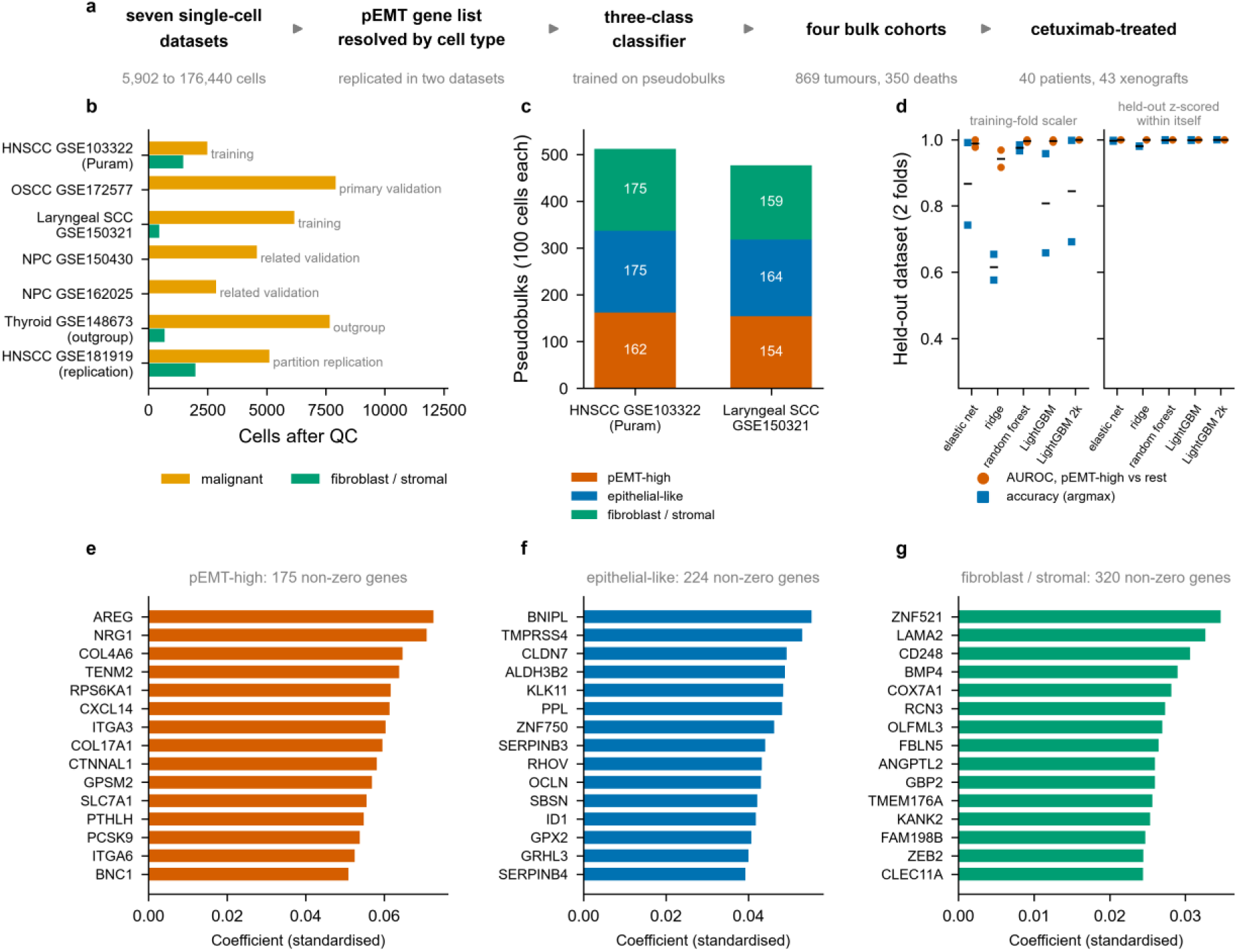
Study design and classifier construction. **a** The five stages of the analysis, from the single-cell datasets through the partition of the pEMT gene list by cell type to the cetuximab-treated patients and patient-derived xenografts. **b** Single-cell datasets used, with the number of malignant cells and fibroblasts each contributes and its role: training, validation, outgroup, or replication of the partition (GSE181919, tumour tissue only). **c** Composition of the labelled training set by class, in pseudobulk mixtures of 100 cells each. **d** Held-out performance under leave-one-dataset-out cross-validation for five learners fitted on identical folds, showing the area under the receiver operating characteristic curve for pEMT-high against the other two classes and the accuracy of the argmax class assignment, with the horizontal bar the mean across folds. In the left half the held-out dataset is standardised with the training fold’s scaler. In the right half it is standardised within itself, as every bulk cohort is, and the fitted logistic layer is then applied (tree models are refitted on the standardised training fold). **e**, **f**, **g** The largest positive elastic-net coefficients for the pEMT-high, epithelial-like and fibroblast or stromal classes, with the number of genes carrying a non-zero coefficient stated above each. Held-out discrimination reflects the label definition as well as the biology. The full learner comparison is in Supplementary Fig. 5.

Earlier corrections for stroma adjusted whole bulk profiles^15,22^ or excluded stroma-rich tumours^20^. Assigning each gene of the list to the cells that express it gives two gene sets that can be scored in every bulk tumour, including archived cohorts without single-cell data, and shows which part of the programme is associated with outcome under cetuximab.

## Results

### Expression of the pEMT genes ranges from mainly malignant to mainly stromal

A bulk pEMT score sums the expression of its genes regardless of which cells express them, so we first traced each gene to its source. Of the 100 pEMT genes, only vimentin is a canonical mesenchymal marker, the EMT transcription factors and N-cadherin are absent, and many of the laminins, integrins, collagens and other matrix genes are also expressed by fibroblasts. In the dataset in which the list was defined, the 5,902 cells of Puram et al. (the discovery dataset), we computed, from counts per 10,000, the share of each gene’s expression contributed by malignant cells and by fibroblasts (Figure 2a–c, Supplementary Data 1). The five laminin-332 and hemidesmosome genes in the list were the most malignant-restricted. COL17A1 drew 99.3% of its expression from malignant cells, LAMB3 99.1%, LAMC2 98.3% and LAMA3 96.0%, a family median of 98% against 42% for the other genes (one-sided Mann-Whitney p = 4.6 × 10⁻⁶). At the other end, COL1A1 drew 96% of its expression from fibroblasts, TAGLN 98%, MMP2 88% and vimentin 66%. Genes at both ends were expressed more highly in pEMT-high malignant cells (the top quarter by Puram pEMT score) than in epithelial-like cells, which score high for epithelial differentiation (Figure 2c).

**Figure 2.**
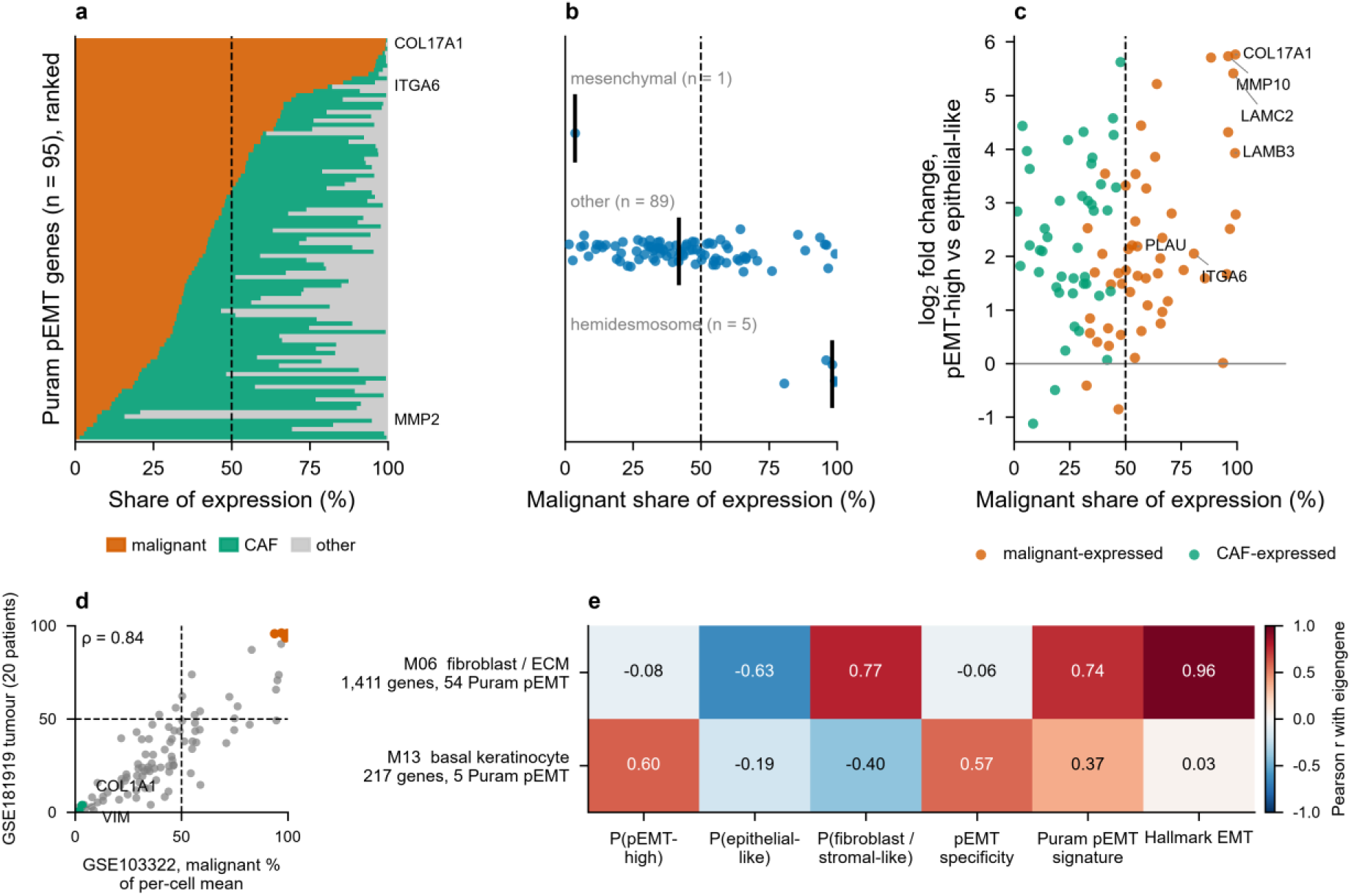
Expression of the pEMT genes ranges from mainly malignant to mainly stromal. **a** Each of the 95 detected Puram pEMT genes, ranked by the share of its total expression (linear scale, counts per 10,000) contributed by malignant cells across all 5,902 cells of GSE103322. Orange is the malignant share, green the fibroblast and myofibroblast share, grey all other cell types. The dashed line marks equal contribution. Selected genes are named in the right margin. **b** Malignant share by gene family, one point per gene, with the vertical line marking the median. Families are restricted to genes in the pEMT list, so the hemidesmosome and laminin-332 family has five members (COL17A1, ITGA6, LAMA3, LAMB3 and LAMC2) and the classical mesenchymal family one (vimentin). **c** Log2 fold change between pEMT-high and epithelial-like malignant cells (vertical axis) against malignant share (horizontal axis), coloured by whether total share assigns each gene to the malignant or the stromal side. Genes above zero are induced in pEMT-high cells, and induced genes lie at both ends of the horizontal axis. **d** Replication with a composition-independent measure: the malignant share of mean expression per cell for each pEMT gene in GSE103322 against the tumour tissue of GSE181919 (20 patients), with hemidesmosome and laminin-332 genes in orange and COL1A1 and vimentin in green. Dashed lines mark equal malignant and fibroblast shares. ρ is the Spearman correlation coefficient across the 94 genes detected in both datasets (p = 1.1 × 10⁻²⁶). The other two dataset pairs are in Supplementary Data 2c. **e** Pearson correlation between the eigengenes of the two co-expression modules discussed in the text and the classifier outputs and signature scores, across 520 TCGA-HNSC primary tumours. M06 is the fibroblast and extracellular-matrix module that carries most of the Puram pEMT genes. M13 is the basal keratinocyte module that tracks pEMT specificity. Module size and the number of Puram pEMT genes each contains are given with the row label. The full thirteen-module matrix is in Supplementary Fig. 2.

By share of total expression, which is weighted by cell numbers, 52 of the 95 detected genes are malignant-predominant, and by mean expression per cell 33 are. We repeated the per-cell measure in the two other datasets with annotated fibroblasts, laryngeal GSE150321 (two patients)^24^ and GSE181919 (tumour tissue from 20 patients)^25^, where 23 and 20 genes were malignant-predominant. The ordering of the 94 genes detected in all three datasets replicated, with Spearman correlations of 0.84 between the discovery data and GSE181919 (p = 1.1 × 10⁻²⁶, Figure 2d), 0.88 between the discovery data and GSE150321 (p = 6.2 × 10⁻³²) and 0.91 between GSE150321 and GSE181919 (p = 4.5 × 10⁻³⁶, Supplementary Data 2 to 2d). The hemidesmosome family had a median per-cell malignant share of 99%, 92% and 96% in the three datasets. In each of the 13 GSE181919 patients with at least 20 malignant cells and 20 fibroblasts, COL17A1, the laminin-332 chains and ITGA6 were malignant-predominant, and COL1A1, COL5A2, MMP2, TAGLN and vimentin were fibroblast-predominant. We therefore treat the list as a gradient with two stable ends, the malignant and stromal arms. For scoring, we took two small gene sets from the ends of the gradient: the malignant core, the 12 genes with at least 80% malignant share in the discovery data, and the stromal core, the 14 genes with at least 60% fibroblast share (Supplementary Data 3). Throughout, “core” refers only to these two gene sets. Ten of the 12 malignant-core genes were malignant-predominant in all three datasets, and MMP10 and CXCL14 were at parity, with similar malignant and fibroblast shares, in GSE181919. All 14 stromal-core genes were fibroblast-predominant in all three. The two cores give each arm a gene set that holds across datasets and patients, so they can be used to ask which arm a bulk score reports.

### In bulk tissue the stromal arm dominates pEMT co-expression

Whole tumours contain both arms, so we tested which one dominates their co-expression. In a signed co-expression network of the 520 TCGA-HNSC primaries (Supplementary Fig. 1, Supplementary Data 4), 54 of the 76 pEMT genes present fell into one 1,411-gene module, M06 (hypergeometric p = 1.1 × 10⁻²⁴). The M06 eigengene, its first principal component, correlated with Hallmark EMT at r = +0.96 and with the stromal core at ρ = +0.91 (Supplementary Data 3b), and its most connected genes included FBN1, VCAN, PDGFRB and COL6A3 (Figure 2e, Supplementary Fig. 2). In bulk HNSCC the canonical Puram pEMT signature therefore measures the fibroblast and extracellular-matrix compartment, even though malignant cells also express many of its genes.

### A classifier trained on single-cell pseudobulks recovers the malignant arm

Bulk co-expression cannot isolate the malignant arm, so we trained a three-class elastic-net classifier (pEMT-high malignant, epithelial-like malignant, fibroblast) on pseudobulks, synthetic bulk-like profiles pooled from labelled cells of two HNSCC datasets (Figure 1a–c). The labels are thresholds on Puram scores whose genes are also features, so held-out discrimination (area under the curve 0.978 to 1.000 with the training fold’s scaler and 1.000 with each held-out dataset standardised within itself, as bulk cohorts are, Figure 1d) reflects the label definition and does not by itself validate the classifier. Figure 1e–g shows the largest coefficients for each class. In the two training datasets every malignant and fibroblast pseudobulk was assigned to its own class. In the four datasets not used for training the separation was incomplete (Figure 3a, b, Supplementary Figs. 3 and 4, Supplementary Data 5). Every fibroblast pseudobulk was called fibroblast, but so were 5 of the 26 malignant pseudobulks. Most nasopharyngeal malignant pseudobulks carried a fibroblast probability of 10% to 35%, and one malignant pseudobulk of the thyroid outgroup reached a pEMT-high probability of 0.85. The four other classifiers gave probabilities close to the elastic net’s (Spearman 0.92 to 0.96, Supplementary Fig. 5, Supplementary Data 6). We summarise the output as pEMT specificity, the pEMT-high probability minus the larger of the epithelial-like and fibroblast probabilities (Supplementary Data 7). Because this difference is bounded, 15% of TCGA primaries fell below −0.9. A log-odds summary of the same probabilities removed this accumulation, correlated with pEMT specificity at 0.97 to 0.995 and gave the same conclusions, except that its pooled survival interval just included 1 (0.996 to 1.43) and its pooled xenograft odds ratio excluded 1 (2.58, 1.12 to 5.94, Supplementary Data 8).

**Figure 3.**
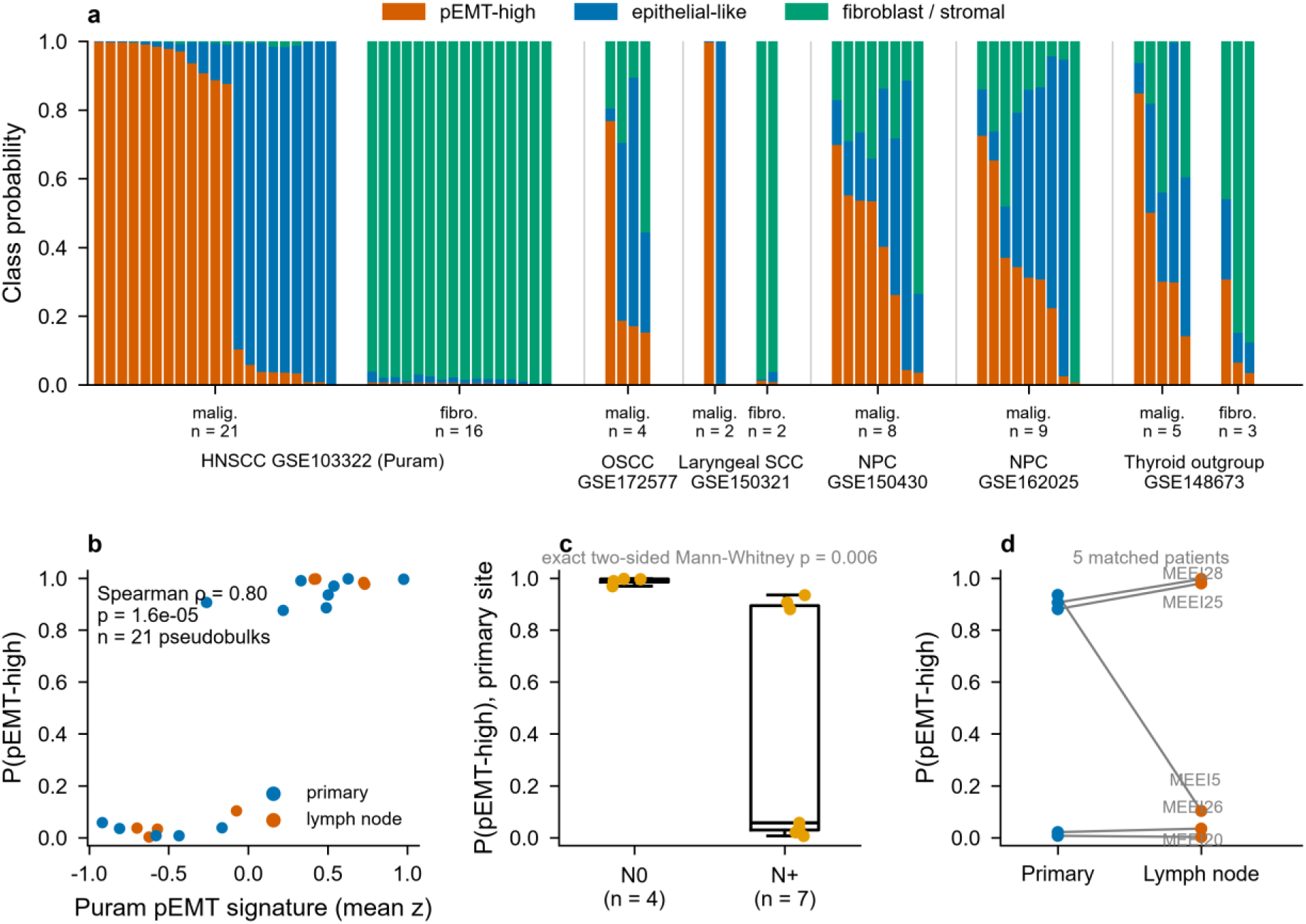
The classifier in the six single-cell datasets. GSE103322 and GSE150321 supplied the training cells. GSE172577, GSE150430, GSE162025 and GSE148673 were not used for training. **a** Classifier probabilities for malignant and fibroblast pseudobulks across all six datasets, one profile per patient and anatomical site, requiring at least 20 cells per profile. **b** Agreement between the classifier ranking and the Puram pEMT signature ranking within malignant pseudobulks. **c** P(pEMT-high) of the primary-site malignant pseudobulk in GSE103322 patients with node-negative and node-positive disease, with the exact two-sided Mann-Whitney p-value. All 11 patients are in the training data. **d** P(pEMT-high) in matched primary and lymph-node samples from the same patients, joined by grey lines. Boxes show the median and interquartile range and whiskers extend to 1.5 times the interquartile range.

**Figure 4.**
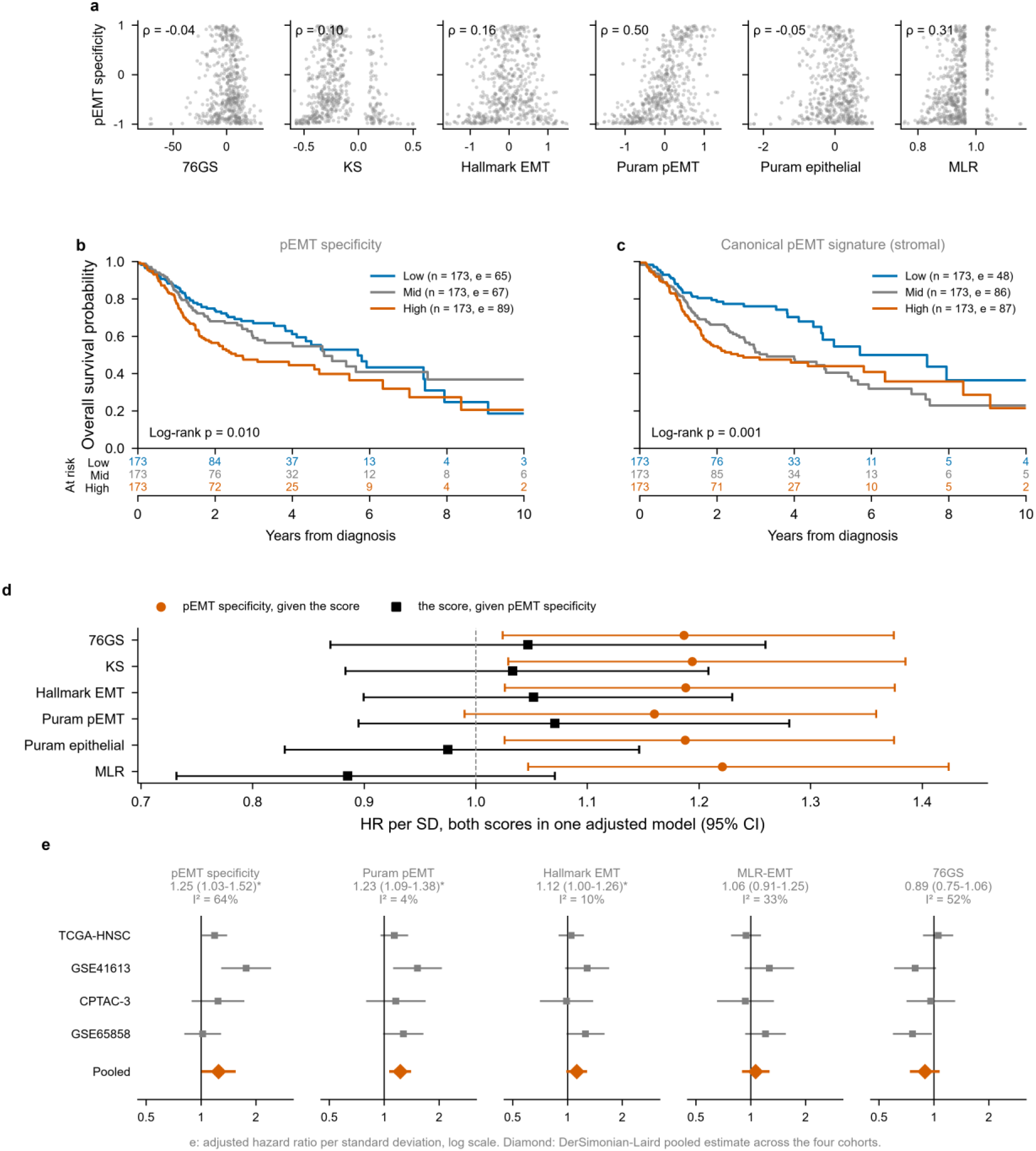
pEMT specificity against published EMT scores and its association with overall survival. **a** pEMT specificity plotted against each published score in TCGA-HNSC primaries, one point per tumour, with the Spearman correlation stated on each panel. MLR-EMT is implemented as published by George et al. **b** Kaplan-Meier curves for tertiles of pEMT specificity in TCGA-HNSC primaries, with numbers at risk beneath and a log-rank test. **c** The same analysis for the canonical pEMT signature, which reflects the stromal programme. **d** Head-to-head models in which pEMT specificity and one published score are entered together, showing the hazard ratio of each given the other. **e** Random-effects meta-analysis of the adjusted hazard ratio per standard deviation for five scores across TCGA-HNSC, GSE41613, CPTAC-3 and GSE65858, on a logarithmic axis. Grey squares are the per-cohort estimates with their 95% confidence intervals and the orange diamond is the DerSimonian-Laird pooled estimate. Hartung-Knapp intervals, which are wider when heterogeneity is present, are in Table 2 and Supplementary Data 39b. The pooled estimate and the I² statistic are stated above each panel. An asterisk marks a pooled p-value below 0.05. Vertical black lines mark a hazard ratio of 1. The single-cohort TCGA adjusted hazard ratio per score is in Supplementary Fig. 11 together with the TCGA-only adjusted Cox model.

Three results show that pEMT specificity does not simply reproduce its training labels. First, it tracks M13 (r = +0.57), a 217-gene basal keratinocyte module of the same TCGA co-expression network, and not M06 (r = −0.06). Second, the pEMT-high state can be recovered without any Puram programme gene. Trained on the 17,708 genes left after removing all 533 of them, a classifier separated the states in held-out patients (area under the curve 0.925, Supplementary Fig. 6, Supplementary Data 9). Third, the 10 pEMT genes among the classifier’s 100 largest pEMT-high coefficients are all malignant-predominant in the discovery data, six of them in all three datasets, and none is in the stromal core. pEMT specificity therefore reports the malignant arm, which the next sections test against other approaches and known composition.

### The partition differs from deconvolution of bulk EMT signatures

A related division has been made for EMT genes more broadly. Starting from the Hallmark and Tan EMT genes and genes correlated with SNAI1, SNAI2, TWIST1, VIM, ZEB1 and ZEB2, Tyler and Tirosh deconvolved bulk expression from TCGA into cancer-cell and fibroblast programmes and annotated them with single-cell data that included the Puram cohort^22^. Of those six markers, only vimentin is in the pEMT list, and 40 of the 100 pEMT genes fall in their gene set. In 520 TCGA primary tumours, the Tyler and Tirosh fibroblast compartment correlated with Hallmark EMT at +0.92 and their cancer compartment with pEMT specificity at 0.00 (Supplementary Fig. 7, Supplementary Data 10 and 11). Their approach removes fibroblast signal from a mesenchymal programme, whereas ours divides a largely non-mesenchymal programme between the cells that express it, so decoupling mesenchymal signatures does not recover the malignant arm.

The cancer-cell meta-programmes of Gavish and colleagues^26^ align with the two arms. EMT-II contains 7 of the 12 malignant-core genes (hypergeometric p = 1 × 10⁻¹⁶) and correlated with the malignant core at +0.84 and with pEMT specificity at +0.62 in TCGA, and EMT-I contains 10 of the 14 stromal-core genes and correlated with the stromal core at +0.96 (Supplementary Data 12). In GSE65021 EMT-II tracked progression-free survival (area under the curve 0.72) but added nothing to the malignant core, and of all 41 meta-programmes only epithelial senescence and a pancreatic-cancer programme passed a false discovery rate of 0.05, a post hoc result.

Two partial EMT gene sets defined independently of this study divide the same way (Supplementary Data 13 to 13e). The malignant partial EMT programme that Simkin and colleagues derived from spatial transcriptomes of 26 HNSCC^27^ shares 7 of the 12 malignant-core genes. It drew a median 67% of its expression from malignant cells in the discovery data and correlated with the malignant core at +0.93 in TCGA, and their fibroblast programme correlated with the stromal core at +0.93. The partial EMT cluster of a mammary EMT signature^28^ shares no gene with the malignant core, yet in HNSCC it was also expressed by malignant cells (median share 81%), whereas its full EMT cluster drew a median 15% from malignant cells and correlated with the stromal core at +0.90. In breast cancer the basal subtype tracks partial more closely than full EMT^29^. In TCGA-HNSC the Basal centroid score correlated with the mammary partial EMT cluster at +0.37 and with the full EMT cluster at −0.35 (Steiger p = 5 × 10⁻⁵⁵), and with the Simkin partial EMT and fibroblast programmes at +0.49 and −0.24. In GSE65021 both partial EMT sets separated long from short progression-free survival (area under the curve 0.86 for each, p = 2 × 10⁻⁴), and the full EMT cluster and the fibroblast programme did not (0.35). Each partial EMT set correlated with the Basal score at 0.84 and kept no association beside it in a joint model. The division into a malignant and a stromal arm, and the link of the malignant arm to the Basal subtype, therefore hold for partial EMT gene sets defined in other data and in another tissue.

### Scored from its core gene set, the malignant arm tracks pEMT specificity

Each arm can also be scored without the classifier, as the mean z-score of the genes in its core set. In TCGA the malignant core correlated with pEMT specificity at +0.67 and with M13 at +0.59 (Supplementary Data 3 and 3b). The broader 52-gene malignant arm defined by total share correlated at +0.97 with the full Puram signature, because its genes near parity carry fibroblast signal in bulk tissue. pEMT specificity and the malignant core are therefore the primary measures of the malignant arm.

### In simulated tumours the malignant measures do not rise with stroma

To test the measures against known composition, we built 450 synthetic tumours from GSE181919 cells, which the classifier never saw. We varied the share of pEMT-high malignant cells, the fibroblast fraction (5% to 60%) and the immune fraction (5% to 35%) independently (Methods, Supplementary Data 14 and 14b). The canonical Puram signature rose with the malignant state (standardised coefficient 0.68) and almost as much with the fibroblast fraction (0.62), and Hallmark EMT followed fibroblasts almost entirely (0.95). The malignant core followed the malignant state (0.79) and fell as fibroblasts (−0.39) and immune cells (−0.31) diluted the tumour. pEMT specificity behaved likewise (0.66 for the malignant state and −0.46 for fibroblasts). Neither malignant measure rises with stroma, but both partly track tumour purity. BayesPrism deconvolution^30^, with GSE181919 cells as the reference, gave an independent check in TCGA and GSE65021. The fibroblast fraction correlated with the stromal core at 0.89 and 0.90, and the malignant core scored on deconvolved malignant-cell expression correlated with the bulk core at 0.95 and 0.94 (Supplementary Data 15). In TCGA the deconvolved malignant-cell pEMT score was highest in the Mesenchymal and Basal subtypes, the two that Puram and colleagues merged as malignant-basal, but it also rose with the fibroblast fraction (0.40), so on RNA-seq the deconvolution did not fully separate the arms. In GSE65021 it was independent of the fibroblast fraction (0.04). Both primary measures therefore report the malignant state, but purity has to be accounted for when they are tested against outcome.

### pEMT specificity is largely distinct from published EMT scores

Existing EMT scores might already capture pEMT specificity. We therefore computed 76GS (a 76-gene score that is high in epithelial tumours), the Kolmogorov-Smirnov (KS) statistic, Hallmark EMT, the Puram programmes and MLR-EMT (the ordinal model of George et al.^18^) on the same tumours. The published scores share one epithelial-to-mesenchymal axis, with KS correlating with Hallmark EMT at 0.77 and with MLR-EMT at 0.63. pEMT specificity correlated with them at −0.04 to 0.31 and with the Puram pEMT signature at 0.50 (Figure 4a, Supplementary Data 16). MLR-EMT was designed for partial EMT, which it calls the hybrid state between epithelial and mesenchymal. Implemented as published (Methods), it called every GSE65858 tumour and 88 of 97 GSE41613 tumours hybrid (Supplementary Data 17). On RNA-seq the state calls depend on how values are mapped to the microarray scale, with the ranking unchanged (Methods). The hybrid-state probability correlated with pEMT specificity at 0.35 in TCGA (Supplementary Data 18). pEMT specificity therefore carries information that the published EMT scores do not.

### pEMT specificity reflects a hemidesmosome and EGFR-ligand programme

We grouped genes by family to find which drive pEMT specificity in bulk tumours. In TCGA, eight hemidesmosome and laminin-332 genes, the five in the pEMT list plus ITGB4, ITGA3 and PLEC, correlated with pEMT specificity at a median of +0.65, EGFR ligands and the receptor at +0.40, basal keratinocyte genes at +0.43, and mesenchymal and fibroblast genes near zero (Figure 5a, Supplementary Data 19). The genes with the classifier’s 100 largest coefficients overlap far beyond chance with three HNSCC programmes derived by other routes (Table 1, Supplementary Data 19b). These are an EGFR-induced EMT signature from EGF-stimulated cell lines^31^, a set of EGFR-mediated invasion genes including nine that each separated long from short progression-free survival on cetuximab^32^, and a tumour-budding signature from spatial transcriptomics^33^. Five of the nine cetuximab-predictor genes (COL17A1, ITGA3, ITGB4, LAMA3 and LAMC2) are among the top 100 coefficients, where 0.04 would be expected by chance. pEMT specificity correlated with the nine cetuximab-predictor genes at +0.68 to +0.78 across four bulk cohorts, whereas 76GS never exceeded +0.17 against any published set (Figure 5b, c, Supplementary Data 19c). The canonical Puram signature correlated more strongly with the invasion and tumour-budding sets in TCGA (+0.82 to +0.88, against +0.55 to +0.71 for pEMT specificity) and similarly with the cetuximab-predictor set (+0.74 and +0.78, Figure 5b), so these correlations separate pEMT specificity from epithelial scores without favouring it over the canonical signature. The EGFR ligands came from the malignant cells themselves, which supplied 94% of TGFA and NRG1 expression, 84% of EGF and 75% of EREG, and the fibroblast-to-malignant signalling most specific to pEMT-high cells ran from collagens, fibronectin and thrombospondins to integrin α3β1 (Supplementary Note 1, Supplementary Fig. 8, Supplementary Data 20). The programme behind pEMT specificity is therefore adhesion with autocrine EGFR ligands in a basal keratinocyte context, which points to the Basal subtype.

**Figure 5.**
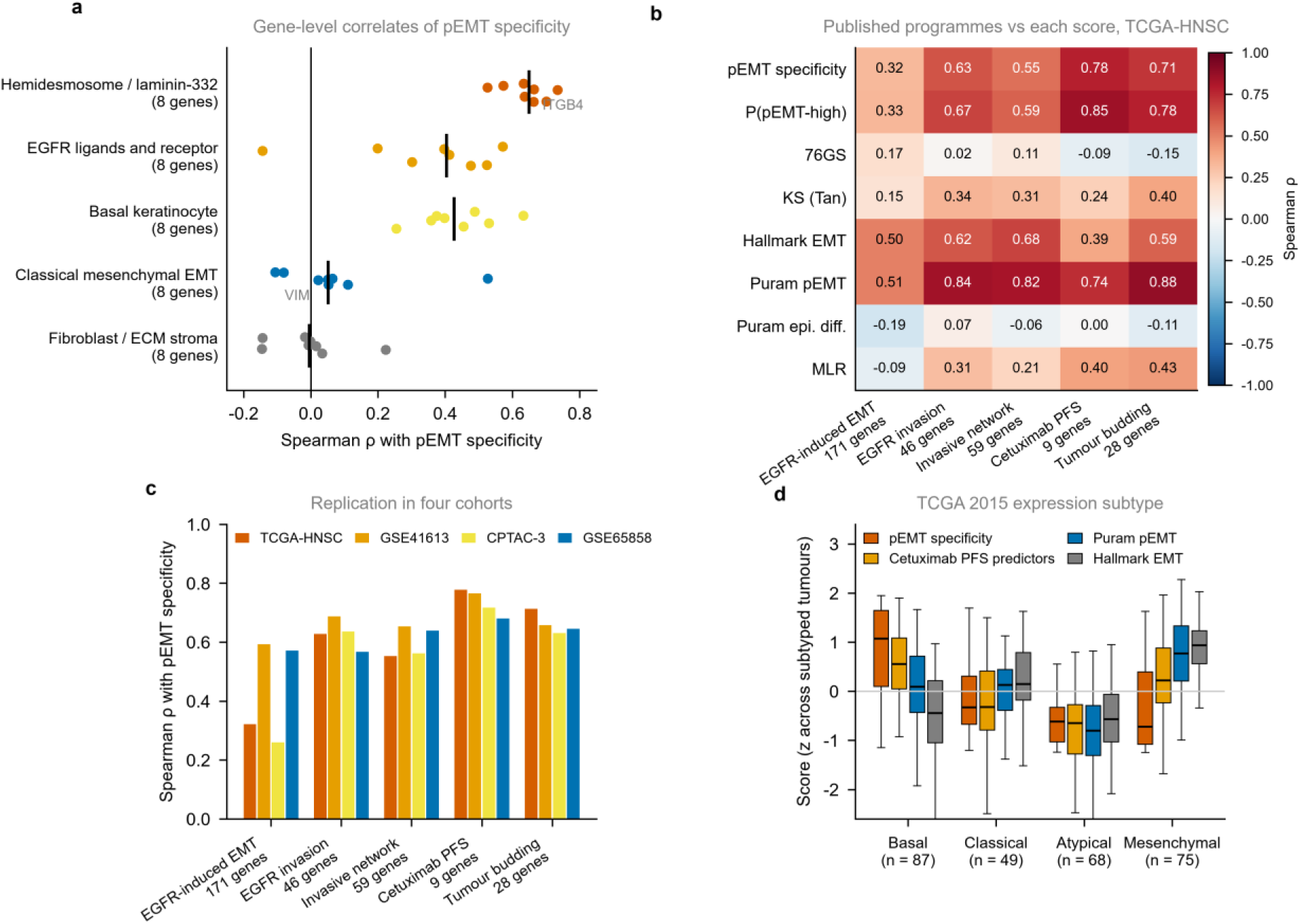
Independent validation of the scores against published programmes and molecular subtypes. **a** Spearman correlation between pEMT specificity and individual genes, grouped into five classes of eight genes, one point per gene, with the black vertical line marking the class median. The hemidesmosome and laminin-332 class adds ITGB4, ITGA3 and PLEC, which are not in the pEMT list, to the five pEMT genes of Figure 2b. **b** Spearman correlation between each published gene programme and each score, in TCGA-HNSC primaries. Red indicates positive correlation and blue negative, as shown by the colour bar. **c** Correlation between pEMT specificity and each published programme in all four cohorts. **d** Score by TCGA 2015 four-class expression subtype in the 279 matched primary tumours, for pEMT specificity, the published cetuximab progression-free-survival predictors, the Puram pEMT signature and Hallmark EMT. Each score is standardised across the subtyped tumours. Boxes show the median and interquartile range and whiskers extend to 1.5 times the interquartile range.

**Table 1.**
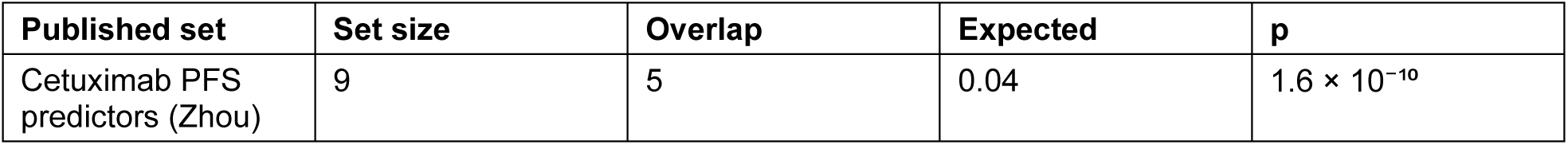
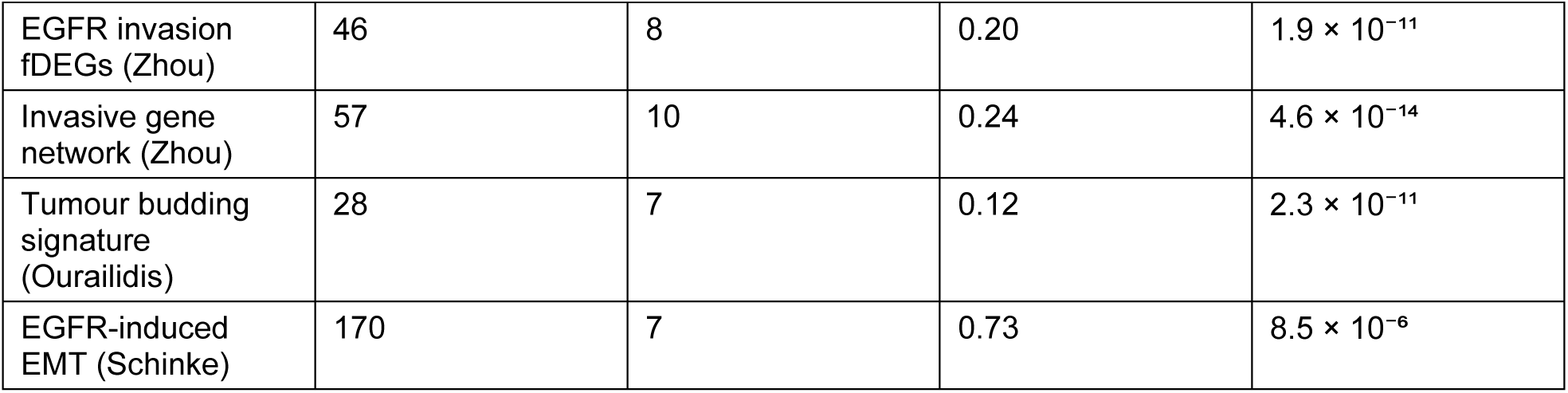
Overlap of the classifier’s top-weighted genes with independently published gene sets. Hypergeometric test of the genes with the 100 largest positive pEMT-high classifier coefficients against each published gene set, over the 23,436-gene classifier universe. Set size is the number of genes of that set present in the universe. Expected is the overlap expected by chance. The classifier was trained on single-cell pseudobulk labels alone and had no access to the data from which any of these sets was derived.

### pEMT specificity marks the Basal subtype

Because hemidesmosome genes and EGFR ligands mark basal keratinocytes, we compared the scores across the TCGA expression subtypes. In 279 TCGA primaries with published subtype calls^34^, pEMT specificity was highest in Basal tumours (Kruskal-Wallis p = 2.8 × 10⁻¹⁸), whereas KS, Hallmark EMT and the Puram signature peaked in the Mesenchymal subtype (Figure 5d). This agrees with the malignant-basal subtype of Puram et al., in which pEMT was highest once non-malignant signal was removed^15^. pEMT specificity places the programme in the Basal subtype, as their corrected analysis did, but because it subtracts the fibroblast probability it cannot test whether Mesenchymal tumours are malignant-basal tumours with more stroma. In patient-derived xenografts, 13 of 15 basal models responded to cetuximab, compared with 1 of 5 mesenchymal models^35^. GSE65858 comes with its own cluster labels under the same names, and these do not match the Walter centroid calls^36^ used in TCGA (Supplementary Data 21), so the subtype result rests on TCGA.

### Under platinum and cetuximab, only the malignant arm tracks progression-free survival

The EGFR-ligand programme of Basal tumours suggests dependence on EGFR, so both arms were tested against outcome under cetuximab. GSE65021^37^ contains 40 patients with recurrent or metastatic HNSCC treated with first-line platinum plus cetuximab. The deposited cohort consists of two outcome extremes balanced on the main clinical factors, 14 patients with long progression-free survival (beyond 12 months) and 26 with short progression-free survival (below 5.6 months). The classifier was applied without refitting (Supplementary Data 22 and 23), and all tests are two-sided.

Both malignant measures were higher in patients with long progression-free survival. pEMT specificity had a median of +0.27 against −0.50 (p = 0.004, Figure 6a), an area under the curve of 0.78 (95% confidence interval 0.62 to 0.92, Figure 6b) and a Firth-penalised odds ratio of 2.85 per standard deviation (1.40 to 6.73). With 14 events, the odds ratio adjusted for seven clinical covariates (3.48, 1.46 to 11.5) is reported only as a sensitivity analysis (Supplementary Data 24). The malignant core gave an area under the curve of 0.82 (0.68 to 0.94) and an odds ratio of 4.22 (1.61 to 15.4, p = 0.001), and the stromal core 0.42 (0.24 to 0.61) and 0.94 (Figure 6c, Supplementary Data 3c and 3d).

**Figure 6.**
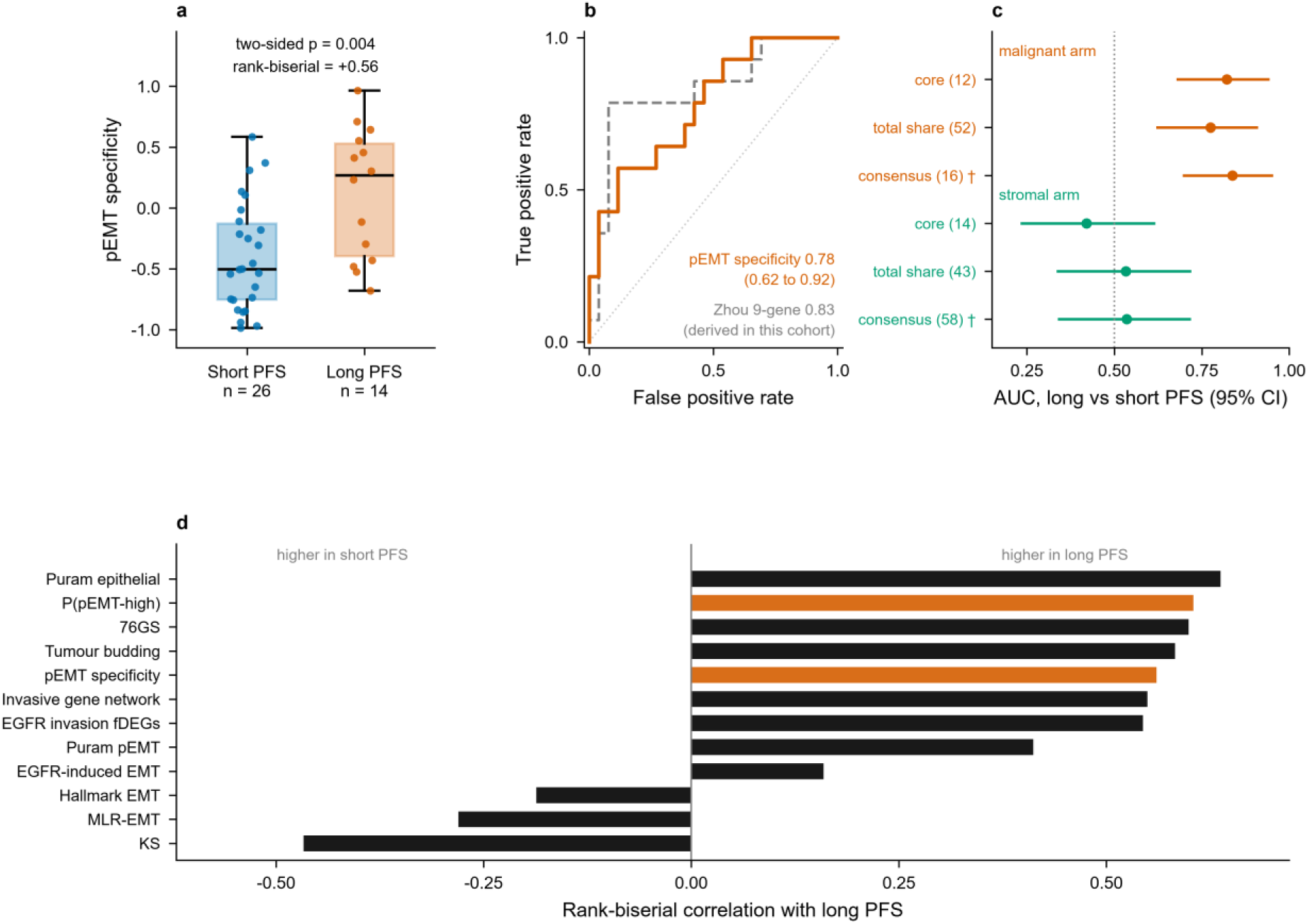
The malignant and stromal arms of the pEMT gene list and progression-free survival on platinum plus cetuximab in GSE65021 (exploratory). **a** pEMT specificity in the 26 patients with progression-free survival under 5.6 months and the 14 exceeding 12 months. Boxes show the median and interquartile range, whiskers extend to 1.5 times the interquartile range, and each point is one patient. The two-sided Mann-Whitney p-value and the rank-biserial correlation are stated. **b** Receiver operating characteristic curves for prolonged progression-free survival. pEMT specificity is shown in orange with its bootstrap confidence interval. The Zhou nine-gene set is shown in grey and dashed because it was derived on this cohort and serves as a positive control. **c** Scores computed directly from the genes of each arm: area under the curve with 2,000-resample bootstrap 95% confidence interval for three definitions of the malignant arm (orange) and of the stromal arm (green), with the number of genes in brackets. Core (primary definition), at least 80% malignant or 60% fibroblast share of total expression in the discovery data. Total share, the full partition. Consensus, genes assigned to the same arm by mean expression per cell in all three single-cell datasets. † Added after the first two definitions had been examined in this cohort. **d** Rank-biserial correlation with prolonged progression-free survival for every independently derived score, with positive values indicating higher scores in the long progression-free survival group. The classifier outputs are in orange. MLR-EMT is implemented as published by George et al.

Three sensitivity analyses added after these results agreed with them. Malignant-core thresholds from 60% to 90% gave areas under the curve of 0.78 to 0.81. Removing the three core genes that Zhou and colleagues selected in this cohort left 0.82, and the consensus arms (Methods) gave 0.84 for the malignant arm and 0.54 for the stromal arm. With 16 scores treated as one family, the false discovery rate was 0.008 for the malignant core and 0.009 for pEMT specificity (Supplementary Data 25). Standardising each patient against the other 39 left both areas under the curve unchanged. Scoring each patient alone against an external cohort on the same array family left the malignant core at 0.81, and pEMT specificity was less stable (Supplementary Data 26). Both measures partly track purity, but with the stromal core and an 18-gene immune score in the model the odds ratios stayed at 4.43 (1.59 to 18.1, p = 0.003) for the malignant core and 2.64 (1.27 to 6.32, p = 0.008) for pEMT specificity (Supplementary Data 27).

The strongest single scores in this cohort were two epithelial scores, Puram epithelial differentiation (0.82) and 76GS (0.80). Neither primary measure outperformed them or the Zhou nine-gene set derived on these patients (0.83). The paired difference from 76GS was +0.02 (−0.19 to 0.24) for the malignant core (Supplementary Data 28). The core did outperform the canonical pEMT signature, by 0.12 (0.02 to 0.23), and kept its association when the two were entered together (Firth odds ratio 8.90, 1.66 to 81.1, p = 0.007), as did pEMT specificity (2.56, 1.11 to 6.68, p = 0.027), although the collinearity makes these estimates imprecise (Supplementary Data 24b). The core correlated with 76GS at +0.33, and both remained associated when entered together, so the core is not a proxy for epithelial differentiation (Supplementary Data 3d and 24c). A post hoc combination of pEMT specificity with epithelial differentiation reached 0.91 and is shown only as exploratory (Supplementary Fig. 9, Supplementary Data 29).

MLR-EMT was not associated with outcome (Figure 6d). Its state calls, 23 hybrid and 17 mesenchymal, were not associated with outcome (Fisher p = 0.09). Its hybrid probability gave an area under the curve of 0.64 (p = 0.15), was uncorrelated with pEMT specificity (−0.06) and the malignant core (−0.01), and correlated with 76GS (+0.68), so in this cohort it measured how much of the epithelial state a tumour retained (Supplementary Data 18). KS was lower in patients with long progression-free survival (area under the curve 0.27, p = 0.029) and carried no information once epithelial differentiation was in the model (Supplementary Data 24d). pEMT specificity was collinear with the published EGFR programmes and added nothing to the Zhou gene sets or the tumour-budding signature in joint models (p = 0.10 to 0.62), although it remained associated alongside the Schinke signature (p = 0.008, Supplementary Data 24c).

On deconvolved malignant-cell expression, the canonical pEMT genes separated long from short progression-free survival (area under the curve 0.80, 0.65 to 0.92, p = 0.002), as did the malignant core (0.74, p = 0.012). The deconvolved fibroblast stromal core did not (0.57), and the fibroblast fraction was lower in patients with long progression-free survival (0.26, p = 0.012) (Supplementary Data 15b).

Because pEMT specificity is highest in Basal tumours, we asked whether the association reflects the subtype itself. With the centroids behind the published TCGA calls, 12 GSE65021 patients were called Basal, 11 Mesenchymal, 8 Atypical and 9 Classical, and 10 of the 12 Basal patients had long progression-free survival, against 4 of the other 28 (Fisher p = 6.0 × 10⁻⁵). The Basal centroid score gave an area under the curve of 0.93 (0.83 to 1.00), exceeded the malignant core by 0.11 (0.02 to 0.23) and correlated with it at 0.80, and removing the pEMT and Zhou genes from the centroids left 0.94. Entered together with the Basal score, neither the malignant core (Firth odds ratio 0.51, 0.14 to 2.45) nor pEMT specificity (0.64, 0.12 to 2.49) kept its association, but with 14 events and a correlation of 0.80 a joint model cannot apportion the association between them (Supplementary Data 30). Both scores could be computed one patient at a time. Scored against an external cohort on the same array family, or with no reference at all, the Basal score kept an area under the curve of 0.93 and the malignant core 0.81 to 0.83 (Supplementary Data 31). In this cohort the malignant-arm association therefore cannot be separated from the Basal subtype, which discriminated better, so we repeated the comparison in the xenograft panels below.

### Malignant core and Basal scores are higher in responsive xenografts

No second public patient cohort pairs baseline expression with cetuximab outcome. Two public HNSCC xenograft panels do, GSE84713 (26 patients, responder labels dichotomised from tumour volume)^35^ and GSE183881 (17 tested models, sensitive or resistant)^38^. Xenograft stroma is murine, so human signal from stromal genes comes from tumour cells or from mouse transcripts measured as human (GSE183881 discarded reads classified as mouse, but mouse transcripts can cross-hybridise on the GSE84713 microarrays), and the two arms cannot be separated in these panels. Both panels were scored without refitting (Supplementary Data 32 to 32c).

The malignant core was higher in responders in both panels, with areas under the curve of 0.75 (0.50 to 0.95, p = 0.041) and 0.73 (p = 0.16), and a pooled odds ratio of 2.52 per standard deviation (1.13 to 5.62, p = 0.024). pEMT specificity pointed the same way, with areas under the curve of 0.67 (p = 0.18) and 0.83 (p = 0.037), but its pooled odds ratio of 2.25 (0.99 to 5.12) did not exclude 1. The stromal core was also higher in responders, with areas under the curve of 0.64 in GSE84713 and 0.83 in GSE183881 (p = 0.037), equal to pEMT specificity and the largest in that panel, and it correlated with pEMT specificity at +0.64 and +0.70. The Zhou nine-gene set gave 0.82 in GSE84713 (p = 0.010), the largest in that panel, and 0.78 in GSE183881, and it had the largest pooled odds ratio (3.35, 1.29 to 8.69). 76GS showed no association (0.60 and 0.47). The post hoc combination did not improve on the core (Supplementary Data 32d to 32f). The Basal centroid score gave areas under the curve of 0.79 (p = 0.018) and 0.95 (p = 0.002) and a pooled odds ratio of 3.55 (1.23 to 10.2, p = 0.019). It correlated with the malignant core at only 0.54 and 0.51. Entered together, the malignant core kept no association in either panel (Firth odds ratios 1.63, p = 0.37, and 1.14, p = 0.82), and the Basal score kept its association in GSE183881 (7.65, 1.32 to 1,133, p = 0.019). The subtypes reported by the GSE84713 depositors agree, with 12 of 15 basal tumours responding against 1 of 5 mesenchymal (Supplementary Data 33 to 33e). In both settings the Basal programme carries the cetuximab association at least as well as the malignant arm.

In the GDSC1 screen^39^, 22 head and neck lines had DepMap expression^40^, and no score was associated with sensitivity (malignant core Spearman −0.19, p = 0.39). In a broader squamous panel of 65 lines the malignant core correlated with sensitivity (−0.28, p = 0.027), and pEMT specificity and 76GS did not (Supplementary Data 34). The treated models therefore support the malignant genes without separating them from stroma, and cultured lines add little.

### The arms separate at the protein level, with no increase in EGFR phosphorylation per molecule

Transcripts do not show whether the EGFR pathway is active, so the arms were examined in the proteomes and phosphoproteomes of 108 CPTAC-3 tumours^41^. pEMT specificity correlated with hemidesmosome and laminin-332 protein (median Spearman +0.49) and not with collagen and matricellular protein (−0.01). The canonical pEMT score tracked both (+0.45 and +0.49, Figure 7a, Supplementary Data 35 to 37). The 105 phosphosites on EGFR-to-MAPK cascade proteins did not correlate with pEMT specificity as a set (permutation p = 0.22, Figure 7b). EGFR and ITGB4 sites correlated with it only before adjustment for their parent protein (EGFR Y1172 fell from +0.27 to −0.02, Figure 7c). Tumours with high pEMT specificity carry more of these proteins (EGFR +0.37, ITGB4 +0.56). ERBB2 and ERBB3 did not follow (−0.03 and +0.08), which keeps the median of the EGFR-family set near zero in Figure 7a. The protein data therefore point to more receptor in tumours with high pEMT specificity, with no sign of a more active receptor.

**Figure 7.**
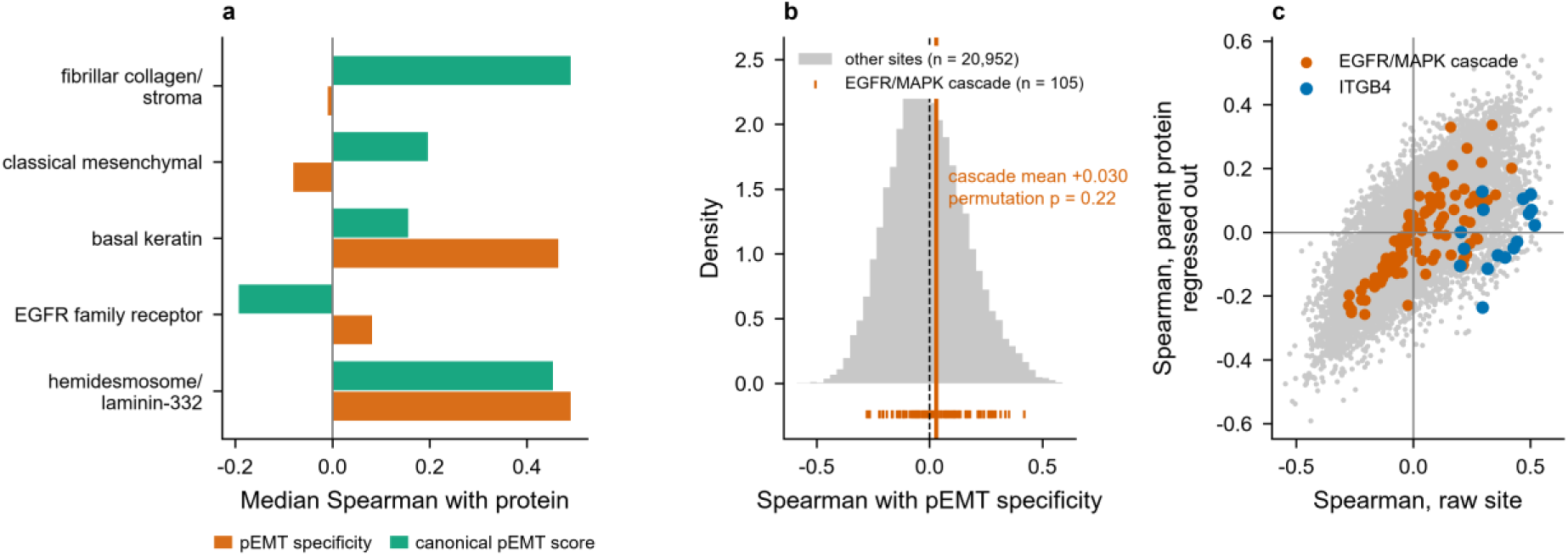
Protein and phosphoprotein correlates of the malignant and stromal arms in CPTAC-3. **a** Median Spearman correlation between each score and the abundance of proteins in five sets fixed in advance, across 108 tumours. Orange is pEMT specificity, green the canonical pEMT score. **b** Distribution of the correlation between pEMT specificity and each of the 20,952 phosphosites outside the cascade, in grey, with the 105 sites on EGFR to MAPK cascade proteins marked beneath as a rug and their mean as a vertical line. The p-value is from 2,000 permutations of the score across tumours, with the set statistic recomputed in full each time. **c** Correlation of each phosphosite with pEMT specificity before and after regressing out the abundance of its own parent protein. Points on the diagonal are unaffected by adjustment. Points falling towards zero on the vertical axis are sites whose correlation is explained by protein abundance. Cascade sites and ITGB4 sites are highlighted.

### In cohorts not selected for cetuximab, the malignant arm does not mark favourable prognosis

A marker of good prognosis in general could explain the cetuximab result, so the arms were tested in four bulk cohorts not selected for cetuximab, TCGA-HNSC, GSE41613^42^, CPTAC-3 and GSE65858^43^, with 869 tumours and 350 deaths. Hazard ratios are per standard deviation and adjusted for stage, age, site and HPV status where recorded (Table 2, Figure 4b–e, Supplementary Figs. 10 to 12, Supplementary Data 38 to 41). Pooled estimates are also given with the Hartung-Knapp adjustment, which suits four heterogeneous cohorts (Tables 2 and 3, Supplementary Data 39b). The canonical signature was the most consistent, with a pooled hazard ratio of 1.23 (1.09 to 1.38, Hartung-Knapp 1.01 to 1.49) and I² of 4%. pEMT specificity gave 1.25 with I² of 64% from no identified source, and its interval excluded 1 under DerSimonian-Laird (1.03 to 1.52) but not under Hartung-Knapp (0.88 to 1.78). GSE41613 gave the largest estimate (1.77), and without it the pooled hazard ratio was 1.15 (1.02 to 1.29, Hartung-Knapp 0.93 to 1.42) with I² of 0%. The adjusted estimate for the malignant core was 1.19 (Hartung-Knapp 0.90 to 1.56), and only its univariable estimate excluded 1 under Hartung-Knapp (1.18, 1.07 to 1.31). The stromal core gave 1.11 (0.87 to 1.43, Table 3, Supplementary Data 3e and 3f). MLR-EMT was not prognostic (1.07, 0.83 to 1.37). Excluding the 148 TCGA patients recorded as receiving systemic therapy left pEMT specificity at 1.15 (Supplementary Data 3g). In these cohorts there was thus no evidence that the malignant arm marks favourable prognosis. They differ from the cetuximab cohort, mostly primary against recurrent or metastatic disease, so the contrast cannot test a treatment interaction, but it argues against good prognosis in general as the explanation of the cetuximab association.

**Table 2.**
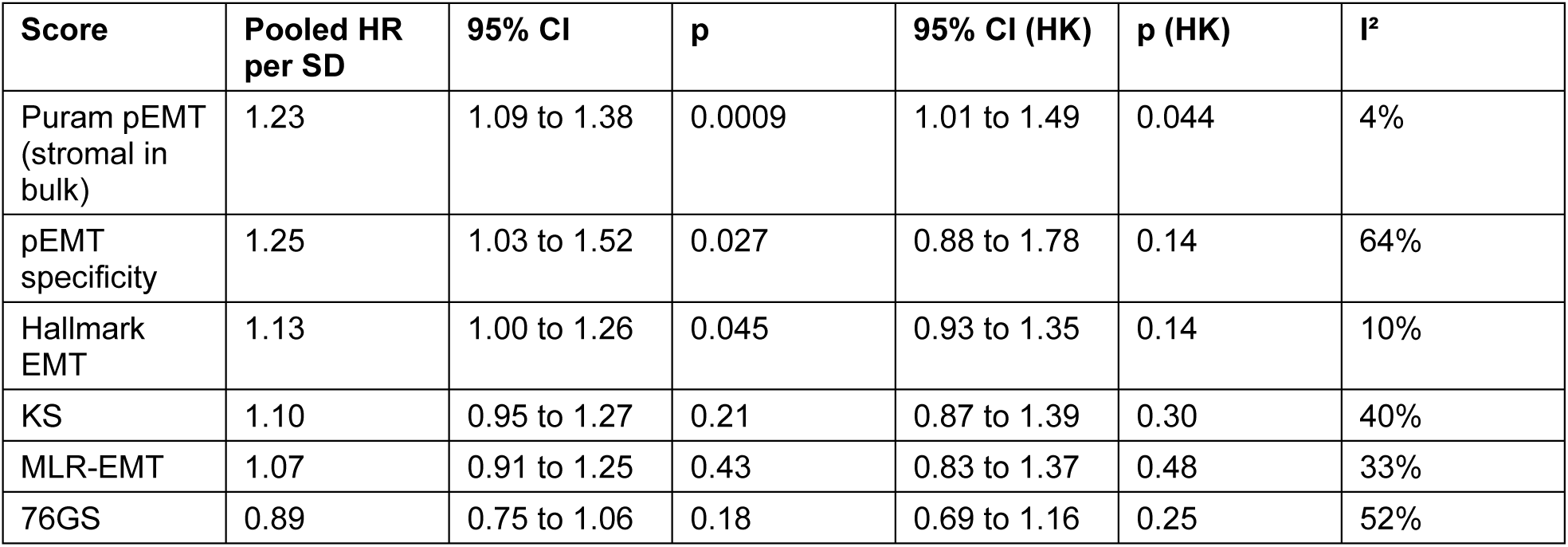
Pooled adjusted hazard ratio per standard deviation for the published scores and the classifier. Random-effects meta-analysis of TCGA-HNSC, GSE41613, CPTAC-3 and GSE65858, comprising 869 tumours and 350 deaths. Each score was standardised within its cohort and fitted with adjustment for pathological stage, age, anatomical site and HPV status where the cohort recorded them. The pooled hazard ratio uses the DerSimonian-Laird estimate of between-cohort variance. The first interval and p-value come from the DerSimonian-Laird estimate, and the second interval and p-value apply the Hartung-Knapp adjustment, which is more conservative with four cohorts. CI, confidence interval. HK, Hartung-Knapp. I² is the percentage of the variation across cohort estimates that is due to heterogeneity and not to sampling error.

**Table 3.** Pooled adjusted hazard ratio per standard deviation for the arms scored directly from their genes. Random-effects meta-analysis over TCGA-HNSC, GSE41613, CPTAC-3 and GSE65858 (869 tumours, 350 deaths). Adjustment and intervals as in Table 2. The cores are the primary direct measures of each arm. The consensus arms are a sensitivity analysis. Univariable estimates are in Supplementary Data 3e and 39b.

| Score | Genes | Pooled HR per SD | 95% CI | p | 95% CI (HK) | p (HK) | I<sup>2</sup> |
| --- | --- | --- | --- | --- | --- | --- | --- |
| Malignant core | 12 | 1.19 | 1.01 to 1.39 | 0.035 | 0.90 to 1.56 | 0.14 | 31% |
| Stromal core | 14 | 1.11 | 0.96 to 1.29 | 0.17 | 0.87 to 1.43 | 0.27 | 40% |
| Malignant arm, total share | 52 | 1.25 | 1.09 to 1.42 | 0.0009 | 1.01 to 1.54 | 0.045 | 10% |
| Stromal arm, total share | 43 | 1.19 | 1.05 to 1.36 | 0.008 | 0.97 to 1.47 | 0.074 | 20% |
| Malignant arm, consensus | 16 | 1.28 | 1.05 to 1.56 | 0.013 | 0.90 to 1.83 | 0.11 | 52% |
| Stromal arm, consensus | 58 | 1.19 | 1.07 to 1.33 | 0.002 | 0.99 to 1.43 | 0.054 | 0% |

### pEMT specificity is not higher in node-positive tumours

Puram and colleagues linked the pEMT programme to nodal metastasis, so we tested that association. In the 11 discovery-dataset patients with recorded nodal status, all of them in the training data, P(pEMT-high) of the primary tumour was higher in node-negative patients, with complete separation (median 0.99 against 0.06, Mann-Whitney U = 0, exact two-sided p = 0.006). This is the opposite of the reported association, and the Puram pEMT signature was also higher in node-negative patients (U = 2, p = 0.024, Figure 3c, d, Supplementary Data 5b). In 420 TCGA tumours pEMT specificity was not associated with nodal stage. Among pathological variables, perineural invasion and histological grade showed weak associations (p = 0.020 and 0.045, Supplementary Fig. 13, Supplementary Data 42, Supplementary Note 2). The nodal association reported for the programme therefore does not hold in these data.

## Discussion

The pEMT programme describes malignant cells when scored in single cells and mostly fibroblasts when scored in bulk tumours. Carcinoma cells transcribe its laminin-332 and hemidesmosome genes, and fibroblasts its fibrillar collagens, metalloproteinases and vimentin, in all three single-cell datasets and within individual patients, and in bulk HNSCC the stromal genes dominate. The malignant part, recoverable from bulk data with a single-cell classifier or a 12-gene core, is hemidesmosome and laminin-332 adhesion with autocrine EGFR ligands in a basal keratinocyte context, with almost no mesenchymal component. We keep the name partial EMT because these are the pEMT genes that malignant cells express, and Puram’s definition included no EMT transcription factor.

Schinke et al. observed the same confounding in three bulk cohorts and removed TCGA tumours with an estimated fibroblast fraction of 48% or more^20^. Filtering tumours keeps malignant and stromal genes in one score, and the excluded tumours receive no score. Dividing the genes lets both arms be scored in every tumour.

Two recent studies support this division. In spatial transcriptomes of HNSCC, Simkin and colleagues found malignant partial EMT led by the laminin-332 and hemidesmosome genes of our malignant core, with COL1A1, COL5A2, MMP2, TAGLN and vimentin in a separate fibroblast programme^27^. They described two modes of malignant partial EMT, one at the invasive edge beside fibroblasts and driven by TGF-β, and one inside tumour nests induced by myeloid cells through oncostatin M. The edge mode places malignant partial EMT next to the stroma that dominates bulk scores, which may be one reason bulk pEMT scores mix the two arms. Bulk profiles cannot tell the two modes apart, and spatial profiling of pre-treatment biopsies could test whether they differ in their association with outcome under cetuximab. In breast cancer, the basal subtype tracks partial more than full EMT^29^. The HNSCC Basal score behaved the same way, correlating positively with partial EMT gene sets, including a mammary one^28^ that malignant cells also express in HNSCC, and negatively with full EMT and fibroblast gene sets.

Among 40 patients treated with platinum and cetuximab, the malignant arm was higher in those with long progression-free survival by every definition we tested, and the stromal arm was not. Bossi et al. found basal traits and strong EGFR signalling in the same patients with long progression-free survival^37^, and Zhou et al. identified nine genes that each separated long from short progression-free survival, five of them hemidesmosome or laminin-332 components and receptors^32^. A Basal-subtype score discriminated better than either malignant measure, and the two could not be separated, so in this cohort the malignant arm is best read as the part of the pEMT list that carries the Basal programme, in line with Bossi et al. and with the malignant-basal subtype of Puram et al.^15^. The malignant core also did not outperform epithelial-differentiation scores or the Zhou gene set derived on the same patients. In two xenograft panels it was higher in responders, but there it could not be separated from the stromal genes, and the Basal score again did as well or better. The partition therefore explains why pEMT scores and hemidesmosome-based predictors disagree. A single-arm cohort has no treatment interaction to estimate, so the malignant arm is associated with outcome under cetuximab and its value for predicting benefit remains untested. In cohorts not selected for cetuximab there was no evidence that it marks favourable prognosis, which makes general good prognosis an unlikely explanation.

MLR-EMT, implemented as published, called most bulk tumours hybrid and was not associated with outcome. In a cohort without epithelial tumours, a more hybrid tumour retains more of the epithelial state, so MLR-EMT measures a different quantity from the malignant arm. pEMT specificity does not measure a mesenchymal state (its correlation with vimentin is +0.02 and with ZEB1 −0.11 in TCGA).

The arms also separate in the proteome of an independent set of patients, where tumours with high pEMT specificity carry more EGFR and ITGB4 protein without more phosphorylation per molecule, and receptor abundance alone failed as a biomarker in RTOG 0522. Signatures opposing the pEMT-high profile were enriched for MEK and EGFR inhibitors against random gene sets, although no class reached a false discovery rate of 0.05, and drug and CRISPR screens showed no matching effect on viability (Supplementary Note 2, Supplementary Figs. 14 to 18, Supplementary Data 43 to 50), as expected of a perturbation that changes cell state without killing cells, as cetuximab and erlotinib do in oral squamous cell carcinoma lines^44^.

Network proximity pointed to the pharmacology of activated fibroblasts. In a neoadjuvant window study of the MEK inhibitor trametinib in oral cavity cancer, 11 of 17 evaluable patients had a clinical response, a reduction in tumour size of at least 25% on examination^45^.

The cetuximab cohort has 40 patients in two extreme groups, which inflates discrimination, and the confidence interval for pEMT specificity is wide (0.62 to 0.92). All patients received platinum, so the association with outcome under cetuximab cannot be separated from chemosensitivity or from prognosis within recurrent disease. Progression-free survival is available only as a dichotomy. The malignant arm could not be separated from the Basal subtype in the patient cohort, and in the xenografts the Basal score did as well or better. Several arm definitions were examined, and three were added after the first GSE65021 results. The xenograft panels are small and cannot separate the two arms, and they define response in the mouse. The prognostic value of pEMT specificity varied between cohorts (I² of 64%). The subtype result rests on TCGA alone. Both malignant measures fall as non-malignant cells dilute a tumour, so they partly track purity, although adjusting for stromal and immune content did not weaken the cetuximab association. BayesPrism did not fully separate the arms on RNA-seq, and the L1000 signatures come from cell lines.

The findings can be tested by scoring pre-treatment biopsies for the malignant core, pEMT specificity, the Basal subtype and epithelial differentiation in a larger cetuximab-treated cohort with unselected outcomes, ideally with a comparator arm. The malignant core and the Basal score both kept their association when each patient was scored alone, so a targeted assay appears feasible, whereas pEMT specificity would need a reference for each platform. Scoring the paired specimens of the trametinib window study would test the reversal prediction.

In summary, the pEMT gene list joins two programmes made by different cells. Bulk pEMT scores mostly report the fibroblast programme. The malignant programme, which can be scored in any bulk profile from 12 genes or with the single-cell classifier, is a basal, hemidesmosome-rich epithelial state with little mesenchymal character. This malignant part, together with the Basal subtype it accompanies, was associated with outcome under cetuximab in patients and xenografts, and the stromal part was not. Dividing a single-cell signature gene by gene between the cells that express it applies to any programme defined in single cells and scored in bulk tissue, and for pEMT it should precede use as a biomarker.

## Methods

### Single-cell datasets and quality control

Six single-cell HNSCC and related datasets were obtained from TISCH2 with harmonised cell-type annotation^46^: HNSC_GSE103322 (Puram et al., 5,902 cells, 2,488 malignant, 1,454 fibroblast and myofibroblast), LSCC_GSE150321^24^ (12,687 cells from two patients deposited as two GEO samples, 6,172 malignant, 451 fibroblast and myofibroblast), OSCC_GSE172577^47^ (57,503 cells, 7,914 malignant, no annotated fibroblasts), NPC_GSE150430^48^ and NPC_GSE162025^49^ (nasopharyngeal, 45,959 and 176,440 cells, no annotated fibroblasts), and THCA_GSE148673^50^ as a non-head-and-neck outgroup. Cells with fewer than 200 detected genes and genes detected in fewer than 10 cells were removed, and counts were library-size normalised to 10,000 per cell and log-transformed. Only GSE103322 and GSE150321 contain both malignant cells and fibroblasts and were therefore the only datasets eligible for training.

The remainder were held out for validation. Because GSE150321 contributes few fibroblasts, the classifier learns the fibroblast class mainly from GSE103322. A seventh dataset, GSE181919, was used only to replicate the partition of the pEMT gene list (below).

Programme scores were computed as the mean z-score across the genes of each Puram et al. programme^15^ present in the dataset, requiring at least three present genes.

### Partition of the pEMT programme by cell type

For each of the 100 genes of the Puram pEMT programme, the share of total expression contributed by malignant cells, fibroblasts and myofibroblasts, endothelium and immune cells was computed across all 5,902 cells of GSE103322, together with the proportion of cells of each type in which the gene was detected. Shares were computed on the linear scale, in counts per 10,000. The TISCH2 matrices are the natural logarithm of one plus counts per 10,000 (the back-transformed values of every cell sum to 10,000), so they were back-transformed before summing. A gene was assigned to the malignant arm by total share when its malignant share exceeded its fibroblast share and to the stromal arm otherwise. Gene family membership (laminin-332 and hemidesmosome, classical mesenchymal, remainder) was used for annotation and for a one-sided Mann-Whitney comparison of malignant share between families, and entered no other calculation. Within malignant cells, the log2 fold change between pEMT-high and epithelial-like cells was computed from mean linear expression with a pseudocount of 0.1.

Because share of total expression depends on the number of cells of each type a dataset contains, the partition was repeated with a composition-independent measure, the malignant share of mean expression per cell (the mean in malignant cells divided by the sum of the means in malignant cells and fibroblasts), in GSE103322, in GSE150321 and in GSE181919^25^.

GSE181919 was obtained from GEO as a UMI count matrix with the authors’ cell annotation. It was normalised to counts per 10,000, and restricted to primary tumour and lymph-node metastasis tissue, giving 5,113 malignant cells and 1,981 fibroblasts from 20 patients. In GSE181919 the per-cell measure was also computed separately in each patient with at least 20 malignant cells and 20 fibroblasts (13 patients). Agreement between datasets is reported as Spearman correlation of the per-cell shares and Cohen’s kappa for the assignment of each gene to the malignant or fibroblast side of parity.

### Bulk cohorts

TCGA-HNSC STAR-derived transcripts per million were obtained from the Genomic Data Commons and transformed to log₂(TPM + 1), with primary tumours identified from the sample barcode. CPTAC-3 HNSCC RNA-seq^41^ was assembled from the same GDC STAR workflow (GENCODE v36), collapsing Ensembl identifiers to gene symbols by highest mean expression and retaining one tumour per patient, the first primary-tumour aliquot by sample identifier.

Overall survival was taken from the current GDC clinical release. The frozen 2021 publication snapshot records only 16 deaths against the current 40. GSE41613 (Affymetrix U133 Plus 2)^42^ and GSE65858 (Illumina HT-12 v4)^43^ were parsed from GEO series matrices, with probes collapsed to gene symbols by highest mean probe and values log₂-transformed where the platform reported linear intensities.

### Weighted gene co-expression network analysis

WGCNA was run on log₂ TPM in primary tumours^51^ using the 8,000 most variable genes and a signed network. The soft-thresholding power was selected automatically as the lowest power achieving a scale-free topology fit of R² ≥ 0.80, which gave 13. Modules were detected by dynamic tree cutting with a minimum module size of 30, deep split 2, and a merge cut height of 0.25, followed by a partitioning-around-medoids reassignment step. Module eigengenes were correlated with classifier probabilities and with each EMT score by Pearson correlation. Enrichment of each module for the Puram programmes, Hallmark EMT and immune and stromal marker sets was tested by hypergeometric test with Benjamini-Hochberg correction. Network seeds were defined as module members with module membership kME ≥ 0.7.

### Training-label assignment

Labels were assigned within malignant cells by quantile thresholds on the Puram pEMT and epithelial differentiation scores. Cells above the 75th centile of the pEMT score were labelled pEMT-high and cells below the 25th centile that were also above the 75th centile of the epithelial score were labelled epithelial-like. To prevent the classifier learning confounded programmes, cells above the 90th centile of either the hypoxia or the cell-cycle score were excluded from both malignant classes. Annotated fibroblasts and myofibroblasts formed the third class under a strict definition that admits no other stromal cell type. All remaining cells were excluded from training.

### Pseudobulk construction

Training examples were synthetic mixtures of cells, so that the classifier is trained on the same kind of object it is applied to. For each training dataset, 1,000 mixtures were generated by drawing Dirichlet-distributed class proportions and sampling 100 cells accordingly, and, because bulk expression is the logarithm of summed expression, each mixture summed the sampled cells’ linear counts per 10,000, renormalised the sum to 10,000 and log-transformed it. As a sensitivity analysis, the classifier was refitted on mixtures of the same cells built as the mean of their log-normalised profiles. The refitted classifier’s pEMT specificity correlated with the main score at 0.98 in TCGA and in GSE65021, shared 55 of the 100 largest pEMT-high coefficients, and gave an area under the curve for prolonged progression-free survival in GSE65021 of 0.74 (0.58 to 0.89), against 0.78 for the main classifier (Supplementary Data 51 and 51b). A mixture received a class label only if that class contributed at least 60% of its cells. Mixtures without a dominant class were discarded. Datasets contributing fewer than 10 cells of a class, or fewer than 50 labelled cells in total, did not contribute that class.

For the single-cell validation in Figure 3, pseudobulks were instead constructed from real groupings: one profile per malignant (patient, anatomical site) combination and one per patient for fibroblasts, requiring at least 20 cells per profile.

### Multinomial elastic-net classifier

A multinomial logistic regression with elastic-net penalty was fitted with the SAGA solver, an L1 ratio of 0.5, inverse regularisation strength C of 0.1, tolerance 10⁻³ and a maximum of 10,000 iterations, on genes passing a variance floor of 10⁻⁶. Cross-validation was grouped by dataset, which with two training datasets is leave-one-dataset-out. Performance is reported as held-out accuracy and one-versus-rest area under the receiver operating characteristic curve per class. The held-out fold was standardised with the training fold’s scaler, whereas each bulk cohort is standardised within itself. The gene variance filter was applied to all training pseudobulks before the folds were split. Standardising each held-out dataset within itself, as for bulk cohorts, gave an accuracy of 0.997 and a pEMT-high area under the curve of 1.000 (Figure 1d, Supplementary Data 51).

The classifier was benchmarked against ridge logistic regression, random forest and gradient boosting (LightGBM, both on all genes and on the 2,000 most variable) on identical folds.

Agreement between learners is reported as the Spearman correlation of the pEMT-high probability. Every learner was also evaluated with the held-out dataset standardised within itself. For the two linear models only the fitted logistic layer was then applied, and the tree models, which have no scaler, were refitted on the training fold standardised with its own scaler. The elastic net was retained for coefficient interpretability.

### Projection to bulk transcriptomes

For each bulk cohort, the classifier genes present in the cohort were z-scored gene-wise within that cohort, genes absent from the cohort were given a z-score of zero, and the fitted coefficients were then applied. Within-cohort standardisation makes the projection insensitive to platform and batch but renders the class probabilities cohort-relative, so absolute proportions are not interpretable and the score cannot be computed for a single sample in isolation. We summarise the projection as pEMT specificity, defined as P(pEMT-high) − max(P(epithelial-like), P(fibroblast/stromal-like)). Because pEMT specificity is bounded at ±1, the same probabilities were also summarised as log P(pEMT-high) − log max(P(epithelial-like), P(fibroblast/stromal-like)), with probabilities floored at 10⁻⁶, a value no tumour reached, and analysed with the same procedures as a sensitivity analysis.

### Label-free recovery of the pEMT state

All 533 genes belonging to any of the six Puram programmes were removed from the malignant cells of GSE103322, leaving 17,708. Patients with fewer than 20 malignant cells, genes detected in fewer than 10 cells, and genes without variance within any patient were excluded so that batch correction could be run, leaving 2,441 cells from 14 patients. The 2,000 most variable genes were selected, corrected for patient with ComBat, scaled, and reduced to 30 principal components, from which a nearest-neighbour graph and Leiden clustering at resolution 1.0 were computed. Labels were used at no point in this pipeline. Agreement between clusters and the training labels was measured by adjusted Rand index and adjusted mutual information against 1,000 label permutations, and separation in principal-component space by silhouette against 200 label permutations. The supervised test fitted L2-penalised logistic regression on the same non-Puram genes under leave-one-patient-out cross-validation, pooling held-out predictions for a single area under the curve.

### Comparison with the Tyler and Tirosh decoupling approach

An EMT signature gene set was assembled following the description in their paper: the MSigDB Hallmark EMT set, the Tan et al. tumour signature, and the 250 genes most correlated in TCGA-HNSC with their six seed regulators SNAI1, SNAI2, TWIST1, VIM, ZEB1 and ZEB2, taken from ‘initial_genes’ in their published code, with the top-250 cut matching the ‘genes_filter_fun’ they apply to HNSC. Correlations with the seeds were Spearman correlations, computed by rank transform across the 520 primary tumours. The assembled set comprised 643 genes present in TCGA, of which 633 were detected in GSE103322. Each gene was assigned to a compartment in GSE103322 according to whether malignant cells or fibroblasts contributed the larger share of its total expression on the linear scale, giving 209 cancer-compartment and 424 fibroblast-compartment genes, and the 520 primary tumours were scored on each compartment as a mean per-gene z-score. This approach implements their decoupling principle but does not reproduce their pipeline, which additionally uses CCLE reference profiles and deconvolves within molecular subtype with per-subtype gene weighting. Tyler and Tirosh deposited neither per-sample TCGA scores nor their deconvolved HNSC gene sets, so the comparison uses this implementation.

### Direct scores of the two arms

Each arm was scored in every bulk cohort as the mean per-gene z-score of its genes, like the published signatures. All gene sets were taken from single-cell data only, but they were not all specified at the same time, and Supplementary Data 3h gives when each was fixed relative to the first analysis of GSE65021. The cores (12 genes with a malignant share of total expression of at least 80% and 14 with a fibroblast share of at least 60%) and the total-share arms (52 and 43 genes) were specified together, as rules applied to the discovery partition, before any arm score was examined in GSE65021. Their membership was recomputed when the partition was corrected to the linear scale, after the first GSE65021 results were known. The malignant core was unchanged. The stromal core gained vimentin and four other genes, so its current membership was set after the cohort had been examined. The cores are the primary direct measures, because the 52-gene arm includes genes near parity that carry fibroblast signal in bulk and correlates +0.97 with the canonical signature in TCGA. After the first GSE65021 results, three sensitivity analyses were added: alternative core thresholds from 60% to 90%, removal of the Zhou nine-gene set, and the consensus arms (the 16 and 58 genes on the same side of parity by the per-cell measure in all three single-cell datasets). The direct arm scores are pooled over the four survival cohorts like every other score.

### Composition simulation

Tumour and lymph-node cells of GSE181919 were converted to counts per 10,000 with library sizes over all genes. Malignant cells were split by their own Puram pEMT score, the mean per-gene z-score of log1p expression over the pEMT genes, into a pEMT-high pool (top quartile, 1,279 cells) and a pEMT-low pool (bottom quartile, 1,279 cells). Fibroblasts (1,981 cells) and immune cells (T, B and plasma, macrophage and dendritic cells) formed the other pools. Each synthetic tumour comprised 300 cells drawn with replacement, under a full factorial design over the share of malignant cells drawn from the pEMT-high pool (0, 0.25, 0.5, 0.75, 1), the fibroblast fraction (0.05, 0.15, 0.30, 0.45, 0.60) and the immune fraction (0.05, 0.20, 0.35), with six replicates, giving 450 tumours. Each tumour was the mean of its cells’ linear counts per 10,000, log1p-transformed. The 450 tumours were scored as one bulk cohort, with pEMT specificity computed with within-cohort standardisation and each gene set scored as a mean per-gene z-score. Each standardised score was regressed on the three standardised design factors, and the partial R squared of each factor was computed by dropping it from the model.

### Reference-based deconvolution

BayesPrism 2.2.3^30^ was run in R 4.4.2 with the tumour and lymph-node cells of GSE181919 as the reference: malignant cells with each patient as a separate cell state, fibroblasts, endothelial cells, T cells, B and plasma cells, myeloid cells, mast cells and myocytes, with at most 3,000 cells per type. The targets were the STAR gene counts of the 520 TCGA-HNSC primaries and the linear intensities of GSE65021, a microarray, for which the method is an approximation.

Malignant-cell and fibroblast expression (Z) was scaled to counts per million of each cell type’s total per sample, log2(x + 1) transformed and standardised per gene within the cohort, and gene sets were scored as mean per-gene z-scores. For TCGA the fractions are those of the initial Gibbs step, because the second step, which refines the fractions only, was stopped after seven hours, and Z comes from the initial step in either case.

### EMT score panel

Five published scores were computed on log₂ expression in primary tumours. 76GS is computed as described by Chakraborty et al.^16^: the expression of each of the 76 Byers et al. genes^17^ is weighted by that gene’s Pearson correlation with CDH1 in the cohort, the weighted values are summed and centred, and the score is oriented towards the epithelial pole. KS is the per-sample two-sample Kolmogorov-Smirnov statistic between the mesenchymal and epithelial gene distributions of the Tan et al. tumour signature^12^, signed as in the reference implementation. Hallmark EMT^52^ and the Puram programmes^15^ are mean per-gene z-scores.

MLR-EMT is the ordinal model of George et al.^18^, implemented as in the reference code of its authors (csb-iisc/EMT_Scoring_RNASeq on GitHub, formerly Cancer-Systems-Biology-Lab) and the published supplement^16^. RNA-seq values, log₂(TPM + 1), are first mapped to a microarray-like scale (0.57 + 0.37 × log₂(TPM + 1)). Microarray data enter on their log₂ scale. Each cohort is normalised by a single offset, the mean of 20 published normaliser genes in the cohort minus their mean in the 59 NCI-60 training lines, subtracted from every value. The predictors are CLDN7 and the ratio of VIM to CDH1, and the published ordinal coefficients give the probabilities of the epithelial, hybrid and mesenchymal states. The score is P(hybrid) if P(epithelial) exceeds P(mesenchymal) and 2 − P(hybrid) otherwise, on a scale from 0 (epithelial) through 1 (hybrid) to 2 (mesenchymal). Besides the score, the state probabilities were retained, and each tumour was assigned to its most probable state. The probability of the hybrid state, P(hybrid), corresponds to partial EMT. Two details differ from the reference code and follow the published method: NCI-60 rows are located by probe identifier instead of row number, and the same 20 normaliser genes are averaged in the cohort and in NCI-60. Applied to the NCI-60 training lines the implementation returns 12 epithelial, 35 hybrid and 12 mesenchymal lines.

Because the RNA-seq state calls depend on the mapping, both RNA-seq cohorts were also scored with log₂(TPM + 1) entered directly. The offset moved from −2.43 to −1.30 in TCGA, and 263 of 520 tumours were called hybrid, 253 epithelial and 4 mesenchymal. In CPTAC-3, 94 of 110 were hybrid and 16 epithelial. The score kept its ranking of tumours (Spearman 0.997 and 0.998 with the mapped score), and the correlation of the hybrid probability with pEMT specificity in TCGA was 0.36, against 0.35 with the mapped score. The microarray cohorts, GSE65021 among them, do not use the mapping. A simplified implementation that standardises each predictor within the cohort, uses the difference of VIM and CDH1 in place of their ratio and sums the state probabilities differently gives materially different results in GSE65021 (Supplementary Data 17b), so only the published implementation is used.

### Published gene sets and molecular subtypes

Gene lists were taken from the supplementary files of the source publications and not reconstructed. They were the 171-gene EGFR-induced EMT signature of Schinke et al., three sets from Zhou et al. (46 functional differentially expressed genes, a 59-gene invasive network and 9 cetuximab progression-free-survival predictors) and the 28-gene tumour budding signature of Ourailidis et al. Each was scored as a mean per-gene z-score. Overlap between the genes with the 100 largest positive pEMT-high classifier coefficients and each published set was tested by hypergeometric test against the 23,436-gene classifier universe.

TCGA four-class expression subtypes were taken from the supplementary tables of the TCGA 2015 HNSCC publication and cross-checked against the UCSC Xena PanCanAtlas subtype table, which agreed on all 279 samples. Scores were compared across subtypes by Kruskal-Wallis test with Mann-Whitney comparisons of each subtype against the remainder. GSE65858 publishes its own de novo consensus clusters, which we analysed separately and did not treat as equivalent to the TCGA calls. In GSE65021, subtypes were assigned with the 728-gene centroids that the TCGA HNSCC study derived from the classifier of Walter et al. and deposited on its GDC publication page. Each centroid gene was median-centred within the cohort, and each sample was given the subtype whose centroid it correlated with most strongly (Pearson, 674 shared genes). The Basal score was the correlation with the Basal centroid, and the procedure reproduced all 279 published TCGA calls when applied to the TCGA classification matrix. The 41 cancer meta-programmes of Gavish et al.^26^ were taken from their Supplementary Table 2 and scored as mean per-gene z-scores, with Benjamini-Hochberg correction over the 41 programmes for progression-free survival in GSE65021. Two further partial EMT gene sets served as external controls. The mammary EMT signature of Knutsen et al.^28^ was taken as its three clusters (EMT_down, 88 genes, EMT_partial, 81 genes, and EMT_up, 95 genes) from Supplementary Data 1 of Sahoo et al.^29^, and the spatial meta-programmes of Simkin et al.^27^ (partial EMT, 47 genes, fibroblast and epithelial, 50 genes each) from the file Final_Metaprograms_Extended.csv deposited with that study. Their cell of origin was measured in GSE103322 as for the pEMT genes. Each set was scored as a mean per-gene z-score in TCGA-HNSC and GSE65021. Correlations of the Basal score with two sets were compared with the Steiger test for dependent correlations, and sets associated with progression-free survival were entered with the Basal score in a joint Firth model.

### Cetuximab-treated cohort

GSE65021^37^ comprises 40 patients with recurrent or metastatic HNSCC treated with first-line platinum plus cetuximab, deposited as two extreme progression-free survival groups: 14 patients exceeding 12 months and 26 below 5.6 months, balanced by the original investigators on performance status, weight loss, prior radiotherapy, tumour grade, primary site and residual disease at the primary site. Tumour specimens were taken from the recurrence where available and otherwise from the primary, giving 31 primary and 9 recurrence or metastasis specimens. Expression was measured on the Illumina Whole-Genome DASL HumanHT-12 v4 BeadChip (GPL14951), 29,377 probes, deposited on a linear scale.

Because GPL14951 is the DASL variant of the HumanHT-12 v4 array and shares the ILMN probe namespace with GPL10558, probe-to-symbol mapping used the GPL10558 annotation, which resolves 27,548 of the 29,377 probes (93.8%) to 19,436 unique symbols. Values were log₂-transformed and probes collapsed to the highest-mean probe per gene, as for GSE41613 and GSE65858. The classifier was applied exactly as for every other bulk cohort, with within-cohort z-scoring of the classifier genes. Of the 694 genes with a non-zero coefficient, 669 are present.

No survival times are deposited, so a time-to-event model was not possible and the analysis is a two-group comparison: Mann-Whitney U with rank-biserial correlation as the effect size and two-sided p-values with Benjamini-Hochberg correction across the scores tested, receiver operating characteristic area under the curve with a 2,000-sample bootstrap percentile interval, and logistic regression of prolonged progression-free survival per standard deviation of each score, unadjusted and adjusted for age, stage, grade, prior radiotherapy, oral cavity and oropharynx site indicators, and specimen type. With 14 events, maximum-likelihood estimates were unstable whenever a strong epithelial score or the seven covariates were in the model, so odds ratios are reported from Firth-penalised logistic regression^53,54^ with profile penalised-likelihood intervals and penalised likelihood-ratio p-values. The implementation was checked against direct numerical maximisation of the penalised likelihood and reproduces the Haldane correction on a separated two-by-two table. Maximum-likelihood estimates are retained in Supplementary Data 23, 52 and 53. Head-to-head models entered pEMT specificity and one other score together, both standardised. Separate models paired each score in which higher values mean a less epithelial state (MLR-EMT, KS) with each epithelial score (76GS, Puram epithelial differentiation). A combined marker, the equal-weight sum of standardised pEMT specificity and standardised Puram epithelial differentiation, was specified after the single-score results were known, with 76GS and sign-reversed MLR-EMT as alternative epithelial components in sensitivity analyses, and is reported as exploratory. Its area under the curve is reported with a 2,000-resample bootstrap percentile interval and a paired bootstrap interval for the gain over pEMT specificity alone, alongside the leave-one-out cross-validated area under the curve of a fitted two-term logistic model. A sensitivity analysis refitted the comparison on the 31 primary-tumour specimens. To test whether the associations reflect tumour purity, the malignant core and pEMT specificity were each refitted with the stromal core, with an immune score, and with both. The immune score was the mean per-gene z-score of 18 markers fixed in advance of this analysis (PTPRC, CD2, CD3D, CD3E, CD247, CD8A, CD19, MS4A1, CD79A, CD68, CD14,

CD163, LYZ, FCGR3A, ITGAM, CSF1R, CD74 and HLA-DRA). Each primary measure was compared with 76GS, Puram epithelial differentiation, the canonical Puram pEMT signature and the Zhou nine-gene set by the paired bootstrap distribution of the difference in area under the curve (2,000 resamples of patients). For multiplicity, Benjamini-Hochberg correction was applied over one family comprising every independently derived published score and the four arm scores specified before the cohort was examined. Because every score is standardised within the cohort, transportability was assessed in two ways: by standardising each patient against the other 39 patients only, and by scoring each patient alone against the gene means and standard deviations of GSE65858, an external cohort on the whole-genome version of the same Illumina array family. The Basal centroid score and the malignant core were scored the same ways, the centroid against gene medians, and also with no reference, as the correlation of the patient’s uncentred log2 expression with the Basal centroid and the mean log2 expression of the 12 core genes.

The nine cetuximab predictor genes of Zhou et al.^32^, scored here as one set, were identified in this cohort, and five of them are among the classifier’s top 100 coefficients, so the set is reported as a positive control only and excluded from the multiple-testing correction.

With 14 patients in one group and 26 in the other the cohort is powered for a large effect only, and the two groups were selected as outcome extremes, so the discrimination reported here is an upper bound on what would be seen in an unselected population.

### Patient-derived xenograft panels

GSE84713 was used as deposited (GCRMA-normalised log2 expression on Affymetrix HG-U133 Plus 2). Probes were collapsed to gene symbols by highest mean probe, and the responder label was taken from the series matrix. The deposit dichotomises response from tumour volume at a cut-off it does not state. The two pairs of models derived from one patient (11269A and B, 11437A and B, concordant in label) were averaged for the primary per-patient analysis. For GSE183881, whose depositors classified reads as human or mouse with Xenome and discarded mouse reads before mapping, human isoform-level TPM from the deposited RSEM output was summed to genes through the GENCODE v38 transcript-to-HGNC map and log2-transformed, and the 17 models with a cetuximab response label were analysed. Scores were computed with within-panel standardisation, as for every cohort, and compared between responders and non-responders by two-sided Mann-Whitney test, bootstrap area under the curve and Firth odds ratio per standard deviation. The Mann-Whitney test is the primary inference. The bootstrap percentile interval is descriptive and can reach 0.5 when the p-value is just below 0.05. The two panels were pooled by inverse-variance weighting of the log odds ratios, with standard errors approximated from the profile-likelihood intervals. For the 14 GSE84713 models also reported by Klinghammer and colleagues in 2020^55^, cetuximab relative tumour volume was transcribed from their Table 1 and correlated with each score by Spearman correlation. The paired bootstrap comparison of areas under the curve and the equal-weight combination of pEMT specificity with Puram epithelial differentiation were applied as in the patient cohort, without refitting. The Basal centroid score was computed in each panel as in the patient cohort, as the Pearson correlation of median-centred expression of the centroid genes with the TCGA 2015 Basal centroid, and was compared with the malignant core by a joint Firth model, the paired bootstrap difference in area under the curve and the pooled odds ratio.

Response was also tabulated against the molecular subtypes reported by the GSE84713 depositors.

### Cell lines

Cetuximab response (GDSC1 drug 1114, fitted area under the dose-response curve and ln IC50) was taken from the GDSC release 8.5 fitted dose-response file and matched by Sanger model identifier to DepMap expression (log2(TPM + 1), default entry per model). The primary panel was the head and neck lines with expression data (22 of 35). A secondary squamous panel added oesophageal squamous (Oncotree ESCC), lung squamous and cervical squamous lines (65 in total). Scores were standardised within each panel, and each was correlated with the dose-response area and ln IC50 by Spearman correlation.

### Cell-cell communication

Ligand-receptor inference used LIANA’s rank aggregate^56,57^, a consensus of CellPhoneDB, Connectome, log2FC, NATMI and SingleCellSignalR, on the log-normalised matrix with a minimum expression proportion of 0.1 and 1,000 permutations. The share of each ligand’s expression contributed by each cell group was computed on the linear scale, as for the partition. Cell groups were fibroblasts and myofibroblasts as one CAF group, malignant cells split into pEMT-high, epithelial-like and remaining by the training labels, and endothelial and immune cells retained for context. Specificity of a pair for one malignant state over the other was measured as the difference in log10 specificity rank. Because the specificity rank is a permutation p-value and pEMT-high cells outnumber epithelial-like cells, the comparison was repeated with pEMT-high downsampled to the epithelial-like cell count under five random seeds, and a pair was reported only if it favoured pEMT-high in at least four of the five matched runs.

### CPTAC-3 proteome and phosphoproteome

TMT proteome at gene level and phosphoproteome at site level for CPTAC-3 HNSCC tumours were taken from LinkedOmics, and joined to the classifier scores already computed for this cohort by CPTAC case identifier, giving 108 tumours with all three data types. Protein-level analysis correlated each score with the abundance of each member of five gene sets fixed in advance (hemidesmosome and laminin-332, EGFR family receptors, fibrillar collagen and stroma, classical mesenchymal, basal keratin), by Spearman correlation with Benjamini-Hochberg correction, summarised as the median correlation per set.

Phosphosites quantified in fewer than 30 tumours were dropped, leaving 21,057. Each was correlated with pEMT specificity by Spearman with Benjamini-Hochberg correction across all sites. Because a phosphosite rises with its parent protein, each site was also correlated after regressing the site on the abundance of its own parent protein in the same tumours by ordinary least squares and taking the residual. The adjusted correlation is treated as evidence of altered phosphorylation, and the unadjusted one is retained because adjustment also removes activation driven by receptor abundance. The EGFR to MAPK cascade set was specified in advance as EGFR, ERBB2, ERBB3, SHC1, GRB2, SOS1, HRAS, KRAS, NRAS, RAF1, BRAF, ARAF, MAP2K1, MAP2K2, MAPK1, MAPK3, RPS6KA1, RPS6KA3, ELK1, JUN, FOS, PTPN11, GAB1 and CBL. Its statistic is the mean correlation over its sites, tested against a null in which the score is permuted across tumours and the whole statistic recomputed 2,000 times, so that the correlation between sites measured on the same tumours is preserved under the null.

### Survival analysis

Overall survival was modelled by Cox proportional hazards regression on complete cases, with every score standardised to unit variance so that hazard ratios are per standard deviation. Adjusted models included pathological stage, age, anatomical site (oropharynx as reference) and HPV status where the cohort recorded them. GSE41613 records only stage and age. Kaplan-Meier curves are shown by tertile with log-rank tests and numbers at risk. Concordance was assessed by repeated five-fold cross-validation. Head-to-head models placed pEMT specificity and one published score in the same adjusted model, with likelihood-ratio tests for the addition of either to the other. In TCGA, a sensitivity analysis excluded patients whose legacy clinical record (targeted_molecular_therapy) marks systemic therapy, since the harmonised clinical files carry no drug names.

### Meta-analysis and sensitivity analyses

Per-cohort log hazard ratios and their standard errors were taken directly from the fitted Cox models instead of being back-calculated from confidence intervals, and pooled by inverse-variance weighting under both fixed-effect and DerSimonian-Laird^58^ random-effects models. Because the DerSimonian-Laird interval can be too narrow when few studies are pooled and heterogeneity is present, every random-effects estimate was also given with the Hartung-Knapp adjustment^59^, which rescales the variance of the pooled estimate by the weighted residual heterogeneity and takes the interval and p-value from a t distribution with three degrees of freedom. Heterogeneity is reported as Cochran’s Q, I² and τ². Because a pooled estimate from four studies is sensitive to specification, every score was pooled under four combinations of univariable or adjusted models with fixed-effect or random-effects pooling. Sensitivity was further assessed by leave-one-cohort-out pooling, by truncating follow-up in every cohort at 36 and 60 months, and, in GSE65858, by stratifying on the treatment modality that cohort records (Supplementary Data 54). Subgroup analyses standardised each score within the subgroup so that all estimates remain per standard deviation.

### LINCS signature reversal

The top 150 pEMT-high classifier coefficients and the top 150 epithelial-like coefficients were submitted to the SigCom LINCS two-sided enrichment interface^60^ as up and down sets against the L1000 compound, CRISPR knockout and shRNA libraries, with up to 2,000 signatures returned per direction. Perturbagens were ranked by the number of signatures reversing the query against the number mimicking it, tested by a one-sided sign test for an excess of reversing signatures, with Benjamini-Hochberg correction. The test is one-sided because the question is directional. Four compounds reached a false discovery rate of 0.05, and all four remain significant under a two-sided test (Supplementary Data 43b). Mechanism-of-action annotation was taken from the PRISM compound metadata, since the LINCS metadata does not carry it. Specificity was assessed against 50 random gene sets of matched size, drawn from the same 4,000-gene universe, with the same random seed, as an initial control that used five sets, with an add-one empirical p-value for each mechanism class and Benjamini-Hochberg correction across classes.

### DepMap sensitivity and dependency

PRISM secondary-screen dose-response curves were filtered^61^ to rows passing short-tandem-repeat profiling, and the area under the curve was aggregated as the median per compound and model. Compounds screened in at least five matched models were correlated with pEMT specificity by Spearman correlation with Benjamini-Hochberg correction, within the HNSCC lineage and within a wider pan-squamous set. CRISPR dependency used Chronos gene effects^62^ over the same models.

Mechanism-of-action classes and network-predicted compound sets were tested against a permutation null over cell lines instead of by a rank-sum test across compounds. Compounds correlated with the same phenotype in the same models are not independent observations, and a rank-sum test comparing one class against all other compounds is anti-conservative. For the permutation test, the area under the curve was ranked within each compound across the models in which it was screened, the phenotype was ranked across all matched models, and the class statistic, the mean Spearman correlation over the compounds of the class, was recomputed over 5,000 permutations of the phenotype labels across cell lines. Observed and permuted statistics are computed in the same way, so the test is exact under the label-permutation null.

### Network proximity

Drug targets were taken from the curated targets of DrugBank^63^, in the licensed database export of 19 April 2025 held by the Milner Therapeutics Institute, and, separately, from STITCH associations^64^. The interactome was the physical subnetwork of STRING v11.0^65,66^, restricted to edges with experimental or curated-database evidence and a combined physical score of at least 400, mapped to gene symbols, giving 13,285 proteins. Proximity was computed as in Guney et al.^67^ as the mean shortest-path distance from a drug’s targets to the nearest seed gene, standardised against 1,000 degree-matched random draws of both the targets and the seed genes, with a minimum degree-bin size of 100. A drug was called a hit at z ≤ −1.96 with normal-approximation false discovery rate ≤ 0.05. Because a single seed gene can dominate the statistic, each hit was retested with every seed gene removed in turn, and a hit was retained only if it remained significant in the worst case. Three seed sets were used: the canonical Puram pEMT genes within module M06, the M13 module hubs, and the top 100 positive pEMT-high classifier coefficients. A fourth, independent of the classifier, comprised the 52 Puram pEMT genes assigned to the malignant arm by total share, of which 48 are in the interactome and none is an EGFR-family gene. It was run with the same code, random seed and null, with and without exclusion of each drug’s targets that are themselves seeds.

### Statistics and reproducibility

Analyses used Python 3.11 with scanpy^68^, scikit-learn, lifelines, statsmodels, scipy, igraph, matplotlib and LIANA. WGCNA was implemented in Python following the steps of the R package, with the hybrid tree cut from the Python port of dynamicTreeCut. Correlations are Spearman unless stated. Multiple testing is controlled by the Benjamini-Hochberg procedure^69^ within each family of tests, and families are stated with each result. Tests are two-sided unless stated. Random seeds are fixed throughout and recorded in the configuration file, which is distributed with the code so that every figure and table is reproducible from a single script that runs every stage in order.

### Ethics and data use

This study analysed only publicly available, de-identified human data and generated no new patient samples. Ethical approval and informed consent for the collection and sharing of every dataset used were obtained by the original investigators and are documented in the source publications and repository records: TCGA-HNSC and CPTAC-3 under their respective consortium protocols, and GSE103322, GSE150321, GSE172577, GSE150430, GSE162025, GSE148673, GSE181919, GSE41613, GSE65858 and GSE65021 under the approvals reported by their depositing institutions. The xenograft panels GSE84713 and GSE183881 were generated under the human-tissue and animal approvals reported by their investigators, and the cell lines are established public resources. No additional ethical approval or consent was required for the secondary analyses reported here, which were conducted in accordance with the Declaration of Helsinki and with the data-use terms of each repository.

## Data availability

All data are public except DrugBank, which was used under an academic licence. TCGA-HNSC was obtained from UCSC Xena and the GDC, and CPTAC-3 from the GDC. GSE103322, GSE150321, GSE172577, GSE150430, GSE162025 and GSE148673 were obtained through TISCH2, and GSE41613, GSE65858, GSE65021, GSE181919, GSE84713 and GSE183881 from GEO. The CPTAC-3 proteome and phosphoproteome came from LinkedOmics, and the Hallmark gene sets from MSigDB. DepMap 24Q4 and PRISM came from the DepMap portal, GDSC release 8.5 from the Genomics of Drug Sensitivity in Cancer website, SigCom LINCS through its public API, and the MLR-EMT reference files from the csb-iisc/EMT_Scoring_RNASeq repository (formerly Cancer-Systems-Biology-Lab). We also used STRING v11.0, STITCH v5.1.5 and the GENCODE v38 HGNC metadata. The DrugBank export (19 April 2025) cannot be redistributed, so the supplementary tables give each drug’s DrugBank accession, name and approval status but no DrugBank descriptive text. Supplementary Data 1 to 55 are provided as separate Excel workbooks, with their titles and legends in the Supplementary Information. Supplementary Notes 1 to 3 and Supplementary Figs. 1 to 18 are in the Supplementary Information.

## Code availability

All analysis code is available at https://github.com/GehadYoussef/pemt-hnscc under the MIT licence and is archived at Zenodo (https://doi.org/10.5281/zenodo.23065628)^70^. The code in src/ is organised as numbered stages that share one package. Every parameter and random seed is in config/config.yaml, the small public reference files the code reads are in data/references/, and data/README.md lists the source of every other input. The scripts run_all.sh and run_all.ps1 execute the stages in order and reproduce every figure and supplementary table from the raw inputs, and the README names the script behind each one. The partition of the pEMT gene list and its replication are in src/09_gene_partition/, the analyses of the cetuximab-treated patients, xenografts and cell lines in src/10_cetuximab/, the composition simulation, proteome and ligand-receptor analyses in src/11_composition_and_mechanism/, the revision analyses (Hartung-Knapp pooling, the Basal centroid comparator, the log-odds summary, the Gavish meta-programmes, the BayesPrism deconvolution, the single-patient scoring, the xenograft Basal comparison and the external partial EMT controls) in src/13_revision/, and the MLR-EMT implementation and Firth regression in src/pemt/.

## Acknowledgements

D.Z.R.A. received stipend support from Animal Free Research UK. G.Y. was funded by the Milner Therapeutics Institute and Cancer Research UK. M.K.J. was supported by the Ramanujan Fellowship (SB/S2/RJN-049/2018) awarded by the Science and Engineering Research Board, Government of India. This research was supported by the National Research Foundation of Korea grant funded by the Ministry of Science and ICT (RS-2025-18362970) and the Brain Pool Plus Fellowship Program funded by the Ministry of Science and ICT (RS-2025-25427881). The funders played no role in study design, data collection, analysis and interpretation of data, or the writing of this manuscript.

An earlier version of this work, “Partial EMT Prognostic Classifier and Drug Repurposing for HNSC”, was presented by D.Z.R.A. at the 103rd General Session of the IADR (Barcelona, Spain, 25 to 28 June 2025), where it received first place in the IADR Hatton Award, Senior Basic Science category. The authors thank the patients and investigators who contributed to TCGA, CPTAC, and the GEO and TISCH2 datasets analysed here.

## Author contributions

D.Z.R.A.: formal analysis, investigation, writing (original draft). S.L.: data curation. P.C. and M.K.J.: methodology, advice on EMT scoring, writing (review and editing). A.B. and M.P.P.: guidance on EMT biology, writing (review and editing). G.Y.: conceptualisation, methodology, formal analysis, supervision, writing (original draft, review and editing). N.H.: supervision, resources, funding acquisition, writing (review and editing). All authors read and approved the final manuscript.

