## Supplementary Information for "Malignant and stromal partial EMT programmes and cetuximab outcome in head and neck cancer"

### Supplementary Notes

**Supplementary Note 1. Signalling between fibroblasts and the two malignant states.** The two arms sit in different cells, so we looked for ligand-receptor signalling between them (Methods). In the discovery dataset, the fibroblast-to-malignant pairs most specific to pEMT-high cells were collagens, fibronectin and thrombospondins signalling to integrin $\alpha3\beta1$. All twelve leading pairs held when group sizes were equalised (Supplementary Fig. 8, Supplementary Data 20). ITGA3 is among the classifier's largest coefficients. The EGFR ligands came from the malignant cells themselves, which supplied 94% of TGFA and NRG1 expression, 84% of EGF and 75% of EREG.

**Supplementary Note 2. Nodal status and pathological variables.** In the single-cell discovery dataset, 11 patients have a primary-site malignant pseudobulk and recorded nodal status. P(pEMT-high) separated the two groups completely in the direction opposite to the one Puram and colleagues reported: median 0.99 in the 4 node-negative patients against 0.06 in the 7 node-positive patients (Mann-Whitney $U = 0$, exact two-sided $p = 0.006$). pEMT specificity gave the same separation ($U = 0$), and the Puram pEMT signature itself was also higher in node-negative patients (median 0.58 against $-0.26$, $U = 2$, $p = 0.024$, Figure 3c, Supplementary Data 5b). All 11 patients belong to the training dataset, whose malignant cells defined the class labels, so this comparison is small and not independent of the classifier. In the five patients with matched primary and nodal samples the two sites showed no consistent difference (Figure 3d, Supplementary Data 5c). Across 420 TCGA-HNSC tumours with recorded pathological N stage, pEMT specificity was not associated with nodal status (node-positive against node-negative, two-sided $p = 0.091$). Of the pathological variables in the TCGA clinical record, perineural invasion and histological grade show weak associations ($p = 0.020$ and $0.045$, Supplementary Fig. 13, Supplementary Data 42). The association between pEMT and nodal metastasis reported in the source study is therefore not reproduced here, and in the small single-cell sample it is reversed.

*Cell line screens.* No PRISM compound reaches significance across 23 HNSCC models (Supplementary Fig. 16a), no module or drug-target gene set shifts towards CRISPR dependency across 63 models (Supplementary Fig. 16b), and no mechanism class reaches significance under a permutation null over cell lines (EGFR inhibitors $p = 0.09$, MEK inhibitors $p = 0.66$, Supplementary Figs. 15 and 16c, Supplementary Data 45 to 48).

*Network proximity.* Network proximity of approved drugs to each seed set (Supplementary Figs. 17 and 18, Supplementary Data 49) links the stromal module M06 to the pharmacology of activated fibroblasts, with 13 hits that survive seed removal among plasminogen-activation and matrix metalloproteinase inhibitors, but nothing specific to the malignant arm. Hits for the classifier's own genes are dominated by EGFR-directed drugs whose single target, EGFR, is itself a seed, and none of these survives removal of that seed. Four approved hits with several targets do survive: amivantamab, the EGFR–MET bispecific antibody, through its MET target, and the ALK inhibitors crizotinib, ensartinib and brigatinib. When we instead seed on the 52 malignant-arm pEMT genes of the total-share partition, which are independent of the classifier and include no EGFR-family gene, and we also exclude each drug's own seed targets, no approved hit survives seed removal with DrugBank targets and two survive with STITCH targets: urokinase, anchored on the plasminogen-activation genes PLA1 and F3, and doxycycline, anchored on MMP10 (Supplementary Data 50). We make no repurposing claim from proximity.

**Supplementary Note 3. Drug signatures, cell line screens and network proximity.** Drugs that might reverse the pEMT-high state were sought among L1000 perturbation signatures. Compared with 50 random gene sets of matched size, signatures opposing the pEMT-high profile were enriched for MEK, EGFR, src, ephrin and other tyrosine kinase inhibitors but not for proteasome or NFkB inhibitors. Each enriched class exceeded every random set (empirical $p = 0.02$, the smallest attainable). MEK inhibitors contributed the largest share of reversing signatures (4.7% against 0.8% for random gene sets). After correction across all classes none reached a false discovery rate of 0.05 (Supplementary Fig. 14, Supplementary Data 43 and 44). Drug and CRISPR screens in 23 and 63 HNSCC lines showed no matching effect on viability (Supplementary Figs. 15 and 16, Supplementary Data 45 to 48). Network proximity analysis pointed to the pharmacology of activated fibroblasts and to nothing specific to the malignant arm (Supplementary Figs. 17 and 18, Supplementary Data 49 and 50).

**Supplementary Fig. 1. Soft-thresholding power for the co-expression network.** **a** Signed scale-free topology fit index against soft-thresholding power, with the dashed horizontal line at the fit threshold of 0.80. **b** Mean connectivity against the same power, on a logarithmic axis. The vertical black line in both panels marks the selected power of 13, the lowest power at which the fit threshold is met.

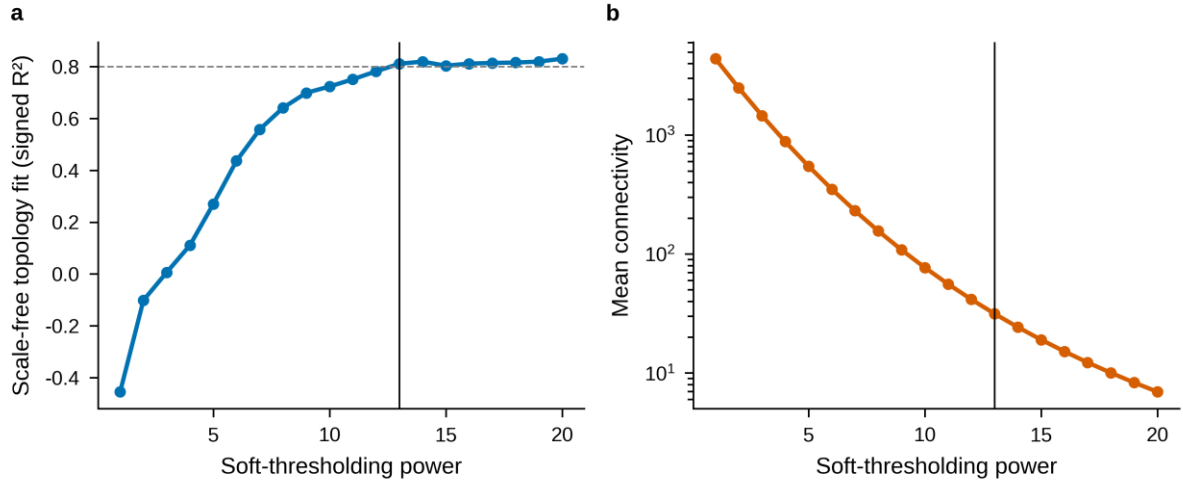

**Supplementary Fig. 2. Co-expression modules and their relationship to the classifier and to the EMT scores.** Heatmap of the Pearson correlation between each module eigengene and each trait, across 520 TCGA-HNSC primary tumours. Rows are the thirteen modules with their sizes, columns are the classifier class probabilities, pEMT specificity and the published EMT scores. Red indicates positive correlation and blue negative, as shown by the colour bar. Module M06 carries 54 of the 76 Puram pEMT genes present in the network and module M13 tracks pEMT specificity. Both are shown with their correlations in Figure 2e.

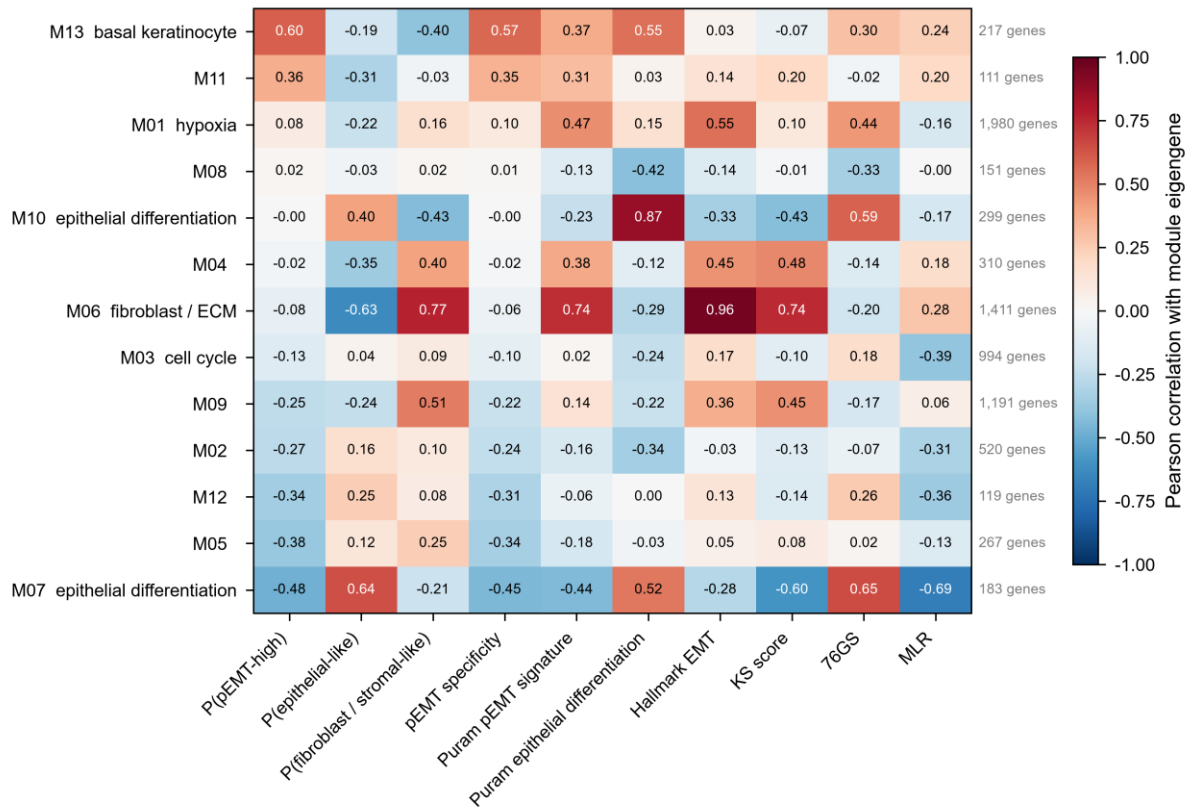

**Supplementary Fig. 3. The three classes in the discovery dataset.** Malignant cells and fibroblasts of GSE103322 on the published two-dimensional embedding, one point per cell. **a** TISCH2 cell type annotation, with the number of cells of each type. **b** Classifier probability of the pEMT-high class. **c** Classifier probability of the fibroblast or stromal-like class. **d** Puram pEMT signature, as the mean z-score of its genes. **e** Hallmark epithelial-to-mesenchymal transition score. **f** The mesenchymal minus epithelial mean z-score of the Tan et al. tumour signature, the gene sets underlying the KS score. Panels b and c share a common scale from 0 to 1.

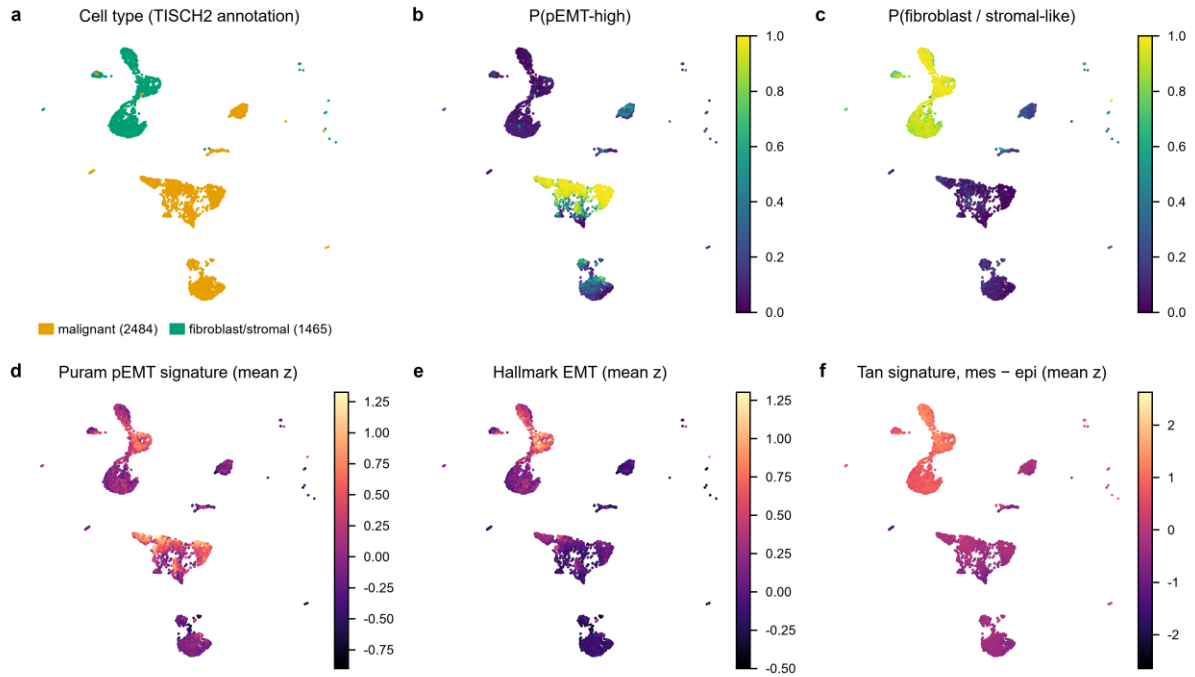

**Supplementary Fig. 4. The classifier across all six single-cell datasets.** One row per dataset, each on that dataset's published embedding, one point per cell. The left column shows the TISCH2 cell type annotation with cell numbers, the middle column the classifier probability of the pEMT-high class on a common scale from 0 to 1, and the right column the Puram pEMT signature as the mean z-score of its genes.

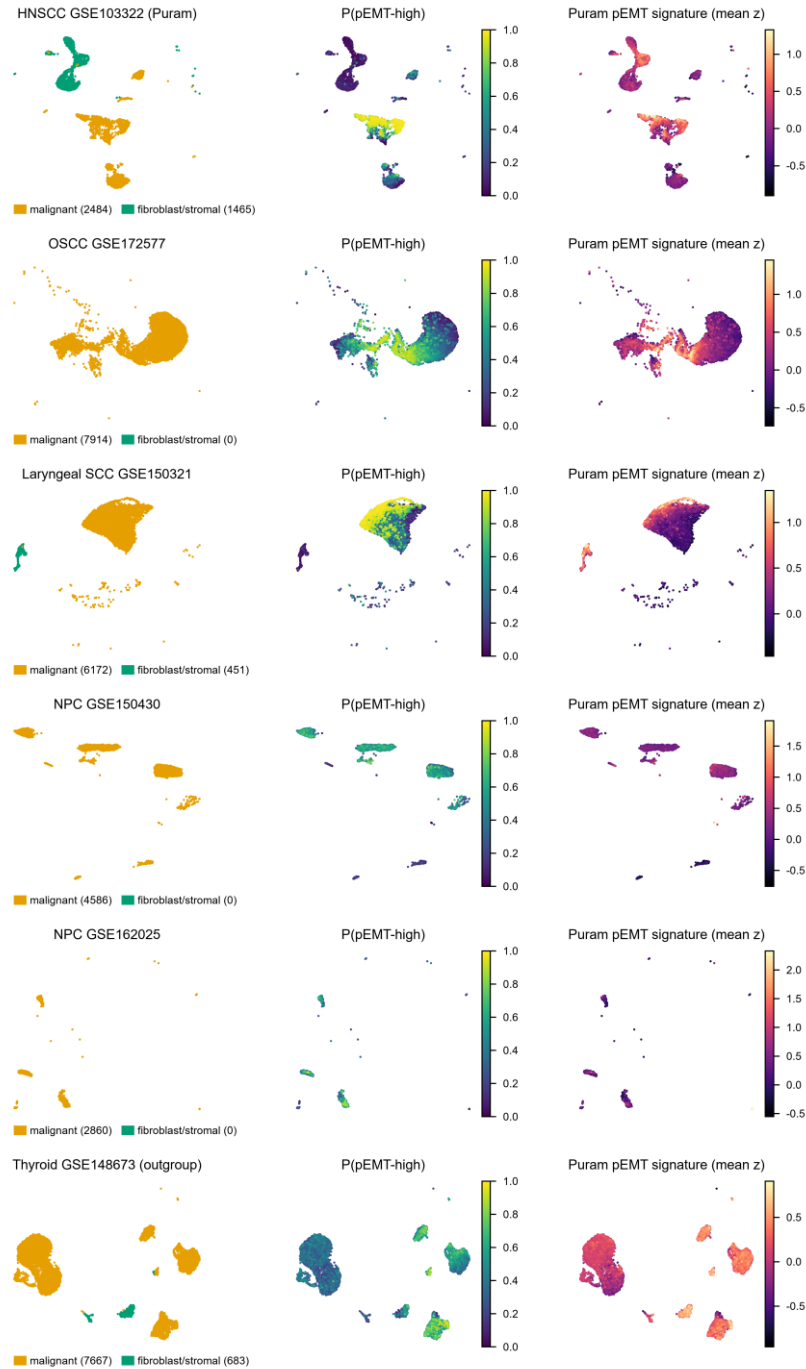

**Supplementary Fig. 5. Classifier benchmarking.** Five learners fitted on identical leave-one-dataset-out folds. Each point is one held-out dataset and the horizontal bar is the mean across folds. **a** Accuracy of the argmax class assignment. **b** Area under the receiver operating characteristic curve for pEMT-high against the other two classes. **c** Log loss, for which lower is better. **d** Spearman correlation between each learner's pEMT-high probability and that of the elastic net, across held-out pseudobulks. The elastic net is shown in orange and the alternative learners in blue. LGBM denotes gradient boosting and LGBM top2k the same learner restricted to the 2,000 most variable genes.

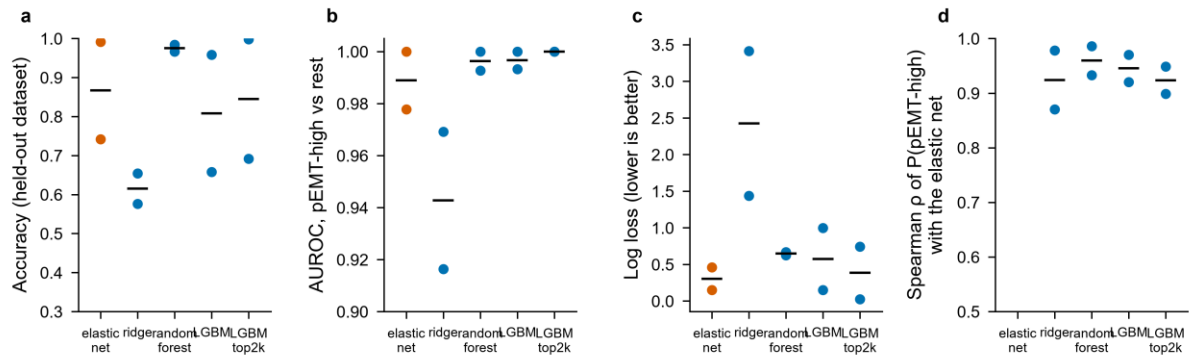

**Supplementary Fig. 6. The pEMT state recovered without the genes that define it.** All 533 genes of the six Puram programmes were removed before any step of this analysis. **a** Malignant cells of GSE103322 on a UMAP built from the remaining genes, with cells carrying a training label coloured and all others grey. **b** The same embedding, with each Leiden cluster coloured by the percentage of its labelled cells that are pEMT-high. Clusters containing no labelled cells are grey. **c** The percentage of labelled cells that are pEMT-high per cluster, with the dashed line at the overall pEMT-high proportion among labelled cells. Labels were used at no point in constructing the embedding or the clusters.

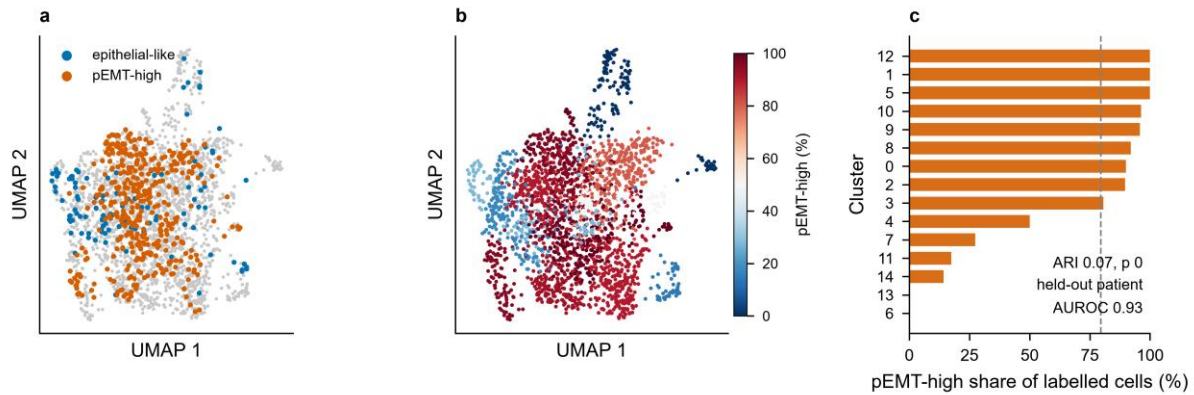

**Supplementary Fig. 7. Comparison with the Tyler and Tirosh decoupling approach. a**

Spearman correlation between each score and the two compartment scores obtained by applying their decoupling principle to the 520 TCGA-HNSC primaries, with their EMT signature gene set resolved into 209 cancer-compartment and 424 fibroblast-compartment genes using GSE103322 on the linear scale. **b** pEMT specificity against their cancer-compartment score, one point per tumour. The analysis applies their published decoupling principle. It does not use their software, which relies on cell line references and subtype-stratified deconvolution.

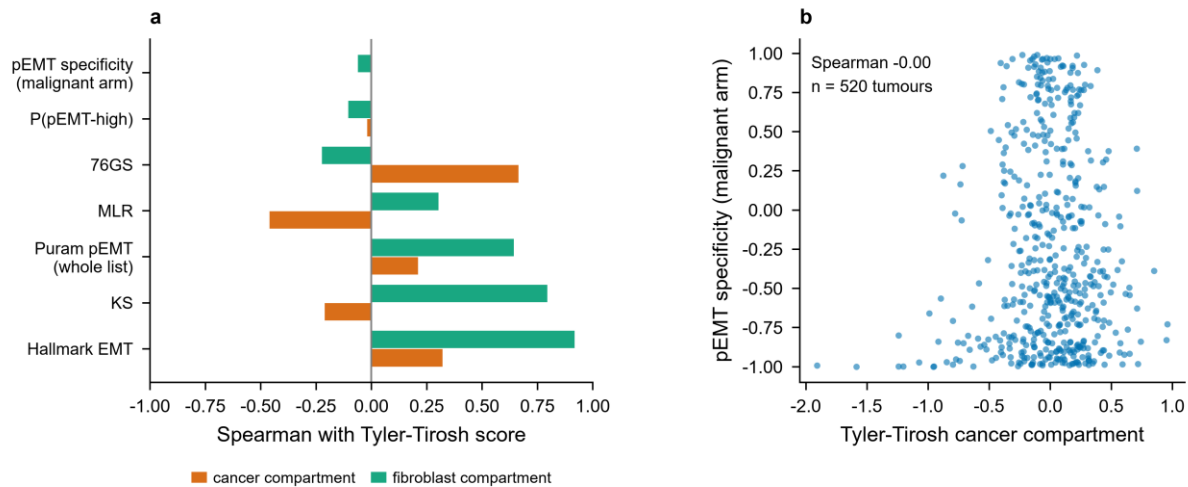

**Supplementary Fig. 8. Signalling between fibroblasts and the two malignant states. a** Share of total expression (linear scale) of each EGFR-family ligand contributed by malignant cells, fibroblasts and other cell types. **b** The same for canonical fibroblast-derived ligands, as a positive control showing that the measurement recovers known stromal sources. **c** Ligand-receptor pairs from fibroblasts, ranked by how specific they are to pEMT-high malignant cells, shown against the same pair's specificity for epithelial-like cells. Specificity is the LIANA consensus rank under a permutation null, plotted as its negative logarithm so that longer bars indicate greater specificity. Every pair shown remained pEMT-high-specific in at least four of five runs with the two malignant groups downsampled to equal cell numbers.

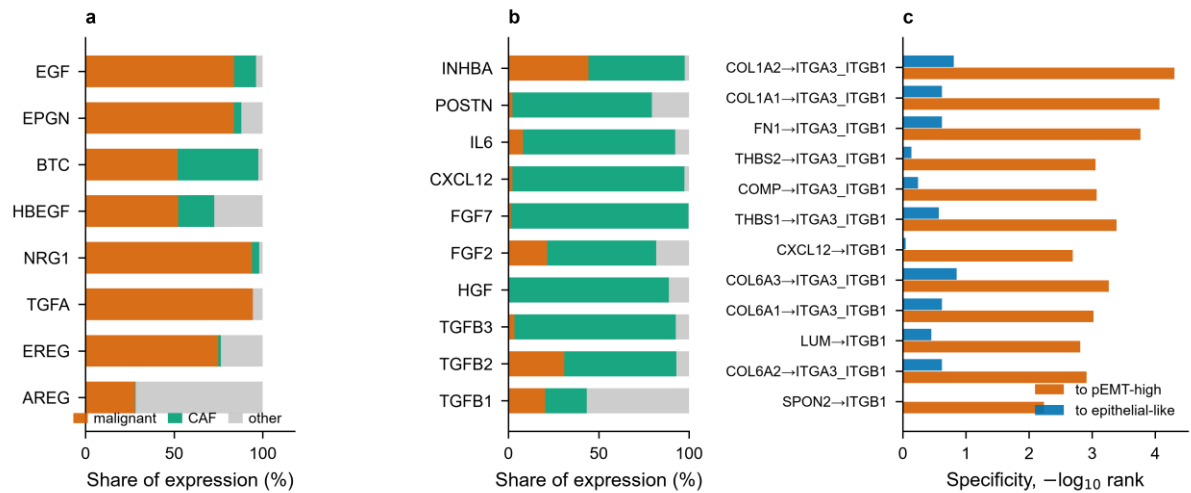

**Supplementary Fig. 9. Exploratory combination of pEMT specificity with epithelial differentiation in GSE65021.** Receiver operating characteristic curves for prolonged progression-free survival on platinum plus cetuximab for pEMT specificity (orange), Puram epithelial differentiation (blue) and their equal-weight combination of standardised scores (black), with the 2,000-resample bootstrap interval of the combination. The combination was specified after the single-score results were known and is evaluated on the same 40 patients in two extreme outcome groups. It is exploratory, and its performance here is an upper bound on what it would achieve in an unselected cohort.

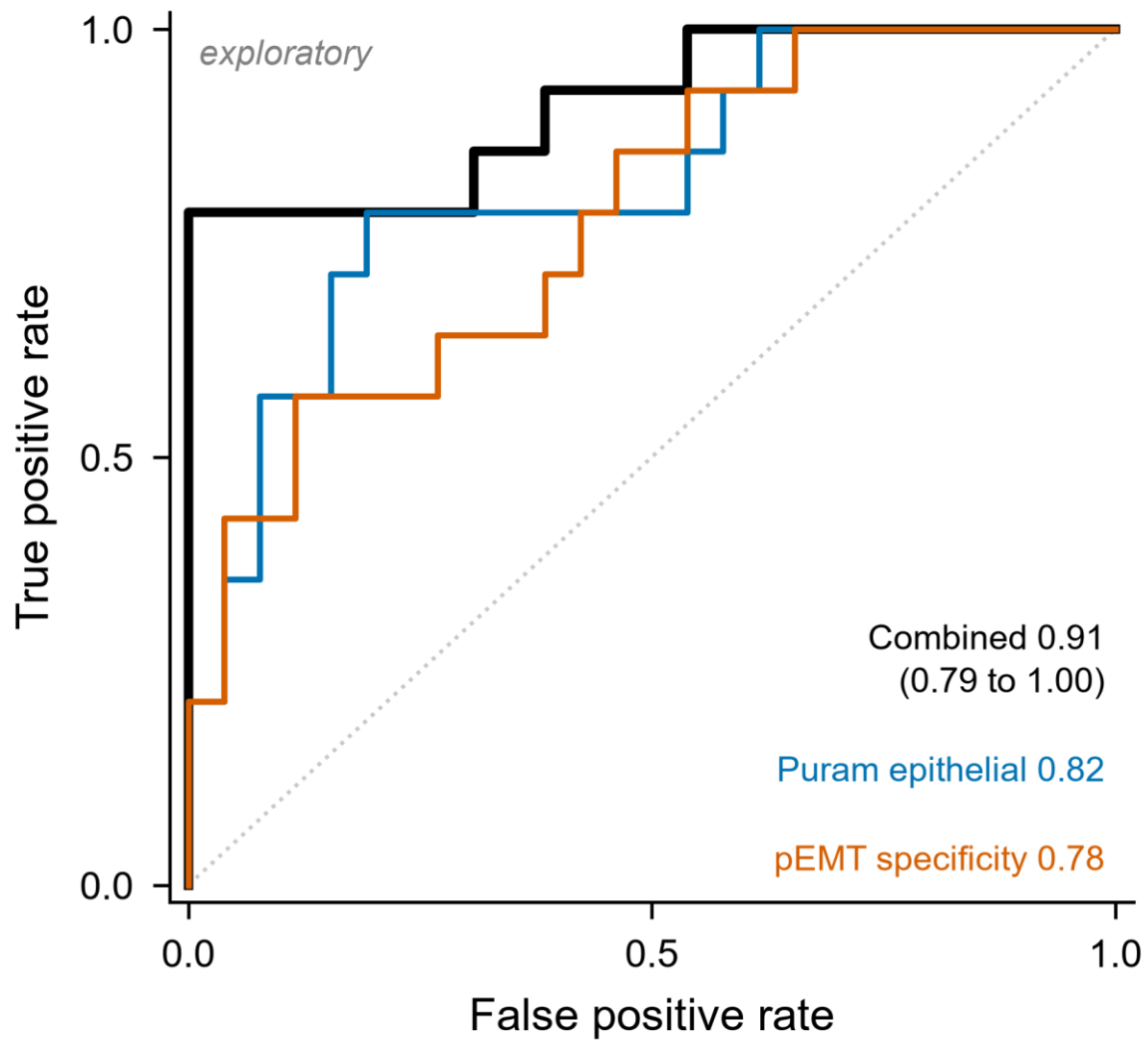

**Supplementary Fig. 10. Survival in the two microarray cohorts.** Kaplan-Meier curves by tertile of pEMT specificity with numbers at risk, and the adjusted hazard ratio per standard deviation for every score, in GSE65858 and GSE41613.

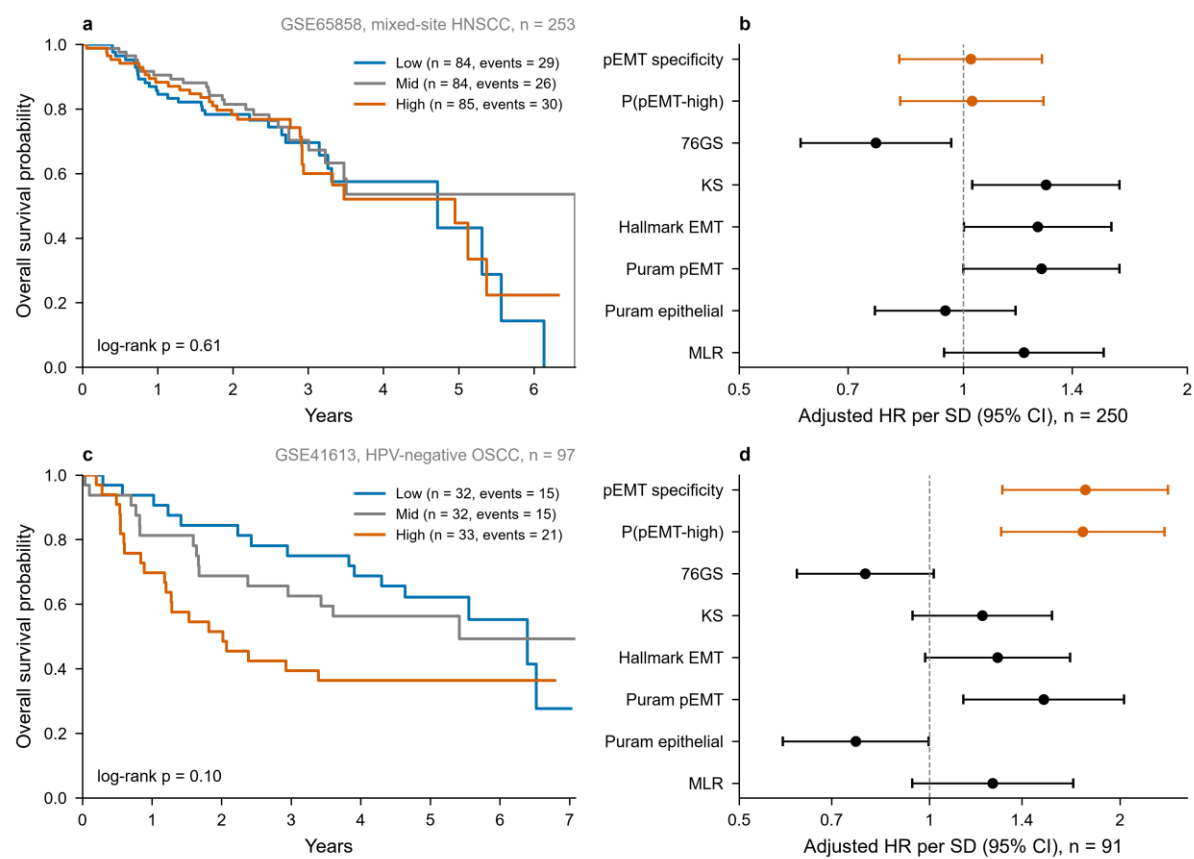

**Supplementary Fig. 11. Adjusted Cox model in TCGA-HNSC.** Forest plot of the hazard ratio for each covariate with its 95% confidence interval, in the adjusted single-cohort model. The estimates pooled across cohorts are in Figure 4e. The model is adjusted for pathological stage, age, HPV status and anatomical site, with oropharynx as the reference site, in 424 tumours with 178 deaths. The vertical black line marks a hazard ratio of 1.

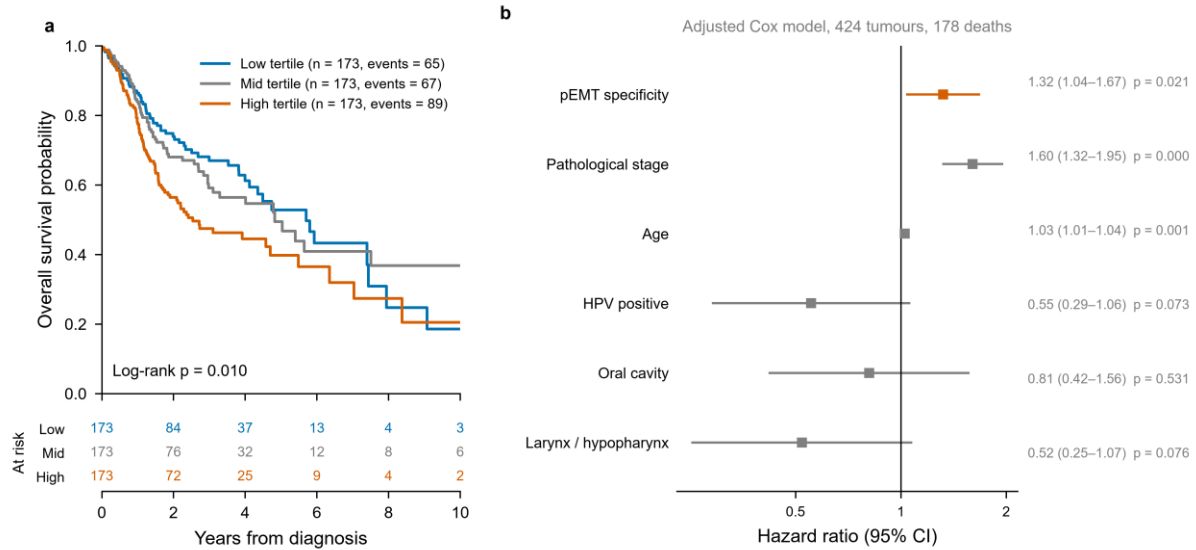

**Supplementary Fig. 12. Survival association by subgroup.** Adjusted hazard ratio per standard deviation for three scores within each anatomical site, HPV stratum and cohort. Grey marks a subgroup with fewer than 30 events. Each score is standardised within the subgroup, so every estimate is per standard deviation of that subgroup.

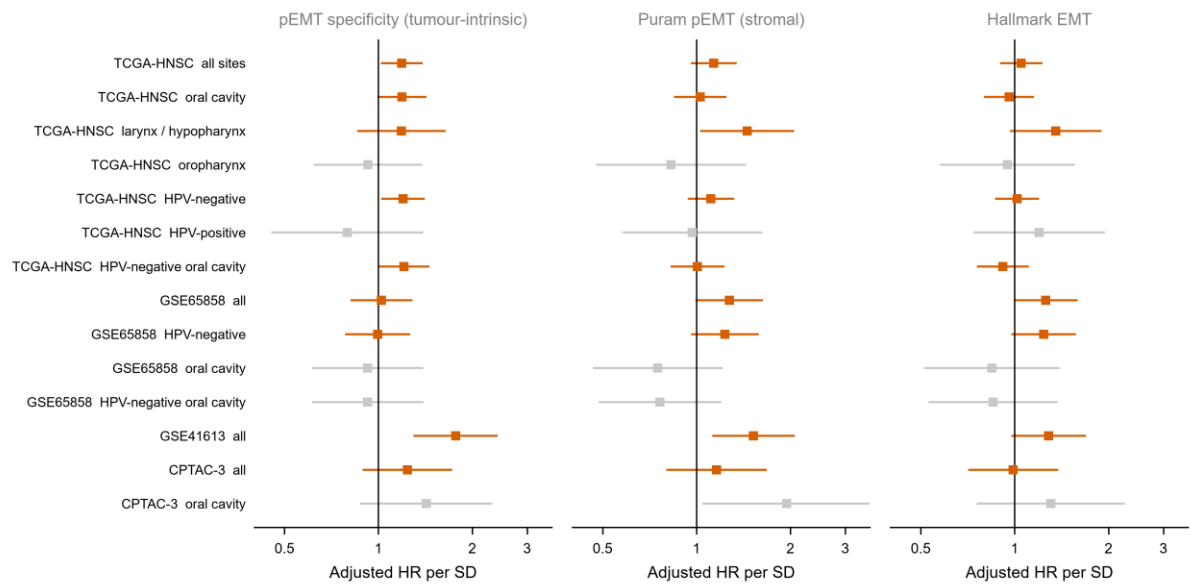

Grey: fewer than 30 events, underpowered. Each score is standardised within the subgroup, so every estimate is per standard deviation of that subgroup.

**Supplementary Fig. 13. Association of pEMT specificity with clinical and pathological variables in TCGA-HNSC.** Panels show pEMT specificity by pathological N stage, T stage, HPV status, anatomical site, histological grade, lymphovascular invasion, perineural invasion and extranodal extension. Boxes show the median and interquartile range and whiskers extend to 1.5 times the interquartile range. The test used is stated on each panel.

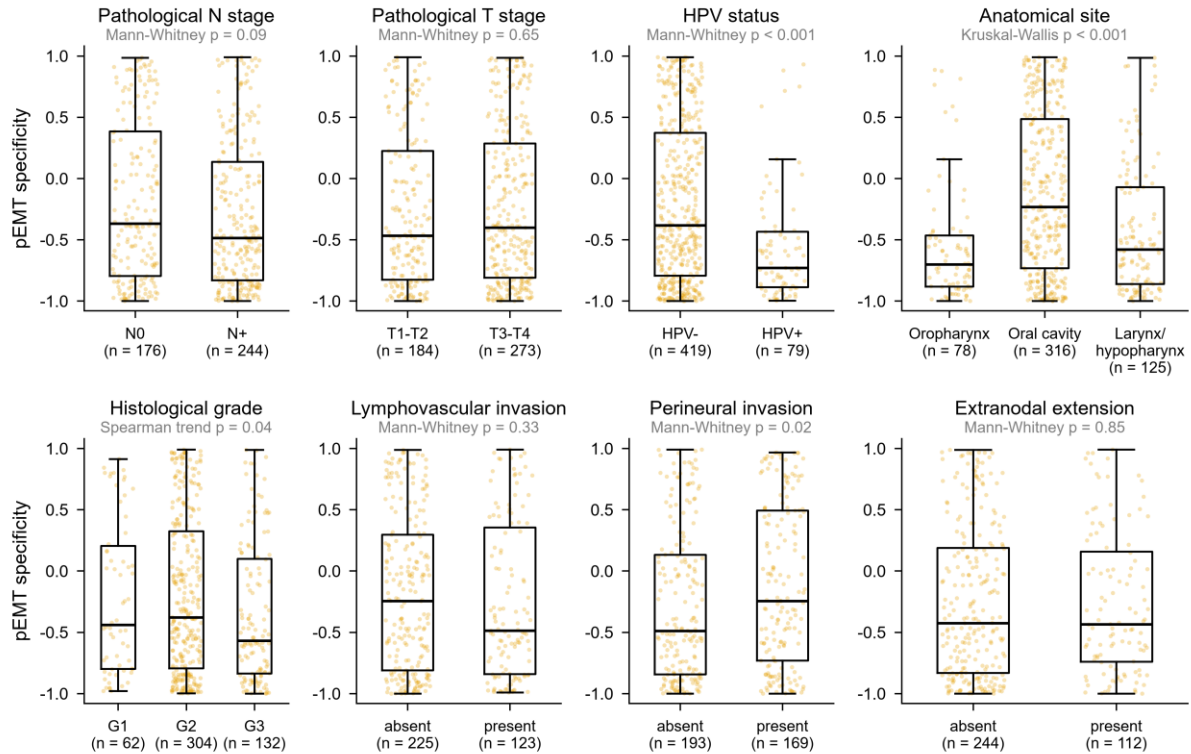

**Supplementary Fig. 14. Reversal of the classifier signature by L1000 perturbations. a** Compounds ranked by the number of L1000 signatures that oppose the query signature, with the number that mimic it stated beside each bar. Dark orange marks a compound significant by a one-sided sign test for excess reversal at a false discovery rate of 0.05 and light orange one that is not. **b** Share of reversing signatures contributed by each mechanism-of-action class, in orange, against the mean share across five random gene sets of matched size, in grey. The comparison with 50 random gene sets is in Supplementary Data 43.

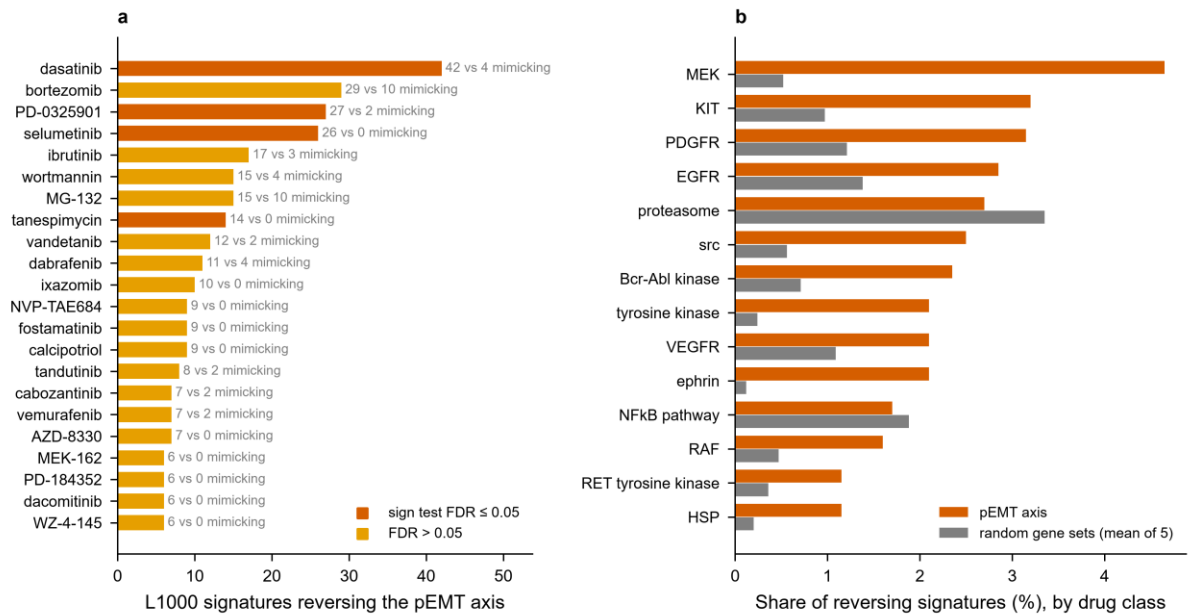

**Supplementary Fig. 15. All mechanism-of-action classes against the permutation null.** The full set of classes with at least five compounds, in HNSCC and pan-squamous models, plotted as in Supplementary Fig. 16c. Grey bars show the central 95% of the permutation null. No class reaches a false discovery rate of 0.05 in either the HNSCC or the pan-squamous models.

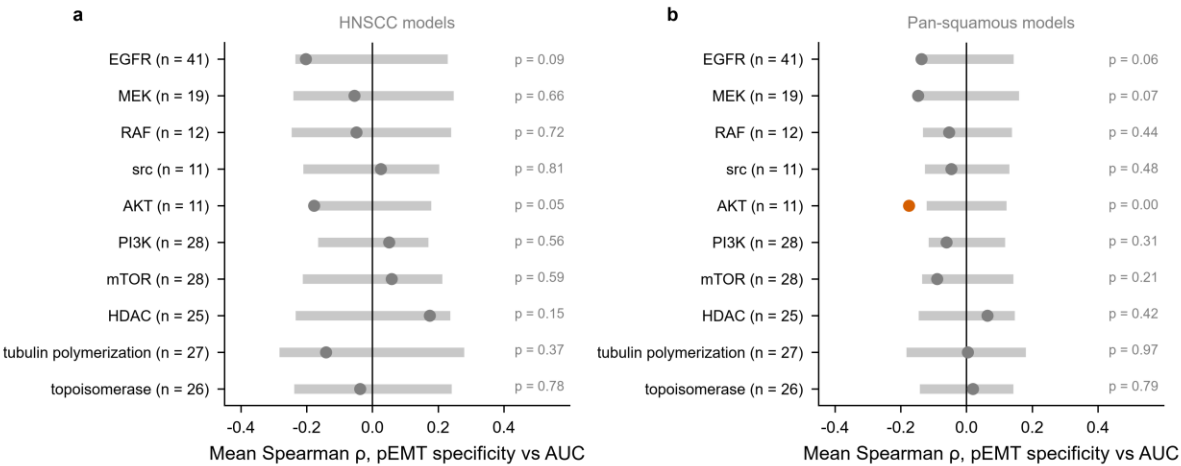

grey bar: 95% of the permutation null (phenotype labels shuffled across cell lines, 5,000 permutations). Negative $\rho$: more sensitive in pEMT-high models.

**Supplementary Fig. 16. Sensitivity and dependency in cell line models.** **a** PRISM secondary screen: Spearman correlation between pEMT specificity and area under the dose-response curve for each of 1,444 compounds across 23 HNSCC models, against the nominal p-value. Negative correlation indicates greater sensitivity in pEMT-high models. Orange marks compounds nominated by the network proximity analysis. **b** CRISPR: the same comparison for Chronos gene effect across 17,631 genes and 63 models, with the two module gene sets highlighted. **c** Mechanism-of-action classes tested against a permutation null over cell lines. Open circles are HNSCC models and open triangles the wider pan-squamous set, each showing the observed mean Spearman correlation for that class. Grey bars show the central 95% of the permutation null from 5,000 permutations of the phenotype labels across cell lines, drawn as a normal approximation from the mean and standard deviation of the null. A filled orange marker indicates a permutation p-value at or below 0.05. The eight classes named in the text are shown here and all classes with at least five compounds are in Supplementary Fig. 15.

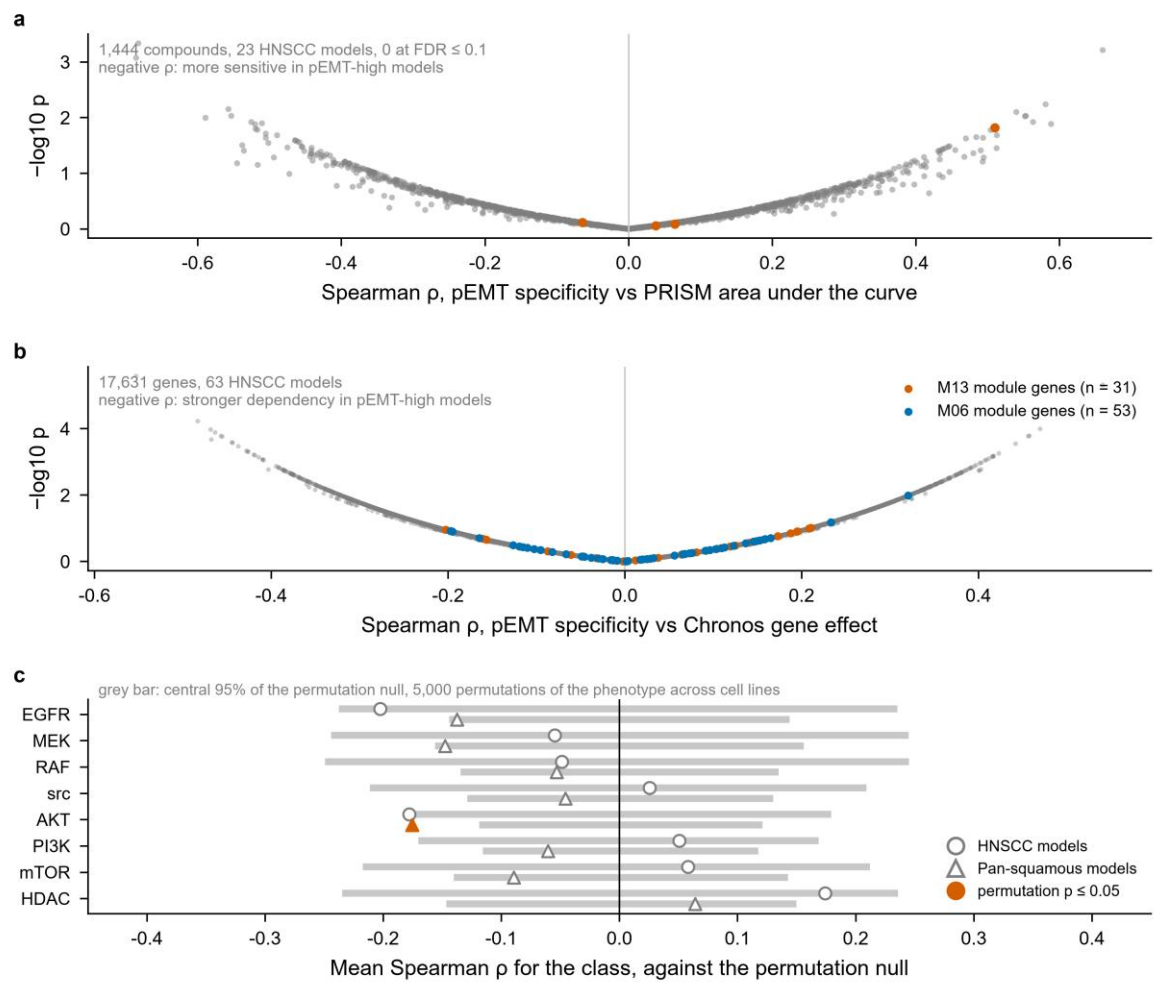

**Supplementary Fig. 17. Network proximity of approved drugs to each seed gene set. a** Canonical pEMT genes within module M06. **b** Module M13 hubs. **c** The genes with the 100 largest positive pEMT-high classifier coefficients. In each panel, approved drugs are ranked by proximity z-score, with the dashed vertical line at the significance threshold of $-1.96$ and the solid line at zero. Dark orange marks a hit that remains significant when any single seed gene is removed, light orange a hit that does not, and grey a drug that is not a hit. The horizontal bar extending from each hit shows its z-score after removal of the most influential seed gene. **d** For the classifier seed set, the number of approved hits attributable to each seed gene, split by whether the hit survives seed removal. A stricter variant, in which a drug's targets that are themselves seed genes are removed before the distance is computed, is reported in Supplementary Data 55.

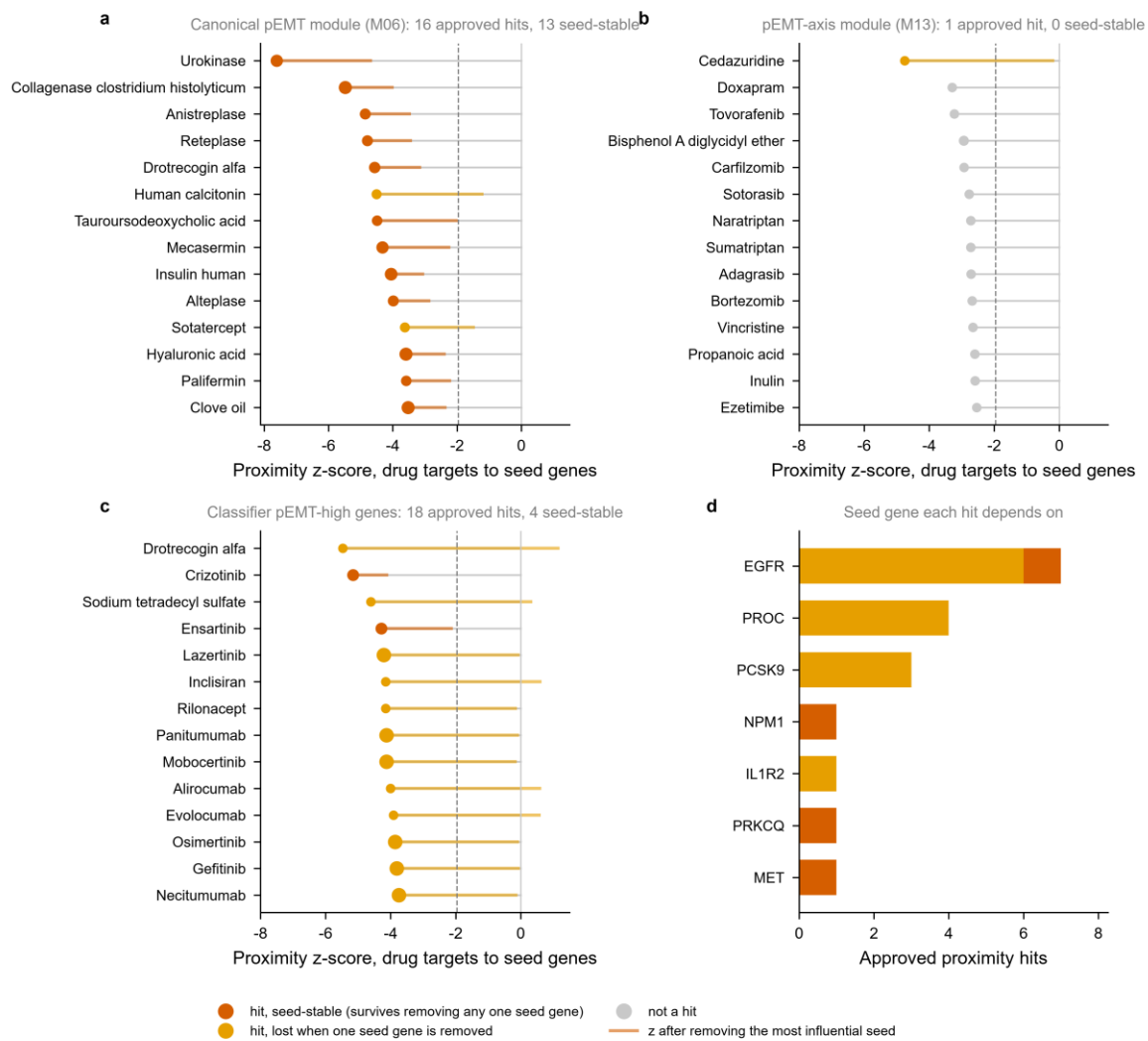

**Supplementary Fig. 18. Agreement between the two drug-target definitions.** Proximity z-score computed with STITCH association targets against the same score computed with DrugBank curated targets, for each seed gene set. Orange marks approved drugs called as hits under both definitions. Dashed lines mark the significance threshold of $-1.96$ on each axis.

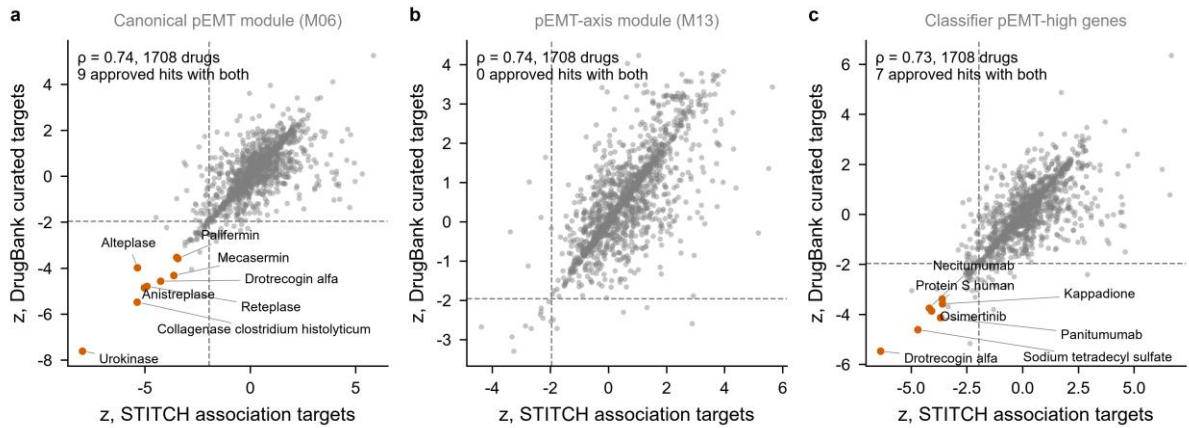

### Supplementary Data

Each Supplementary Data item is an Excel workbook. Its parts are separate sheets, and the first row of each sheet repeats the legend given here.

#### **Supplementary Data 1. Each of the 100 Puram pEMT genes resolved by cell type in GSE103322.**

Supplementary Data 1 (sheet SD1\_pemt\_gene\_partition). Each of the 100 Puram pEMT genes resolved by cell type in GSE103322. Columns give the share of total expression (linear scale) and detection rate in malignant cells, fibroblasts, endothelium and immune cells, the arm the gene is assigned to, its gene family, the log2 fold change between pEMT-high and epithelial-like malignant cells, and whether it is among the classifier's top 100 positive coefficients.

#### **Supplementary Data 2. Each Puram pEMT gene in GSE103322, GSE150321 and GSE181919: malignant share of total expression and the composition-independent malignant share of mean expression per cell, detection rates, and arm by the per-cell measure.**

Supplementary Data 2 (sheet SD2\_partition\_replication). Each Puram pEMT gene in GSE103322, GSE150321 and GSE181919: malignant share of total expression and the composition-independent malignant share of mean expression per cell, detection rates, and arm by the per-cell measure.

Supplementary Data 2b (sheet SD2b\_partition\_repl\_summary). Per dataset: arm counts by the per-cell measure, agreement and kappa with the published total-share assignment, correlation with the published shares, and the hemidesmosome versus other comparison.

Supplementary Data 2c (sheet SD2c\_partition\_repl\_cross). Agreement of the per-cell measure between each pair of GSE103322, GSE150321 and GSE181919, and the genes on the same side of parity in all three.

Supplementary Data 2d (sheet SD2d\_partition\_GSE181919\_pts). GSE181919 per-patient analysis: for each pEMT gene, the number of patients with at least 20 malignant cells and 20 fibroblasts in tumour tissue, the median per-cell malignant share across them, and the fraction of patients in which the gene is malignant-predominant.

#### **Supplementary Data 3. Gene sets used to score the two arms of the pEMT partition directly.**

Supplementary Data 3 (sheet SD3\_arm\_gene\_sets). Gene sets used to score the two arms of the pEMT partition directly. They are the total-share arms (52 and 43 genes), the cores (12

genes with malignant share at least 80% and 14 with fibroblast share at least 60%), the consensus arms (16 and 58 genes on the same side of parity by mean expression per cell in GSE103322, GSE150321 and GSE181919, Supplementary Data 2), and sensitivity sets with the Zhou genes removed or alternative core thresholds.

Supplementary Data 3b (sheet SD3b\_arm\_tcga\_correlations). Spearman correlation of each direct arm score with pEMT specificity, the published scores and the M06 and M13 module eigengenes in the 520 TCGA-HNSC primaries.

Supplementary Data 3c (sheet SD3c\_arm\_GSE65021). Direct arm scores in GSE65021: bootstrap AUC, rank-biserial correlation, two-sided Mann-Whitney p, Firth odds ratio per SD with profile-likelihood interval, and correlation with pEMT specificity and the Puram signature.

Supplementary Data 3d (sheet SD3d\_arm\_GSE65021\_joint). Joint Firth models in GSE65021: malignant and stromal arm together, and the malignant arm with each epithelial score.

Supplementary Data 3e (sheet SD3e\_arm\_survival\_pooled). Hazard ratio per SD for each direct arm score and comparators, univariable and adjusted, pooled over TCGA-HNSC, GSE41613, CPTAC-3 and GSE65858 by random and fixed effects.

Supplementary Data 3f (sheet SD3f\_arm\_survival\_cohorts). Per-cohort Cox estimates underlying Supplementary Data 3e.

Supplementary Data 3g (sheet SD3g\_tcga\_systemic\_therapy). TCGA-HNSC adjusted hazard ratios after excluding patients whose legacy clinical record marks systemic (targeted molecular) therapy, and restricted to those recorded as not receiving it.

Supplementary Data 3h (sheet SD3h\_arm\_definition\_timeline). When each arm definition and each other score analysed in GSE65021 was fixed, relative to the first analysis of that cohort (20 Sep 2026, pEMT specificity and the published scores). Dates are from the project change log. All gene sets come from single-cell data only. The xenograft panels were scored after every definition here was fixed.

**Supplementary Data 4. WGCNA modules (signed, soft power 13, 8,000 most variable genes, 520 primaries): size, correlation of the module eigengene with the classifier outputs and Hallmark EMT, overlap with the Puram pEMT signature (hypergeometric p), best-matching program, top hubs by kME.**

Supplementary Data 4 (sheet SD4\_wgcna\_modules). WGCNA modules (signed, soft power 13, 8,000 most variable genes, 520 primaries): size, correlation of the module eigengene with the classifier outputs and Hallmark EMT, overlap with the Puram pEMT signature (hypergeometric p), best-matching program, top hubs by kME.

Supplementary Data 4b (sheet SD4b\_module\_trait). Module eigengene to trait correlations (Pearson r and p), primaries only.

Supplementary Data 4c (sheet SD4c\_module\_enrichment). Hypergeometric enrichment of each module for the Puram programs, Hallmark EMT and immune/stromal marker sets.

Supplementary Data 4d (sheet SD4d\_gene\_modules). Module assignment for every gene in the WGCNA input (core\_member\_pre\_PAM = assigned before the PAM step) and kME to every module eigengene.

Supplementary Data 4e (sheet SD4e\_network\_seeds). Seed gene sets for the network analysis. pEMT\_axis is the M13 members with kME $\geq 0.7$, canonical\_pEMT the Puram pEMT genes in M06, and classifier\_pEMT\_high the top 100 positive pEMT-high classifier coefficients.

**Supplementary Data 5. Classifier probabilities for every malignant (patient x site) and fibroblast (patient) pseudobulk with $\geq 20$ cells, after within-dataset z-scoring of the classifier genes.**

Supplementary Data 5 (sheet SD5\_sc\_pseudobulk). Classifier probabilities for every malignant (patient x site) and fibroblast (patient) pseudobulk with $\geq 20$ cells, after within-dataset z-scoring of the classifier genes. puram\_pemt\_signature = mean z of the Puram pEMT genes.

Supplementary Data 5b (sheet SD5b\_puram\_nodal\_status). Nodal status in the single-cell discovery data (GSE103322): primary-site malignant pseudobulk scores of node-positive against node-negative patients, all of them training data. p\_one\_sided tests $N^+ > N_0$, the direction Puram et al. reported. p\_two\_sided\_exact is the exact two-sided Mann-Whitney p-value. The groups separate in the opposite direction ($U = 0$).

Supplementary Data 5c (sheet SD5c\_puram\_ln\_vs\_primary). Single-cell validation: puram\_In\_vs\_primary\_paired.tsv

Supplementary Data 5d (sheet SD5d\_sc\_summary\_by\_cell\_type). Single-cell validation: summary\_by\_cell\_type.tsv

Supplementary Data 5e (sheet SD5e\_sc\_rank\_agreement). Single-cell validation: rank\_agreement\_with\_puram\_signature.tsv

Supplementary Data 5f (sheet SD5f\_per\_cell\_score\_correlation). Single-cell validation: per\_cell\_score\_correlations.tsv

**Supplementary Data 6. Alternative classifiers on the identical pseudobulk training set and leave-one-dataset-out folds: accuracy, AUROC per class, log loss, and Spearman correlation of each model's P(pEMT-high) with the elastic net on the held-out fold.**

Supplementary Data 6 (sheet SD6\_classifier\_benchmark). Alternative classifiers on the identical pseudobulk training set and leave-one-dataset-out folds: accuracy, AUROC per class, log loss, and Spearman correlation of each model's P(pEMT-high) with the elastic net on the held-out fold.

Supplementary Data 6b (sheet SD6b\_classifier\_benchmark\_mean). Fold means of Supplementary Data 6\_classifier\_benchmark.

**Supplementary Data 7. TCGA-HNSC samples: classifier probabilities, pEMT specificity, the EMT score panel (76GS, KS, Hallmark EMT, Puram pEMT, Puram epithelial differentiation, MLR), HPV status and clinical variables used in the models.**

Supplementary Data 7 (sheet SD7\_tcga\_scores). TCGA-HNSC samples: classifier probabilities, pEMT specificity, the EMT score panel (76GS, KS, Hallmark EMT, Puram pEMT, Puram epithelial differentiation, MLR), HPV status and clinical variables used in the models.

**Supplementary Data 8. Saturation of pEMT specificity at plus or minus 1 and of its log-odds summary in each cohort.**

Supplementary Data 8 (sheet SD8\_saturation\_tails). Saturation of pEMT specificity at plus or minus 1 and of its log-odds summary in each cohort.

Supplementary Data 8b (sheet SD8b\_saturation\_gse65021). GSE65021: pEMT specificity and its log-odds summaries against long progression-free survival.

Supplementary Data 8c (sheet SD8c\_saturation\_survival). Pooled hazard ratio per SD of pEMT specificity and its log-odds summaries across the four survival cohorts, DerSimonian-Laird and Hartung-Knapp, with each cohort left out in turn.

Supplementary Data 8d (sheet SD8d\_saturation\_xenografts). Xenograft panels: pooled Firth odds ratio of pEMT specificity and its log-odds summaries.

Supplementary Data 8e (sheet SD8e\_saturation\_correlations). Spearman correlation of each summary with pEMT specificity and the malignant core in each cohort.

**Supplementary Data 9. Composition of each Leiden cluster obtained from malignant cells after removing all 533 genes of the six Puram programmes, with labels used at no point in the clustering.**

Supplementary Data 9 (sheet SD9\_label\_free\_clusters). Composition of each Leiden cluster obtained from malignant cells after removing all 533 genes of the six Puram programmes, with labels used at no point in the clustering.

Supplementary Data 9b (sheet SD9b\_label\_free\_summary). Summary statistics for the label-free recovery: genes removed and retained, cluster count, adjusted Rand index and its permutation p-value, adjusted mutual information, silhouette against its null, leave-one-patient-out AUROC on non-Puram genes, and the number of Puram genes among the top 100 classifier coefficients.

**Supplementary Data 10. Each gene of the Tyler and Tirosh style EMT signature set resolved to the cancer or fibroblast compartment in GSE103322 by share of expression.**

Supplementary Data 10 (sheet SD10\_tt\_compartments). Each gene of the Tyler and Tirosh style EMT signature set resolved to the cancer or fibroblast compartment in GSE103322 by share of expression.

**Supplementary Data 11. Spearman correlation matrix among the EMT score panel, the malignant arm, and the cancer and fibroblast compartment scores obtained by applying the Tyler and Tirosh decoupling principle to the 520 TCGA-HNSC primaries.**

Supplementary Data 11 (sheet SD11\_tt\_correlations). Spearman correlation matrix among the EMT score panel, the malignant arm, and the cancer and fibroblast compartment scores obtained by applying the Tyler and Tirosh decoupling principle to the 520 TCGA-HNSC primaries.

**Supplementary Data 12. TCGA-HNSC: Spearman correlation of each of the 41 Gavish et al.**

Supplementary Data 12 (sheet SD12\_gavish\_mp\_tcga). TCGA-HNSC: Spearman correlation of each of the 41 Gavish et al. 2023 cancer meta-programmes with our scores and the published signatures.

Supplementary Data 12b (sheet SD12b\_gavish\_mp\_overlap). Gene overlap (hypergeometric, 23,436-gene classifier universe) of each meta-programme with the malignant core, the stromal core, the classifier's top 100 pEMT-high coefficients and the Puram pEMT genes.

Supplementary Data 12c (sheet SD12c\_gavish\_mp\_gse65021). GSE65021: each meta-programme against long progression-free survival, AUC with bootstrap interval, two-sided Mann-Whitney p and Benjamini-Hochberg q over the 41 programmes.

Supplementary Data 12d (sheet SD12d\_gavish\_mp\_joint). GSE65021: selected meta-programmes entered with the malignant core in one Firth model.

**Supplementary Data 13. GSE103322: cell of origin of two published partial EMT gene sets and their companion sets, the mammary EMT clusters of Knutsen et al.**

Supplementary Data 13 (sheet SD13\_ext\_pemt\_cell\_of\_origin). GSE103322: cell of origin of two published partial EMT gene sets and their companion sets, the mammary EMT clusters of Knutsen et al. 2023 as tabulated by Sahoo et al. 2024 (EMT\_down, EMT\_partial, EMT\_up) and the spatial HNSCC programmes of Simkin et al. 2026 (pEMT, Fibroblast, Epithelial). For each set, genes detected, median share of total expression from malignant cells and from fibroblasts, and genes with a larger malignant than fibroblast share.

Supplementary Data 13b (sheet SD13b\_ext\_pemt\_tcga). TCGA-HNSC primaries: Spearman correlation (and p) of each external set, scored as the mean per-gene z-score, with the TCGA 2015 Basal centroid score, the malignant and stromal cores and pEMT specificity.

Supplementary Data 13c (sheet SD13c\_ext\_pemt\_basal\_steiger). TCGA-HNSC primaries: comparison of the correlation of the Basal centroid score with a partial EMT set and with its full EMT or fibroblast counterpart, Steiger test for dependent correlations.

Supplementary Data 13d (sheet SD13d\_ext\_pemt\_gse65021). GSE65021: each external set against long progression-free survival, with the AUC and its 2,000-resample bootstrap interval, the two-sided Mann-Whitney p and the Spearman correlation with the Basal score and the two cores. Sets with $p < 0.05$ were entered with the Basal centroid score in a joint Firth model (odds ratios per SD). The cores and the Basal score are shown for reference.

Supplementary Data 13e (sheet SD13e\_ext\_pemt\_gene\_sets). The external gene sets as scored: Knutsen et al. 2023 mammary EMT clusters (Sahoo et al. 2024, Table Supplementary Data 1, one Ensembl-only entry dropped) and Simkin et al. 2026 spatial meta-programmes (figshare file Final\_Metaprograms\_Extended.csv).

**Supplementary Data 14. Simulated tumours built from GSE181919 cells: standardised regression of each score on the share of pEMT-high malignant cells (q), the fibroblast fraction (f) and the immune fraction (u), with the partial R squared of each factor.**

Supplementary Data 14 (sheet SD14\_composition\_simulation). Simulated tumours built from GSE181919 cells: standardised regression of each score on the share of pEMT-high malignant cells (q), the fibroblast fraction (f) and the immune fraction (u), with the partial R squared of each factor.

Supplementary Data 14b (sheet SD14b\_simulated\_tumours). Design and scores of the 450 simulated tumours.

**Supplementary Data 15. BayesPrism deconvolution with GSE181919 as the reference: Spearman correlations of the estimated cell fractions and of gene-set scores on deconvolved malignant-cell and fibroblast expression with the bulk scores, in TCGA-HNSC (520 primaries) and GSE65021 (40 patients, microarray).**

Supplementary Data 15 (sheet SD15\_bayesprism\_correlations). BayesPrism deconvolution with GSE181919 as the reference: Spearman correlations of the estimated cell fractions and of gene-set scores on deconvolved malignant-cell and fibroblast expression with the bulk scores, in TCGA-HNSC (520 primaries) and GSE65021 (40 patients, microarray).

Supplementary Data 15b (sheet SD15b\_bayesprism\_gse65021\_pfs). GSE65021: deconvolved scores and cell fractions against long progression-free survival, AUC with 2,000-resample bootstrap interval and two-sided Mann-Whitney p, in all 40 patients and in the 31 primary-tumour specimens, with the bulk scores alongside.

Supplementary Data 15c (sheet SD15c\_bayesprism\_tcga\_subtype). TCGA-HNSC: deconvolved scores and cell fractions by TCGA 2015 expression subtype (279 primaries), Kruskal-Wallis and Basal against the rest.

Supplementary Data 15d (sheet SD15d\_bayesprism\_fractions). Summary of the estimated cell fractions per cohort. TCGA fractions are from the initial Gibbs step.

Supplementary Data 15e (sheet SD15e\_bayesprism\_reference). Composition of the GSE181919 single-cell reference: cells available and used per cell type and state.

**Supplementary Data 16. Spearman correlations among the EMT scores in TCGA-HNSC primaries.**

Supplementary Data 16 (sheet SD16\_emt\_panel\_correlation). Spearman correlations among the EMT scores in TCGA-HNSC primaries.

Supplementary Data 16b (sheet SD16b\_emt\_panel\_cox). EMT score panel: hazard ratio per SD, univariable and adjusted (stage, age, site, HPV), and cross-validated C-index.

Supplementary Data 16c (sheet SD16c\_emt\_panel\_head\_to\_head). Head-to-head: pEMT specificity and each score in the same adjusted Cox model (stage, age, site, HPV). Columns give the HR per SD of each given the other, the likelihood-ratio p for adding either to clinical covariates plus the other, and the cross-validated C-index of clinical + score with and without pEMT specificity.

**Supplementary Data 17. Faithful MLR-EMT in each cohort: cohort normalisation offset, epithelial, hybrid and mesenchymal state counts, and correlation with the approximation and with the other scores.**

Supplementary Data 17 (sheet SD17\_mlr\_versions\_cohorts). Faithful MLR-EMT in each cohort: cohort normalisation offset, epithelial, hybrid and mesenchymal state counts, and correlation with the approximation and with the other scores.

Supplementary Data 17b (sheet SD17b\_mlr\_versions\_GSE65021). MLR-EMT in GSE65021 computed as the earlier approximation and as the faithful implementation of George et al. (Jolly-group reference code): AUC, rank-biserial, odds ratios and correlations.

**Supplementary Data 18. MLR-EMT read as a three-state model.**

Supplementary Data 18 (sheet SD18\_mlr\_emt\_states). MLR-EMT read as a three-state model. For GSE65021 and TCGA-HNSC, the range of each state probability and of the score. For GSE65021, the AUC for prolonged progression-free survival, two-sided Mann-Whitney p and Firth odds ratio per SD. Spearman correlation with pEMT specificity, the malignant core, 76GS and the Puram pEMT signature.

Supplementary Data 18b (sheet SD18b\_mlr\_emt\_state\_counts). GSE65021 patients by most probable MLR-EMT state and progression-free survival group, with the Fisher test of hybrid against mesenchymal.

**Supplementary Data 19. Per-gene Spearman correlation with pEMT specificity in TCGA-HNSC primaries, grouped into hemidesmosome/laminin-332, EGFR ligands and receptor, basal keratinocyte, classical mesenchymal EMT, and fibroblast/ECM stroma.**

Supplementary Data 19 (sheet SD19\_gene\_level). Per-gene Spearman correlation with pEMT specificity in TCGA-HNSC primaries, grouped into hemidesmosome/laminin-332, EGFR ligands and receptor, basal keratinocyte, classical mesenchymal EMT, and fibroblast/ECM stroma.

Supplementary Data 19b (sheet SD19b\_coefficient\_overlap). Hypergeometric overlap of the top 100 positive pEMT-high classifier coefficients with each published gene set, against the 23,409-gene classifier universe. The classifier was trained only on single-cell pseudobulk labels, without the EGF-stimulation, cetuximab-outcome or tumour-budding data behind those sets.

Supplementary Data 19c (sheet SD19c\_published\_correlations). Spearman correlation of five externally published HNSCC programmes (the 171-gene EGFR-induced EMT signature of Schinke 2022, three sets from Zhou 2025 with 46 EGFR invasion fDEGs, a 59-gene invasive network and 9 cetuximab-PFS predictors, and the 28-gene tumour budding signature of Ourailidis 2026) with every score in the panel, in TCGA-HNSC and both external cohorts. Gene lists taken from the source papers' supplementary files.

Supplementary Data 19d (sheet SD19d\_tcga\_subtype). Each score by TCGA 2015 four-class expression subtype (Basal, Classical, Atypical, Mesenchymal, n = 279 matched primaries), with Kruskal-Wallis across subtypes and Basal-vs-rest and Mesenchymal-vs-rest Mann-Whitney.

Supplementary Data 19e (sheet SD19e\_joint\_cox). Adjusted Cox (stage, age, site, HPV) with pEMT specificity and one published signature in the same model, per SD, with likelihood-ratio tests for adding either to the other.

**Supplementary Data 20. Ligand-receptor pairs from fibroblasts to each malignant state by LIANA consensus of five methods, with magnitude and specificity ranks for pEMT-high and epithelial-like targets and the difference in log<sub>10</sub> specificity between them.**

Supplementary Data 20 (sheet SD20\_caf\_lr\_pairs). Ligand-receptor pairs from fibroblasts to each malignant state by LIANA consensus of five methods, with magnitude and specificity ranks for pEMT-high and epithelial-like targets and the difference in log<sub>10</sub> specificity between them.

Supplementary Data 20b (sheet SD20b\_caf\_lr\_matched). The same comparison repeated five times with pEMT-high cells downsampled to the epithelial-like cell count, giving the per-seed difference, its mean, and the number of runs favouring pEMT-high.

Supplementary Data 20c (sheet SD20c\_ligand\_source). Share of total expression and detection rate per cell group for EGFR-family ligands and receptors and for canonical fibroblast-derived ligands, used to test autocrine against paracrine supply.

**Supplementary Data 21. Each score by the consensus expression clusters GSE65858 publishes for itself.**

Supplementary Data 21 (sheet SD21\_gse65858\_subtype). Each score by the consensus expression clusters GSE65858 publishes for itself. These clusters share their names with the TCGA four-class calls but were derived de novo within that cohort, and the externally defined Zhou cetuximab-PFS gene set peaks in a different class here than in TCGA.

**Supplementary Data 22. Per-patient scores and recorded covariates for the 40 patients of GSE65021 treated with platinum plus cetuximab, including progression-free survival**

**group, age, stage, grade, prior radiotherapy, primary site and whether the specimen was the primary or the recurrence.**

Supplementary Data 22 (sheet SD22\_GSE65021\_scores). Per-patient scores and recorded covariates for the 40 patients of GSE65021 treated with platinum plus cetuximab, including progression-free survival group, age, stage, grade, prior radiotherapy, primary site and whether the specimen was the primary or the recurrence.

**Supplementary Data 23. Association of every score with prolonged progression-free survival on cetuximab: medians by group, rank-biserial correlation, one- and two-sided Mann-Whitney p values with Benjamini-Hochberg q values, bootstrap ROC AUC, and maximum-likelihood odds ratios per standard deviation unadjusted and adjusted (Firth estimates are in Supplementary Data 24).**

Supplementary Data 23 (sheet SD23\_GSE65021\_association). Association of every score with prolonged progression-free survival on cetuximab: medians by group, rank-biserial correlation, one- and two-sided Mann-Whitney p values with Benjamini-Hochberg q values, bootstrap ROC AUC, and maximum-likelihood odds ratios per standard deviation unadjusted and adjusted (Firth estimates are in Supplementary Data 24). MLR-EMT is the faithful implementation of George et al. The Zhou nine-gene set is flagged as not independent of this cohort and excluded from the FDR.

**Supplementary Data 24. Firth penalised logistic regression in GSE65021, each score alone, unadjusted and adjusted: odds ratio of prolonged progression-free survival per SD, profile penalised-likelihood interval and penalised likelihood-ratio p.**

Supplementary Data 24 (sheet SD24\_GSE65021\_firth). Firth penalised logistic regression in GSE65021, each score alone, unadjusted and adjusted: odds ratio of prolonged progression-free survival per SD, profile penalised-likelihood interval and penalised likelihood-ratio p.

Supplementary Data 24b (sheet SD24b\_GSE65021\_core\_vs\_canonica). GSE65021: the malignant core and, separately, pEMT specificity entered together with the canonical Puram pEMT signature in one Firth-penalised logistic regression of long progression-free survival, both terms standardised within the cohort. rho\_between\_terms is the Spearman correlation of the two terms. The terms are collinear, so the estimates are imprecise.

Supplementary Data 24c (sheet SD24c\_GSE65021\_firth\_h2h). Firth head-to-head models, pEMT specificity with each other score.

Supplementary Data 24d (sheet SD24d\_GSE65021\_firth\_epi). Firth models entering MLR-EMT or KS with 76GS or Puram epithelial differentiation.

**Supplementary Data 25. One multiplicity family in GSE65021: every independently derived published score and the four arm scores specified before the cohort was examined, with AUC, two-sided Mann-Whitney p and Benjamini-Hochberg q.**

Supplementary Data 25 (sheet SD25\_GSE65021\_fdr\_family). One multiplicity family in GSE65021: every independently derived published score and the four arm scores specified before the cohort was examined, with AUC, two-sided Mann-Whitney p and Benjamini-Hochberg q.

**Supplementary Data 26. Transportability of pEMT specificity and the malignant core in GSE65021: AUC with within-cohort, leave-one-out and external-reference (GSE65858) standardisation, and agreement with the published within-cohort score.**

Supplementary Data 26 (sheet SD26\_GSE65021\_transport). Transportability of pEMT specificity and the malignant core in GSE65021: AUC with within-cohort, leave-one-out and external-reference (GSE65858) standardisation, and agreement with the published within-cohort score.

**Supplementary Data 27. Firth odds ratio per SD in GSE65021 for the malignant core and pEMT specificity, alone and adjusted for the stromal core, an 18-gene immune score, or both, with their correlations with these covariates.**

Supplementary Data 27 (sheet SD27\_GSE65021\_purity). Firth odds ratio per SD in GSE65021 for the malignant core and pEMT specificity, alone and adjusted for the stromal core, an 18-gene immune score, or both, with their correlations with these covariates.

**Supplementary Data 28. Paired bootstrap comparison (2,000 resamples) of the AUC of each primary measure with 76GS, Puram epithelial differentiation, the canonical Puram pEMT signature and the Zhou nine-gene set.**

Supplementary Data 28 (sheet SD28\_GSE65021\_paired\_auc). Paired bootstrap comparison (2,000 resamples) of the AUC of each primary measure with 76GS, Puram epithelial differentiation, the canonical Puram pEMT signature and the Zhou nine-gene set.

**Supplementary Data 29. Exploratory combined marker in GSE65021, specified after the single-score results were known.**

Supplementary Data 29 (sheet SD29\_GSE65021\_combined). Exploratory combined marker in GSE65021, specified after the single-score results were known. It is pEMT specificity plus

Puram epithelial differentiation, as an equal-weight sum of standardised scores, with 76GS and sign-reversed MLR-EMT as sensitivity partners. Columns give the AUC with 2,000-resample bootstrap interval, the paired bootstrap gain over pEMT specificity alone (gain\_over\_axis), leave-one-out AUC of a fitted two-term logistic model, and AUC in the 31 primary-tumour specimens.

**Supplementary Data 30. GSE65021: Basal centroid score (correlation with the TCGA 2015 Basal centroid, 674 genes), with and without the pEMT and Zhou genes, against long progression-free survival: AUC with bootstrap interval, two-sided Mann-Whitney p, Firth odds ratio per SD, and Spearman correlations with our scores.**

Supplementary Data 30 (sheet SD30\_GSE65021\_basal\_centroid). GSE65021: Basal centroid score (correlation with the TCGA 2015 Basal centroid, 674 genes), with and without the pEMT and Zhou genes, against long progression-free survival: AUC with bootstrap interval, two-sided Mann-Whitney p, Firth odds ratio per SD, and Spearman correlations with our scores.

Supplementary Data 30b (sheet SD30b\_GSE65021\_subtype\_calls). GSE65021: centroid subtype calls by progression-free survival group.

Supplementary Data 30c (sheet SD30c\_GSE65021\_basal\_joint). GSE65021: the malignant core and, separately, pEMT specificity entered with the Basal centroid score in one Firth model of long progression-free survival.

Supplementary Data 30d (sheet SD30d\_GSE65021\_basal\_paired\_auc). GSE65021: paired bootstrap difference in AUC of each primary measure against the Basal centroid score (2,000 resamples of patients).

Supplementary Data 30e (sheet SD30e\_TCGA\_basal\_score). TCGA-HNSC: Basal centroid score against pEMT specificity and the cores, and agreement of the centroid calls with the published subtype calls.

**Supplementary Data 31. GSE65021: the Basal centroid score and the malignant core scored for each patient against four references (all 40 patients, the other 39 patients, GSE65858 on the same array family, and no reference), with the AUC for long progression-free survival, its 2,000-resample bootstrap interval and the Spearman correlation with the within-cohort score.**

Supplementary Data 31 (sheet SD31\_single\_patient\_scoring). GSE65021: the Basal centroid score and the malignant core scored for each patient against four references (all 40 patients, the other 39 patients, GSE65858 on the same array family, and no reference), with the AUC for long

progression-free survival, its 2,000-resample bootstrap interval and the Spearman correlation with the within-cohort score.

**Supplementary Data 32. Cetuximab response in two head and neck cancer PDX panels (GSE84713 per patient and over all models, GSE183881 over tested models): AUC with bootstrap interval, two-sided Mann-Whitney p and Firth odds ratio per SD for each score, and correlation with pEMT specificity.**

Supplementary Data 32 (sheet SD32\_pdx\_cetuximab). Cetuximab response in two head and neck cancer PDX panels (GSE84713 per patient and over all models, GSE183881 over tested models): AUC with bootstrap interval, two-sided Mann-Whitney p and Firth odds ratio per SD for each score, and correlation with pEMT specificity.

Supplementary Data 32b (sheet SD32b\_pdx\_pooled). Fixed-effect pooling of the two PDX panels: odds ratio per SD with 95% interval.

Supplementary Data 32c (sheet SD32c\_pdx\_scores\_GSE84713). Per-model scores, response label, patient and molecular subtype for the 28 GSE84713 models.

Supplementary Data 32d (sheet SD32d\_pdx\_continuous\_RTV). GSE84713 models also reported by Klinghammer et al. 2020: Spearman correlation of each score with cetuximab relative tumour volume transcribed from that paper's Table 1 (lower is better).

Supplementary Data 32e (sheet SD32e\_pdx\_paired\_auc). Paired bootstrap AUC differences in the two PDX panels: malignant core and pEMT specificity against 76GS, Puram epithelial differentiation and the canonical Puram signature.

Supplementary Data 32f (sheet SD32f\_pdx\_combined). The patient-cohort combination (pEMT specificity plus Puram epithelial differentiation, equal weights) evaluated in the two PDX panels without refitting.

**Supplementary Data 33. GSE84713 and GSE183881: the TCGA 2015 Basal centroid score, the malignant core, pEMT specificity, the stromal core and the Zhou nine-gene set against cetuximab response, with the AUC and its 2,000-resample bootstrap interval, the two-sided Mann-Whitney p, the Firth odds ratio per SD and the Spearman correlation with the Basal centroid score.**

Supplementary Data 33 (sheet SD33\_pdx\_basal\_association). GSE84713 and GSE183881: the TCGA 2015 Basal centroid score, the malignant core, pEMT specificity, the stromal core and the Zhou nine-gene set against cetuximab response, with the AUC and its 2,000-resample bootstrap interval, the two-sided Mann-Whitney p, the Firth odds ratio per SD and the Spearman correlation with the Basal centroid score. GSE84713 is analysed per patient.

Supplementary Data 33b (sheet SD33b\_pdx\_basal\_joint). GSE84713 and GSE183881: Firth logistic regression of cetuximab response on the malignant core and the Basal centroid score entered together, both standardised, with the Spearman correlation between them.

Supplementary Data 33c (sheet SD33c\_pdx\_basal\_paired\_auc). GSE84713 and GSE183881: the difference in AUC between the malignant core and the Basal centroid score, with its paired 2,000-resample bootstrap interval.

Supplementary Data 33d (sheet SD33d\_pdx\_basal\_pooled). GSE84713 and GSE183881 pooled: inverse-variance weighted Firth odds ratio per SD for each score.

Supplementary Data 33e (sheet SD33e\_pdx\_published\_subtype). GSE84713: cetuximab response by the molecular subtype reported by the depositors (BA basal, CL classical, MS mesenchymal), per patient.

**Supplementary Data 34. Cetuximab sensitivity of cell lines (GDSC1 drug 1114) against each score: Spearman correlation with the dose-response area and $\ln$ IC50 (negative = higher score in more sensitive lines), and AUC for the most sensitive quartile, in the head and neck panel and in a squamous panel.**

Supplementary Data 34 (sheet SD34\_gdsc\_cetuximab). Cetuximab sensitivity of cell lines (GDSC1 drug 1114) against each score: Spearman correlation with the dose-response area and $\ln$ IC50 (negative = higher score in more sensitive lines), and AUC for the most sensitive quartile, in the head and neck panel and in a squamous panel.

Supplementary Data 34b (sheet SD34b\_gdsc\_cetuximab\_scores). Per-line scores and cetuximab response for both cell-line panels.

**Supplementary Data 35. Spearman correlation of every score with the abundance of each protein in the five pre-specified sets across 108 CPTAC-3 tumours, with Benjamini-Hochberg q values.**

Supplementary Data 35 (sheet SD35\_cptac\_protein). Spearman correlation of every score with the abundance of each protein in the five pre-specified sets across 108 CPTAC-3 tumours, with Benjamini-Hochberg q values.

**Supplementary Data 36. Spearman correlation of pEMT specificity with each of 21,057 phosphosites quantified in at least 30 tumours, before and after regressing out the**

**abundance of the site's own parent protein, with q values and membership of the EGFR to MAPK cascade.**

Supplementary Data 36 (sheet SD36\_cptac\_phosphosites). Spearman correlation of pEMT specificity with each of 21,057 phosphosites quantified in at least 30 tumours, before and after regressing out the abundance of the site's own parent protein, with q values and membership of the EGFR to MAPK cascade.

**Supplementary Data 37. Set-level test of the EGFR to MAPK cascade phosphosites against a null in which the score is permuted across tumours and the whole statistic recomputed 2,000 times.**

Supplementary Data 37 (sheet SD37\_cptac\_pathway\_perm). Set-level test of the EGFR to MAPK cascade phosphosites against a null in which the score is permuted across tumours and the whole statistic recomputed 2,000 times.

**Supplementary Data 38. Cox proportional-hazards models for overall survival in TCGA-HNSC primaries (pEMT specificity per unit, site reference oropharynx).**

Supplementary Data 38 (sheet SD38\_cox\_models). Cox proportional-hazards models for overall survival in TCGA-HNSC primaries (pEMT specificity per unit, site reference oropharynx).

Supplementary Data 38b (sheet SD38b\_cv\_cindex). Repeated 5-fold cross-validated concordance index of the clinical model with and without pEMT specificity.

Supplementary Data 38c (sheet SD38c\_km\_logrank). Log-rank test across pEMT specificity tertiles (Kaplan-Meier, Figure 4b).

**Supplementary Data 39. Random-effects (DerSimonian-Laird) meta-analysis of the adjusted hazard ratio per SD across TCGA-HNSC, GSE41613, CPTAC-3 and GSE65858, for every score: pooled HR, 95% CI, Cochran's Q, I-squared and tau-squared.**

Supplementary Data 39 (sheet SD39\_survival\_meta). Random-effects (DerSimonian-Laird) meta-analysis of the adjusted hazard ratio per SD across TCGA-HNSC, GSE41613, CPTAC-3 and GSE65858, for every score: pooled HR, 95% CI, Cochran's Q, I-squared and tau-squared.

Supplementary Data 39b (sheet SD39b\_survival\_meta\_hk). Random-effects pooling of the per-cohort hazard ratios per SD (log HR and SE from Supplementary Data 3f and Supplementary Data 39c) across TCGA-HNSC, GSE41613, CPTAC-3 and GSE65858, univariable and adjusted, with every cohort left out in turn. HR uses the DerSimonian-Laird between-cohort variance. DL\_low, DL\_up and DL\_p are the DerSimonian-Laird interval and p-value. HK\_low,

HK\_up and HK\_p apply the Hartung-Knapp adjustment (t distribution with k - 1 degrees of freedom), which is more conservative with few cohorts.

Supplementary Data 39c (sheet SD39c\_meta\_inputs). The per-cohort adjusted hazard ratios, confidence intervals, sample sizes and event counts that entered the meta-analysis.

**Supplementary Data 40. Adjusted Cox hazard ratio per SD for every score within each anatomical site and HPV stratum of TCGA-HNSC.**

Supplementary Data 40 (sheet SD40\_subgroup\_tcga). Adjusted Cox hazard ratio per SD for every score within each anatomical site and HPV stratum of TCGA-HNSC. The score is standardised inside each subgroup so all rows are per SD and comparable. The survival association is not specific to a site or HPV stratum.

Supplementary Data 40b (sheet SD40b\_subgroup\_external). The same subgroup test inside GSE65858, GSE41613 and CPTAC-3.

**Supplementary Data 41. CPTAC-3 HNSCC (Huang et al.**

Supplementary Data 41 (sheet SD41\_cptac\_cox). CPTAC-3 HNSCC (Huang et al. 2021, HPV-negative, surgically treated): univariable and adjusted Cox hazard ratio per SD for every score. Survival is taken from the current GDC clinical release (40 deaths), not the frozen 2021 publication snapshot (16 deaths).

**Supplementary Data 42. Association of each score with pathological N and T stage, HPV status, site, and (legacy TCGA clinicalMatrix) grade, lymphovascular invasion, perineural invasion, extranodal extension and positive margin (Mann-Whitney / Spearman / Kruskal-Wallis).**

Supplementary Data 42 (sheet SD42\_clinical\_associations). Association of each score with pathological N and T stage, HPV status, site, and (legacy TCGA clinicalMatrix) grade, lymphovascular invasion, perineural invasion, extranodal extension and positive margin (Mann-Whitney / Spearman / Kruskal-Wallis).

Supplementary Data 42b (sheet SD42b\_external\_cohorts\_cox). External cohorts (GEO GSE65858 Illumina HT-12, GSE41613 Affymetrix U133 Plus 2): classifier projected with within-cohort z-scoring. HR per SD for every score, univariable and adjusted for stage, age, site and HPV16 DNA or RNA in GSE65858 and for stage and age band in GSE41613.

Supplementary Data 42c (sheet SD42c\_GSE41613\_scores). [GSE41613] per-sample classifier probabilities, pEMT specificity, score panel and the clinical variables used.

Supplementary Data 42d (sheet SD42d\_GSE65858\_scores). [GSE65858] per-sample classifier probabilities, pEMT specificity, score panel and the clinical variables used.

**Supplementary Data 43. Mechanism-of-action share of reversing L1000 compound signatures for the real query against 50 random gene sets of the same size, with enrichment, empirical p (add-one) and Benjamini-Hochberg q.**

Supplementary Data 43 (sheet SD43\_lincs\_moa\_50\_sets). Mechanism-of-action share of reversing L1000 compound signatures for the real query against 50 random gene sets of the same size, with enrichment, empirical p (add-one) and Benjamini-Hochberg q.

Supplementary Data 43b (sheet SD43b\_lincs\_two\_sided). Per-compound sign test of reversing against mimicking signatures, one-sided as in Supplementary Data 44b and two-sided, each with Benjamini-Hochberg false discovery rate.

**Supplementary Data 44. Mechanism-of-action classes among reversing L1000 signatures, against five random gene sets of the same size (specificity control).**

Supplementary Data 44 (sheet SD44\_lincs\_moa). Mechanism-of-action classes among reversing L1000 signatures, against five random gene sets of the same size (specificity control).

Supplementary Data 44b (sheet SD44b\_lincs\_compounds). SigCom LINCS two-sided enrichment of the classifier signature (top 150 pEMT-high coefficients up, top 150 epithelial-like coefficients down) against the L1000 compound library: per perturbagen, the number of signatures reversing and mimicking the query and a sign test with BH-FDR.

Supplementary Data 44c (sheet SD44c\_lincs\_xpr). Same query against the L1000 CRISPR knockout library.

Supplementary Data 44d (sheet SD44d\_lincs\_shrna). Same query against the L1000 shRNA library.

**Supplementary Data 45. [hnscc] DepMap models: classifier probabilities after within-cohort z-scoring.**

Supplementary Data 45 (sheet SD45\_depmap\_scores\_hnscc). [hnscc] DepMap models: classifier probabilities after within-cohort z-scoring.

Supplementary Data 45b (sheet SD45b\_depmap\_scores\_pan\_squamou). [pan\_squamous] DepMap models: classifier probabilities after within-cohort z-scoring.

**Supplementary Data 46. [hnscc] PRISM secondary screen: Spearman correlation of pEMT specificity with AUC per compound (negative = pEMT-high models more sensitive), BH-FDR, tertile AUC difference and proximity-hit flags.**

Supplementary Data 46 (sheet SD46\_prism\_hnscc). [hnscc] PRISM secondary screen: Spearman correlation of pEMT specificity with AUC per compound (negative = pEMT-high models more sensitive), BH-FDR, tertile AUC difference and proximity-hit flags.

Supplementary Data 46b (sheet SD46b\_prism\_pan\_squamous). [pan\_squamous] PRISM secondary screen: Spearman correlation of pEMT specificity with AUC per compound (negative = pEMT-high models more sensitive), BH-FDR, tertile AUC difference and proximity-hit flags.

Supplementary Data 46c (sheet SD46c\_prism\_set\_test\_hnscc). [hnscc] Shift of proximity-hit compounds towards pEMT-high sensitivity, by the Mann-Whitney test across compounds (anti-conservative, because compounds screened on the same cell lines are not independent) and by the cell-line permutation null, which is the reported test.

Supplementary Data 46d (sheet SD46d\_prism\_set\_test\_pan\_squamo). [pan\_squamous] Shift of proximity-hit compounds towards pEMT-high sensitivity, by the Mann-Whitney test across compounds (anti-conservative, because compounds screened on the same cell lines are not independent) and by the cell-line permutation null, which is the reported test.

**Supplementary Data 47. [hnscc] CRISPR (Chronos) dependency: Spearman correlation of pEMT specificity with gene effect per gene (negative = stronger dependency in pEMT-high models), BH-FDR, and membership of seed, key-protein and drug-target sets.**

Supplementary Data 47 (sheet SD47\_crispr\_hnscc). [hnscc] CRISPR (Chronos) dependency: Spearman correlation of pEMT specificity with gene effect per gene (negative = stronger dependency in pEMT-high models), BH-FDR, and membership of seed, key-protein and drug-target sets.

Supplementary Data 47b (sheet SD47b\_crispr\_pan\_squamous). [pan\_squamous] CRISPR (Chronos) dependency: Spearman correlation of pEMT specificity with gene effect per gene (negative = stronger dependency in pEMT-high models), BH-FDR, and membership of seed, key-protein and drug-target sets.

Supplementary Data 47c (sheet SD47c\_crispr\_sets\_hnscc). [hnscc] Gene-set level CRISPR summary (Mann-Whitney of rho, set vs other genes).

Supplementary Data 47d (sheet SD47d\_crispr\_sets\_pan\_squamous). [pan\_squamous] Gene-set level CRISPR summary (Mann-Whitney of rho, set vs other genes).

**Supplementary Data 48. Every mechanism-of-action class with $\geq 5$ compounds, and the network-predicted compound sets, tested against a permutation null that shuffles pEMT specificity across cell lines and recomputes the class statistic (5,000 permutations).**

Supplementary Data 48 (sheet SD48\_prism\_moa\_permutation). Every mechanism-of-action class with $\geq 5$ compounds, and the network-predicted compound sets, tested against a permutation null that shuffles pEMT specificity across cell lines and recomputes the class statistic (5,000 permutations). Unlike the Mann-Whitney comparison across compounds, this test does not treat compounds screened on the same lines as independent. No class reaches FDR 0.05 in either cohort.

**Supplementary Data 49. [canonical\_pEMT, drugbank targets] Guney network proximity of every drug's targets to the module genes (1,000 degree-matched draws).**

Supplementary Data 49 (sheet SD49\_canonical\_pEMT\_drugbank). [canonical\_pEMT, drugbank targets] Guney network proximity of every drug's targets to the module genes (1,000 degree-matched draws). A drug is a hit (is\_hit) when $z \leq -1.96$ and the normal-approximation FDR $\leq 0.05$, and a seed-stable hit (seed\_stable\_hit) when it is still a hit after removing the single most influential anchor gene. Drug accession, name, type, approval status and ATC level 1 from DrugBank (licensed export of 19 April 2025). DrugBank descriptive text (mechanism of action, ATC titles, categories) is omitted because the licence does not allow redistribution.

Supplementary Data 49b (sheet SD49b\_canonical\_pEMT\_stitch). [canonical\_pEMT, stitch targets] Guney network proximity of every drug's targets to the module genes (1,000 degree-matched draws). A drug is a hit (is\_hit) when $z \leq -1.96$ and the normal-approximation FDR $\leq 0.05$, and a seed-stable hit (seed\_stable\_hit) when it is still a hit after removing the single most influential anchor gene. Drug accession, name, type, approval status and ATC level 1 from DrugBank (licensed export of 19 April 2025). DrugBank descriptive text (mechanism of action, ATC titles, categories) is omitted because the licence does not allow redistribution.

Supplementary Data 49c (sheet SD49c\_classifier\_pEMT\_high\_drug). [classifier\_pEMT\_high, drugbank targets] Guney network proximity of every drug's targets to the module genes (1,000 degree-matched draws). A drug is a hit (is\_hit) when $z \leq -1.96$ and the normal-approximation FDR $\leq 0.05$, and a seed-stable hit (seed\_stable\_hit) when it is still a hit after removing the single most influential anchor gene. Drug accession, name, type, approval status and ATC level 1 from DrugBank (licensed export of 19 April 2025). DrugBank descriptive text (mechanism of action, ATC titles, categories) is omitted because the licence does not allow redistribution.

Supplementary Data 49d (sheet SD49d\_classifier\_pEMT\_high\_stit). [classifier\_pEMT\_high, stitch targets] Guney network proximity of every drug's targets to the module genes (1,000 degree-matched draws). A drug is a hit (is\_hit) when $z \leq -1.96$ and the normal-approximation $FDR \leq 0.05$, and a seed-stable hit (seed\_stable\_hit) when it is still a hit after removing the single most influential anchor gene. Drug accession, name, type, approval status and ATC level 1 from DrugBank (licensed export of 19 April 2025). DrugBank descriptive text (mechanism of action, ATC titles, categories) is omitted because the licence does not allow redistribution.

Supplementary Data 49e (sheet SD49e\_pEMT\_axis\_drugbank). [pEMT\_axis, drugbank targets] Guney network proximity of every drug's targets to the module genes (1,000 degree-matched draws). A drug is a hit (is\_hit) when $z \leq -1.96$ and the normal-approximation $FDR \leq 0.05$, and a seed-stable hit (seed\_stable\_hit) when it is still a hit after removing the single most influential anchor gene. Drug accession, name, type, approval status and ATC level 1 from DrugBank (licensed export of 19 April 2025). DrugBank descriptive text (mechanism of action, ATC titles, categories) is omitted because the licence does not allow redistribution.

Supplementary Data 49f (sheet SD49f\_pEMT\_axis\_stitch). [pEMT\_axis, stitch targets] Guney network proximity of every drug's targets to the module genes (1,000 degree-matched draws). A drug is a hit (is\_hit) when $z \leq -1.96$ and the normal-approximation $FDR \leq 0.05$, and a seed-stable hit (seed\_stable\_hit) when it is still a hit after removing the single most influential anchor gene. Drug accession, name, type, approval status and ATC level 1 from DrugBank (licensed export of 19 April 2025). DrugBank descriptive text (mechanism of action, ATC titles, categories) is omitted because the licence does not allow redistribution.

**Supplementary Data 50. Network proximity seeded on the 52 malignant-arm pEMT genes of the single-cell partition (linear scale), independent of the classifier and containing no EGFR-family gene.**

Supplementary Data 50 (sheet SD50\_malignant\_arm\_proximity). Network proximity seeded on the 52 malignant-arm pEMT genes of the single-cell partition (linear scale), independent of the classifier and containing no EGFR-family gene. Columns give drugs tested, hits, approved hits and hits that survive leaving out any one seed (seed-stable), for DrugBank curated and STITCH targets, in the standard analysis and with each drug's targets that are themselves seeds excluded, with the leading approved hits.

**Supplementary Data 51. Leave-one-dataset-out cross-validation of the classifier for two constructions of the training pseudobulks (the bulk-like sum of linear counts per 10,000, renormalised and log-transformed, as trained, and the mean of the cells' log-normalised**

**profiles), each evaluated with the training-fold scaler and with the held-out fold standardised within itself, as bulk cohorts are scored.**

Supplementary Data 51 (sheet SD51\_pseudobulk\_construction\_cv). Leave-one-dataset-out cross-validation of the classifier for two constructions of the training pseudobulks (the bulk-like sum of linear counts per 10,000, renormalised and log-transformed, as trained, and the mean of the cells' log-normalised profiles), each evaluated with the training-fold scaler and with the held-out fold standardised within itself, as bulk cohorts are scored.

Supplementary Data 51b (sheet SD51b\_pseudobulk\_construction). Classifier refitted on the mean-of-log pseudobulks (same cells): overlap of its top 100 pEMT-high coefficients with the main (summed) classifier, correlation of its pEMT specificity with the main score in TCGA-HNSC and GSE65021, and its AUC for prolonged progression-free survival in GSE65021 beside the main classifier's.

**Supplementary Data 52. Head-to-head logistic models in GSE65021 entering pEMT specificity and one other score together, unadjusted and adjusted, with the correlation between the two scores in this cohort.**

Supplementary Data 52 (sheet SD52\_GSE65021\_headtohead). Head-to-head logistic models in GSE65021 entering pEMT specificity and one other score together, unadjusted and adjusted, with the correlation between the two scores in this cohort.

**Supplementary Data 53. Logistic models in GSE65021 entering one score oriented away from the epithelial state (MLR-EMT or KS) and one epithelial score (76GS or Puram epithelial differentiation) together, unadjusted and adjusted, with the correlation between the two scores in this cohort.**

Supplementary Data 53 (sheet SD53\_GSE65021\_vs\_epithelial). Logistic models in GSE65021 entering one score oriented away from the epithelial state (MLR-EMT or KS) and one epithelial score (76GS or Puram epithelial differentiation) together, unadjusted and adjusted, with the correlation between the two scores in this cohort. MLR-EMT is the faithful implementation. Estimates are maximum likelihood, and the Firth estimates are in Supplementary Data 24d.

**Supplementary Data 54. Pooled hazard ratio per SD under four specifications (univariable and adjusted, each fixed-effect and random-effects), with standard errors taken directly from the Cox fits, not back-calculated from confidence intervals.**

Supplementary Data 54 (sheet SD54\_survival\_sensitivity). Pooled hazard ratio per SD under four specifications (univariable and adjusted, each fixed-effect and random-effects), with standard errors taken directly from the Cox fits, not back-calculated from confidence intervals.

Supplementary Data 54b (sheet SD54b\_sensitivity\_per\_cohort). The per-cohort estimates behind the sensitivity analysis, univariable and adjusted.

Supplementary Data 54c (sheet SD54c\_leave\_one\_cohort\_out). Leave-one-cohort-out random-effects pooling, testing whether either conclusion depends on a single cohort.

Supplementary Data 54d (sheet SD54d\_truncation). Pooling after truncating follow-up in every cohort at 36 and 60 months, since median follow-up differs markedly between cohorts.

Supplementary Data 54e (sheet SD54e\_gse65858\_treatment). GSE65858 stratified by treatment modality (single modality versus multimodal). GSE65858 is the only cohort with mixed treatment and the only one in which pEMT specificity is null. Stratifying does not recover an effect.

**Supplementary Data 55. [canonical\_pEMT] Strict proximity variant: a drug's targets that are themselves seed genes are removed before the distance is computed, so a drug scores only if it lies in the neighbourhood of the programme and not on one of its genes.**

Supplementary Data 55 (sheet SD55\_strict\_proximity\_canonical). [canonical\_pEMT] Strict proximity variant: a drug's targets that are themselves seed genes are removed before the distance is computed, so a drug scores only if it lies in the neighbourhood of the programme and not on one of its genes.

Supplementary Data 55b (sheet SD55b\_strict\_proximity\_classifi). [classifier\_pEMT\_high] Strict proximity variant: a drug's targets that are themselves seed genes are removed before the distance is computed, so a drug scores only if it lies in the neighbourhood of the programme and not on one of its genes.

Supplementary Data 55c (sheet SD55c\_strict\_proximity\_pEMT\_axi). [pEMT\_axis] Strict proximity variant: a drug's targets that are themselves seed genes are removed before the distance is computed, so a drug scores only if it lies in the neighbourhood of the programme and not on one of its genes.
